# Therapy-induced lipid uptake and remodeling underpin ferroptosis hypersensitivity in prostate cancer

**DOI:** 10.1101/2020.01.08.899609

**Authors:** Kaylyn D Tousignant, Anja Rockstroh, Berwyck LJ Poad, Ali Talebi, Reuben RS Young, Atefeh Taherian Fard, Rajesh Gupta, Tuo Zang, Chenwei Wang, Melanie L Lehman, Johan V Swinnen, Stephen J Blanksby, Colleen C Nelson, Martin C Sadowski

## Abstract

**Background:** Metabolic reprograming, non-mutational epigenetic changes, increased cell plasticity and multidrug tolerance are early hallmarks of therapy resistance in cancer. In this temporary, therapy-tolerant state, cancer cells are highly sensitive to ferroptosis, a form of regulated cell death that is caused by oxidative stress through excess levels of iron-dependent peroxidation of polyunsaturated fatty acids (PUFA). However, mechanisms underpinning therapy-induced ferroptosis hypersensitivity remain to be elucidated.

**Methods:** We used quantitative single cell imaging of fluorescent metabolic probes, transcriptomics, proteomics and lipidomics to perform a longitudinal analysis of the adaptive response to androgen receptor-targeted therapies (androgen deprivation and enzalutamide) in prostate cancer (PCa).

**Results:** We discovered that cessation of cell proliferation and a robust reduction in bioenergetic processes were associated with multidrug tolerance and a strong accumulation of lipids. The gain in lipid biomass was fueled by enhanced lipid uptake through cargo non-selective (macropinocytosis, tunneling nanotubes) and cargo-selective mechanisms (lipid transporters), whereas *de novo* lipid synthesis was strongly reduced. Enzalutamide induced extensive lipid remodeling of all major phospholipid classes at the expense of storage lipids, leading to increased desaturation and acyl chain length of membrane lipids. The rise in membrane PUFA levels enhanced membrane fluidity and lipid peroxidation, causing hypersensitivity to glutathione peroxidase (GPX4) inhibition and ferroptosis. Combination treatments against AR and fatty acid desaturation, lipase activities or growth medium supplementation with antioxidants or PUFAs altered GPX4 dependence. Despite multidrug tolerance, PCa cells displayed an enhanced sensitivity to inhibition of lysosomal processing of exogenous lipids, highlighting an increased dependence on lipid uptake in the therapy-tolerant state.

**Conclusions:** Our work provides mechanistic insight into processes of lipid metabolism that underpin the acquisition of therapy-induced GPX4 dependence and ferroptosis hypersensitivity to standard of care therapies in PCa. It demonstrated novel strategies to suppress the therapy-tolerant state that may have potential to delay and combat resistance to androgen receptor-targeted therapies, a currently unmet clinical challenge of advanced PCa. Since enhanced GPX4 dependence is an adaptive phenotype shared by several types of cancer in response to different therapies, our work might have universal implications for our understanding of metabolic events that underpin resistance to cancer therapies.

## Background

Despite significant advancements in detection and treatment over the past decades, prostate cancer (PCa) remains the second most commonly diagnosed cancer among men and the third leading cause of cancer mortality in men worldwide [1]. A major challenge in PCa management stems not from the lack of initial treatment options, but in therapy adaptation leading to resistance. Given the dependence of PCa cells on androgens for growth, proliferation, survival and differentiation, the primary treatment for men with advanced PCa is androgen deprivation therapy (ADT), which is being increasingly supplemented with a new generation of potent androgen synthesis inhibitors (Abiraterone) or androgen receptor (AR) antagonists Bicalutamide, Enzalutamide and Apalutamide [2, 3], hereafter cumulatively termed androgen targeted therapies (ATTs). Despite initial treatment response, unfortunately ATTs will ultimately fail in most PCa patients, and disease progresses to castrate-resistant PCa (CRPC) [3, 4] in which PCa cells have often adopted a number of complex resistance mechanisms, including AR amplification, mutation, or splice variants to maintain AR function in the low androgen environment [3, 5]. Other resistance mechanisms bypass AR by exploiting alternative signaling and metabolic pathways. Understanding the early underlying molecular mechanisms that ultimately result in resistance to ATTs is critical for the discovery and development of co-treatment strategies and the definition of therapeutic windows to extend the efficacy of ATTs and combat therapy resistance.

While previous studies in PCa provided an extensive characterization of PCa models with established ATT resistance [6–8], therapy-induced early metabolic reprogramming that underpins therapy survival and development of drug resistance has remained largely unexplored. Recent work in breast, melanoma, lung and ovarian cancer showed that different targeted anti-cancer therapies induced a transient quiescent state of negligible growth termed drug-induced persister cells [9]. Persister cells are characterized by increased expression of markers of dedifferentiation, enhanced lipid peroxidation, an acquired dependency on lipid hydroperoxidase GPX4, and sensitivity to ferroptosis that offer novel therapeutic strategies to combat resistance [9]. Whether ATTs cause PCa cells to acquire the persister cell phenotype remains to be shown.

Ferroptosis is a form of regulated cell death characterized by iron-independent lipid peroxidation of polyunsaturated fatty acids (PUFAs) that was recently identified as a metabolic vulnerability in several types of cancer, including lymphoma, renal cell carcinoma and breast cancer [10, 11]. In particular, reactive oxygen species-mediated peroxidation of PUFAs arachidonic acid and adrenic acid within the phospholipid class of phosphatidylethanolamines are potent activators of ferroptosis [12]. Glutathione peroxidase 4 (GPX4), a selenocysteine enzyme that utilizes glutathione to reduce toxic lipid peroxides into non-toxic lipid alcohols, is critical for the protection against lipid peroxide-mediated damage and ferroptosis induction [11]. Recently, a second, glutathione-independent protective pathway containing ferroptosis-suppressor-protein-1 (FSP1/AIFM2) was found to act in parallel to GPX4 [13, 14]. FSP1 functions as an oxidoreductase with coenzyme Q10 at the plasma membrane to generate lipophilic radical-trapping antioxidants that halt lipid peroxidation [13]. While ferroptosis is an attractive and highly promising concept to combat resistance to multiple therapies in various types of cancer, the metabolic changes that lead to enhanced ferroptosis sensitivity in the context of therapy resistance remain uncharacterized [9].

Dysregulation of lipid homeostasis is a metabolic hallmark associated with PCa tumorigenesis and disease progression to CRPC in response to ATTs. Yet, therapy-induced metabolic reprograming and lipid supply mechanisms other than *de novo* lipid synthesis have not been studied in this context. Previous data from our longitudinal xenograft model of progression to CRPC demonstrated alterations in lipid metabolism during the development to CRPC [15], including increased intratumoral levels of essential PUFAs derived from dietary sources implicating lipid uptake. Emerging evidence reveals lipid scavenging routes utilized by cancer cells during stress in cell culture and pre-clinical models, including lipid transporters [16], the formation of tunneling nanotubes [17], serum albumin-lipid conjugate uptake and macropinocytosis of necrotic cell debris from the microenvironment [18]. Many of these lipid supply pathways converge in lysosomes, where degradation of endocytic vesicles releases lipids and other nutrients intracellularly [19], highlighting the importance of lysosomal activity for lipid supply.

In the present study, we delineated the longitudinal metabolic changes in PCa cells in response to ATTs that lead to therapy-induced persister cells characterized by increased lipid content and uptake and substantial lipid remodeling, multidrug tolerance and ferroptosis hypersensitivity. We show that acquisition of this therapy-induced phenotype can be efficiently suppressed by several combination treatments directed against lipid processing enzymes. In addition, this work identified enhanced dependence of persister cell on lysosomal processing of exogenous cholesterol. Together, our work gained novel mechanistic insight into therapy-induced ferroptosis hypersensitivity and provided multiple strategies that have the potential to extend the efficacy of ATTs, delay resistance development and disease progression to lethal CRPC.

## Methods

### Cell culture

LNCaP (ATCC, CVCL_0395) and C4-2B (ATCC, CVCL_4784) cells were cultured in RPMI medium (Thermo Fisher) supplemented with 5% FBS. Medium was changed every 3 days and cells were incubated at 37°C in 5% CO2. For treatments representing androgen-deprivation therapy, cells were cultured in RPMI medium supplemented with 5% charcoal stripped-serum (CSS). For AR targeted therapies, cells were cultured in 5% or 10% FBS with AR-antagonist Enzalutamide (Enz, 10 µM). Cells were passaged at approximately 80% confluency by trypsinization. Cell lines were genotyped in March 2018 by Genomics Research Centre (Brisbane) and routinely tested for mycoplasma infection.

### RNA extraction and quantitative real-time polymerase chain reaction (qRT-PCR)

48 hours after seeding, cells were treated with the indicated inhibitors. After an additional 48 hr, RNA was harvested using the RNEasy mini kit (Qiagen) following the manufacturer’s instructions. Before extraction, RNA was treated with DNase (Qiagen) to improve RNA purity. Concentration of RNA was measured using a NanoDrop ND-1000 Spectrophotometer (ThermoScientific) and cDNA was prepared from 2 ug total RNA with Superscript III (Invitrogen). qRT-PCR was performed with SYBR Green PCR Master Mix (Invitrogen) on a ViiA-7 Real-Time PCR system (Applied Biosystems). Determination of relative mRNA levels was calculated using the comparative 2-CT method [20] compared to the expression of housekeeping gene receptor like protein 32 (*RPL32*) in each treatment and calculated as fold change relative to vehicle control (DMSO). All experiments were performed in triplicate and analysis and statistics were performed with GraphPad Prism software. Primer sequences can be found in supplementary information (Table S5).

### Microarray gene expression profiling on the 180k VPC custom arrays

For gene expression profiling, triplicates of each sample were analyzed on a custom 180k Agilent oligo microarray (VPCv3, ID032034, GEO: GPL16604). This array contains probes mapping to human protein-coding as well as non-coding loci; with probes targeting exons, 3’UTRs, 5’UTRs, intronic and intergenic regions. RNA was isolated using the RNeasy Mini Kit (Qiagen, Hilden, Germany) according to the manufacturer’s protocol including an on-column DNAse treatment step. RNA purity and quality were evaluated on a NanoDrop1000 (Thermo Fisher Scientific Inc, Waltham, USA) and Agilent 2100 Bioanalyzer (Agilent Technologies, Santa Clara, USA). 150 ng RNA of each sample were amplified and labelled using the Agilent ‘Low Input Quick Amp Labeling Kit’ for One-Color Microarray-Based Gene Expression Analysis. In brief, the input RNA is reverse transcribed into cDNA, using an oligo-dT/T7-promoter hybrid primer which introduces a T7 promoter region into the newly synthesized cDNA. The subsequent in vitro transcription uses a T7 RNA polymerase, which simultaneously amplifies the target material and incorporates cyanine 3-labeled CTP. cDNA synthesis and in vitro transcription were performed at 40°C for 2 h, respectively. The labelled cRNA was purified using the Qiagen RNeasy Mini Kit and quantified on a NanoDrop1000. Finally, 1650 ng cRNA of each sample were hybridized at 65°C for 17 h and the arrays subsequently scanned on an Agilent C-type Microarray Scanner G2565CA.

### Microarray data analysis

The microarray raw data were processed using the Agilent Feature Extraction Software (v10.7). A quantile between array normalization was applied and differential expression was determined using the Baysian adjusted t-statistic linear model of the ‘Linear Models for Microarray Data’ (LIMMA) [21] package in R. The p-values were corrected for a false discovery rate (Benjamini & Hochberg 1995) of 5% and the gene expression levels are presented as log2 transformed intensity values. Normalized gene expression data have been deposited in NCBI’s Gene Expression Omnibus (GEO) and are accessible through GEO Series accession number GSExxx. Probes significantly different between two groups were identified with an adjusted p-value of ≤0.05, and an average absolute fold change of ≥1.5. For functional annotation and gene network analysis, filtered gene lists were examined using QIAGEN’s Ingenuity® Pathway Analysis (IPA®, QIAGEN, Redwood City, www.qiagen.com/ingenuity) and ‘Gene Set Variation Analysis’ (GSVA) [22], ‘Gene Set Enrichment Analysis’ (GSEA) [23], ‘Gene Ontology enRIchment anaLysis and visuaLizAtion tool’ (GOrilla) [24], and GOsummaries [25].

### Comparative gene signature scoring

Gene sets of indicated signatures were acquired from Kyoto Encyclopedia of Genes and Genomes (KEGG), Gene Ontology, Ingenuity Pathway Analysis, REACTOME and the Molecular Signature Database (hallmark gene sets, Broad Institute). GEO deposited RNAseq data sets GSE104935 [26], GSE88752 [27] and GSE48403 [28] were downloaded as raw counts and processed by an edgeR pipeline with TMM normalization to obtain fragments per kilobase of transcript (fpkm) values. Mean expression was used to collapse multiple isoforms. Microarray data of this study were processed through limma pipeline, and Ensembl v77 probes were collapsed to gene level using mean log2 intensities. GSVA [22] was used for signature scoring, and non-scaled bubble plots were created with Morpheus webtool [29], with color indicating the direction of change of the GSVA scores (red=increased scores/gene sets increase in overall expression, blue=decreased scores/gene sets decrease in overall expression).

### Quantitative single cell analysis (qSCI) of lipid content by fluorescent microscopy

Prior to seeding, 96-well Ibidi optical plates were coated with 150 µl Poly-l-ornithine (Sigma) and washed with PBS to increase cell attachment. PCa cell lines pre-treated with either 5% FBS+Enzalutamide (10 µM) or 5% CSS were harvested three days prior completion of the indicated treatment times by trypsinization and seeded into PLO-coated 96-well Ibidi optical plates at a density of 6,000 cells/well in their corresponding types of media (RPMI medium supplemented with either 5% FBS (D0), 5%FBS+Enzalutamide (10 µM) or 5% CSS). After 3 days, media was removed, and cells were washed with PBS and fixed with 4% paraformaldehyde (PFA). Lipids were stained with 0.25 µg/ml Nile Red (Sigma) overnight as described previously [30], and Nuclear DNA was counterstained with 1 µg/ml 4’,6-diamidino-2-phenylindole (DAPI, Thermo Fisher). Alternatively, free cholesterol was stained with 50 µg/ml Filipin (Sigma), and DNA was counterstained with Draq5 (1 µM, Thermo Fisher). >500 cells/well were imaged using InCell 2200 automated fluorescence microscope system (GE Healthcare Life Sciences). Quantitative analysis was performed with Cell Profiler Software [31] to measure total cellular content of neutral lipids, phospholipids and free cholesterol based on mean fluorescence intensity and morphometric analysis of lipid droplets.

### qSCI of lipid transporter expression by immunofluorescence microscopy

72 hours after seeding of ATT pre-treated LNCaP cells in PLO-coated 96-well Ibidi optical plates and PFA fixation as described above, cells were permeabilized with TBS-0.1% Triton X-100 in PBS for 5 min followed a 10 min treatment with TBS-2% BSA to block non-specific binding. Primary antibodies (LDLR (Abcam, ab52818) and SCARB1 (Abcam, ab217318)) were added at a 1:100 dilution in TBS-2% BSA (60 μL/well). After 24 hours, primary antibody was removed and cells were washed with TBS-0.1% Triton X-100 3 times for 5 minutes each. Secondary antibody (Alexa Fluor 568, Thermo Fisher) was added to cells at a 1:250 dilution in TBS-2% BSA (60 μL /well). After one hour in the dark, secondary antibody was removed and cells were washed with TBS-0.1% Triton X-100 3 times for 5 minutes each. DNA and F-actin were counterstained for 20 min in the dark with 1 μL/ml 4’,6-diamidino-2-phenylindole (DAPI, Invitrogen) and Alexa Fluor 647 phalloidin (Thermo Fisher), respectively. Staining solution was replaced with 200 μL PBS. Imaging and quantitative analysis were carried out as described above using the InCell 2200 System and CellProfiler software.

### qSCI of metabolic processes

ATT pre-treated cells were seeded as described above. For quantifying C16:0 fatty acid uptake, growth media was exchanged with 65 µl/well of serum-free RPMI media (Thermo Fisher) supplemented with 0.2% BSA (lipid-free, Sigma) and C16:0-Bodipy (5 µM, Thermo Fisher) and Mitotracker Orange CMTMRos (0.4 µM, Thermo Fisher) and incubated at 37°C for one hour. For quantifying cholesterol uptake, media was exchanged with 65 µl/well of serum-free RPMI media (0.2% lipid-free BSA) supplemented with 15 µM NBD (22-(N-(7-Nitrobenz-2-Oxa-1,3-Diazol-4-yl) Amino-23,24-Bisnor-5-Cholen-3β-Ol, Thermo Fisher) and incubated at 37°C for 2 hours. For quantifying lipoprotein complex uptake, media was exchanged with 65 µl/well of serum-free RPMI media (0.2% lipid-free BSA) supplemented with 1,1’-Dioctadecyl-3,3,3’,3’-Tetramethylindocarbocyanine (DiI)-labelled acetylated-LDL (15µg/ml, Thermo Fischer) or DiI-labelled LDL (15 µg/ml, Thermo Fisher) and incubated at 37°C for 2 hours. Phosphatidylethanolamine uptake was measured as described above in serum-free RPMI media (0.2% lipid-free BSA) or, where noted, in conditioned media by incubation of cells at 37°C for 1 hour with NBD-PE (5 µM, 22-(N-(7-Nitrobenz-2-Oxa-1,3-Diazol-4-yl)-1, 2-Dihexadecanoyl-*sn*-Glycero-3-Phosphatidylethanolamine) triethylammonium salt, Thermo Fisher). After incubation, cells were fixed with 4% PFA. Cellular DNA and F-actin was then counterstained with DAPI and Alexa Fluor 647 Phalloidin (Thermo Fisher). Image acquisition and quantitative analysis were performed as described above.

For measuring cellular uptake of lyso-PC-A2, cells were seeded and processed as described above and incubated for 60 min at 37°C with 65 µl/well of conditioned media supplemented with 5 µM Red/Green BODIPY® PC-A2 (A10072, 1-O-(6-BODIPY® 558/568-aminohexyl)-2-BODIPY® FL C5-sn-glycero-3-phosphocholine, Thermo Fisher). This phospholipid derivative carries in close proximity a red and green Bodipy fluorophore on the sn-1 and sn-2 acyl chains, respectively that provides a capacity for dual emission fluorescence ratio detection. Cleavage of the BODIPY® FL pentanoic acid substituent at the sn-2 position by type 2 phospholipases results in decreased quenching by fluorescence resonance energy transfer (FRET) to the BODIPY® 558/568 dye attached to the sn-1 position. The result is a lipase-dependent increase in BODIPY® FL fluorescence emission detected in the range 515-545 nm (green). The FRET-sensitized BODIPY® 558/568 fluorescence signal is expected to show a reciprocal decrease. Notably, dose titration experiments with unconjugated BODIPY® FL pentanoic acid (Thermo Fisher) suggested a passive and slow route of cellular uptake when compared to longer chain FAs (C12:0 and C16:0, data not shown). Following the incubation, cells were fixed with 4% PFA, and cellular DNA and F-actin was then counterstained with DAPI and Alexa Fluor 647 Phalloidin (Thermo Fisher). Image acquisition and quantitative analysis were performed as described above. Lyso-PC-A2 uptake was calculated as ratio of green (cleaved lyso-PC-A2) vs red (uncleaved PC-A2) signal.

Membrane fluidity (differences in membrane lipid packaging) was measured by live ratio-fluoresce microscopy of ATT pre-treated cells after incubation in 80 µl/well of DPBS supplemented with 1 µM di-4-ANEPPDHQ (D36802, Thermo Fisher) for 30 min at 37°C. DNA was counterstained with Hoechst 33342 (1 µg/ml, Sigma-Aldrich) and cells were washed once with DPBS. di-4-ANEPPDHQ labeled cells (n>750 cells, 8 wells/sample, 2 biological replicates) were imaged at 560 nm and 650 nm using the InCell 2200 System, where a spectral red shift is indicative of an increase in the disordered/liquid phase of the plasma membrane. Quantitative image analysis and calculation of the 560nm/650nm ratio was carried out with CellProfiler software.

Lipid peroxidation was measured by live ratiometric fluorescence microscopy of ATT pre-treated cells after incubation in 65 µl/well of serum-free RPMI media (0.2% lipid-free BSA) supplemented with 5 µM with BODIPY 581/591 C11 (Thermo Fisher) for 60 min at 37°C. Peroxidation of the polyunsaturated butadienyl portion of the fluorescent dye results in a spectral fluorescence emission shift from orange (~590 nm) to green (~510 nm).

### Measurement of cell confluence and qSCI of cell death

Cell proliferation as a function of increasing cell confluence was measured by live imaging microscopy with the IncuCyte FLR and Zoom system (Essen BioScience, Ann Arbor, Michigan, USA). Parental or ATT pre-treated PCa cells were seeded as described above in 96-well Essen ImageLockTM plates (Essen BioScience, Ann Arbor, Michigan, USA). Images were acquired at 2 hour intervals with a 10x objective for up to 7 days.

For assessment of cell death, cells were seeded in 96 well plates (n=3 wells/treatment with >4000 cells/wells) as described above and treated with indicated compounds. At the end of the treatment period, cells were co-stained with Hoechst 3342 (1 µg/ml, total cell count) and propidium iodide (5 µg/ml, Sigma-Aldrich, dead cells). Cells were imaged using the InCell 2200 System, the images were analyzed with Cell Profiler software (Broad Institute) for total and dead cell count, and the percentage of dead cells was calculated based on the ratio of dead and total cells.

### Lipidome analysis

All extractions were performed in 2 mL glass vials. Methanol (220 µL) was added to cell pellet of approximately 2 million cells and vortexed. Internal standard (40 µL SPLASH Lipid-o-mix deuterated internal standard obtained from Avanti Polar Lipids, Alabaster, AL, and 20 µL n-Nonadecanoic acid (C19:0)) was added and briefly vortexed. 770 µL MTBE was added and mixture was incubated for 1 h at room temperature in a shaker. Phase separation was induced by adding 200 µL NH_4_CH_3_CO_2_ (150 mM). After vortexing for 20 s, the sample was centrifuged at 2,000 g for 5 min. The upper (organic) phase was collected and stored at −80°C, then diluted into 2:1 MeOH:CHCl_3_ with 7.5 mM NH_4_CH_3_CO_2_ for mass spectrometry analysis. Tandem mass spectrometry (MS) of the intact lipids was performed using a triple quadrupole mass spectrometer (6500 QTRAP, SCIEX, ON, Canada). The lipid extracts (as described above) were diluted 40-fold prior to analysis. Samples were directly infused into the electrospray ionization source of the mass spectrometer using a loop injection method, where 100 µL of sample was loaded into a sample loop using an autosampler and subsequently infused into the mass spectrometer by electrospray ionization at a flow rate of 20 µL/min. Lipid classes were targeted using either precursor ion or neutral loss scans. For quantification, SPLASH was added to cells prior to lipid extraction. Tandem MS data was processed using LipidView (version 1.3beta; SCIEX) using predefined target lists.

Hydrolysis and derivatization of lipids extracts to fatty acid methyl esters (FAME) was performed on-line using trimethylsulfonium hydroxide [32]. FAMEs were analyzed with a gas chromatograph coupled to a mass spectrometer (GC/MS – TQ8040; Shimadzu, Kyoto, Japan). The separation was carried out on a RTX-2330 capillary column (cyanopropyl stationary phase, 60 m x 0.25 mm, film thickness 0.20 μM; Restek, Bellefonte, PA, USA) and the electron ionization energy was set at 70 eV. Conditions for the analysis of FAMEs were as follows: carrier gas, He: column flow at 1 mL/min; 22:1 split ratio, injection volume 1 μL; injector temperature 240 °C; interface and ion source temperature 260 °C. GC oven temperature was maintained at 100 °C for 1 min, thermal gradient 100 °C to 140 °C at 10 °C / min, 140 °C to 175 °C at 6 °C / min, 175 °C to 200 °C at 10 °C / min and hold for 1 min, followed by 200 °C to 250 °C at 5 °C / min and hold for 4 mins. The data were acquired with Q3 scan mode from *m/z* 50 – 650. For data collection the MS spectra were recorded from 4.6 min to 28.33 mins. The data was processed in GC/MS solution software (Shimadzy, Kyoto, Japan). Fatty acids were identified based on retention time alignment with reference FAMEs from a mixture of standards (Food Industry FAME mix (Table S6), Restek, Bellefonte, PA USA). All samples were normalized to internal standard C19:0 and peak areas of relevant FAMEs was subtracted from negative control samples.

### ^13^C carbon tracing of metabolites by mass spectrometry

LNCaP cells were treated with DMSO or Enzalutamide (10 µM) for up to 21 days as described above. 72 hours prior to harvesting, media was replaced with complete growth media supplemented with uniformly labelled ^13^C-acetate (500 µM, Sigma-Aldrich). Metabolites were extracted using 80% methanol in water extraction buffer (Sigma) that was supplemented with uniformly deuterated myristic acid (2 µM, Sigma) as an internal control. Samples were normalized based on total protein measurements in precipitates by bicinchoninic acid assay (BCA, Pierce).

GC/MS analyses were performed using an Agilent 7890A GC equipped with a DB-35MS (30 m - 0.25 mm i.d. - 0.25 μm) capillary column (Agilent Technologies), interfaced with a triple quadruple tandem mass spectrometer (Agilent 7000B, Agilent Technologies) operating under ionization by electron impact at 70 eV. The injection port, interface and ion source temperatures were maintained at 230 °C. Temperature of the quadrupoles was maintained at 150°C. The injection volume was 1 μl, and samples were injected at a 1:25 split ratio. Helium flow was kept constant at 1 ml/min.

GC oven temperature was maintained at 60 °C for 1 minute, increased to 300 °C at 10.0 °C / minute. Post-run temperature was 325 °C, kept for 5 min. The mass spectrometer operated in SIM mode and cholesterol-tms derivative was determined from the fragment at m/z 458,4 (C30H54OSi) as M+0, also m/z fragments representing M+1 up to M+16 were detected in order to determine the ^13^C-Acetate incorporation into cholesterol.

### Statistical Analysis

Statistical analysis were performed with Graphpad Prism 8.3 (Graphpad Software, San Diego, CA) and R Studio (RStudio, Boston, MA). Data reported and statistical tests are described in figure legends.

## Results

### Therapy-induced remodeling of metabolic networks leads to cellular quiescence and multidrug tolerance

Our previous longitudinal analysis of LNCaP tumor xenografts in a model of CRPC progression following castration showed that CRPC tumors contained significantly higher lipid levels than tumors resected from sham-castrated mice [33]. These lipids included essential fatty acid (FA) derivatives, indicating that development of CRPC is associated with enhanced uptake of exogenous, diet-derived lipids [16]. A detailed review of transcriptomic data from this study by gene set variation analysis (GSVA) revealed that castration caused a negative enrichment of pathways involved in lipogenesis, mitochondrial activity and oxidative phosphorylation in tumors at nadir of serum PSA levels compared to non-castrated tumor-bearing mice (intact, control). In support of therapy-induced changes to tumor lipid supply, tumors at PSA nadir showed increased enrichment in pathways of lipoprotein metabolism, lipid storage, lipid transporter and lysosomal activity (Fig. 1). The complete longitudinal analysis including that from regressing and recurrent tumor samples is shown in Fig. S1A, highlighting the dynamic changes in response to androgen-targeted therapies (ATTs) during progression to CRPC.

**Figure 1:**
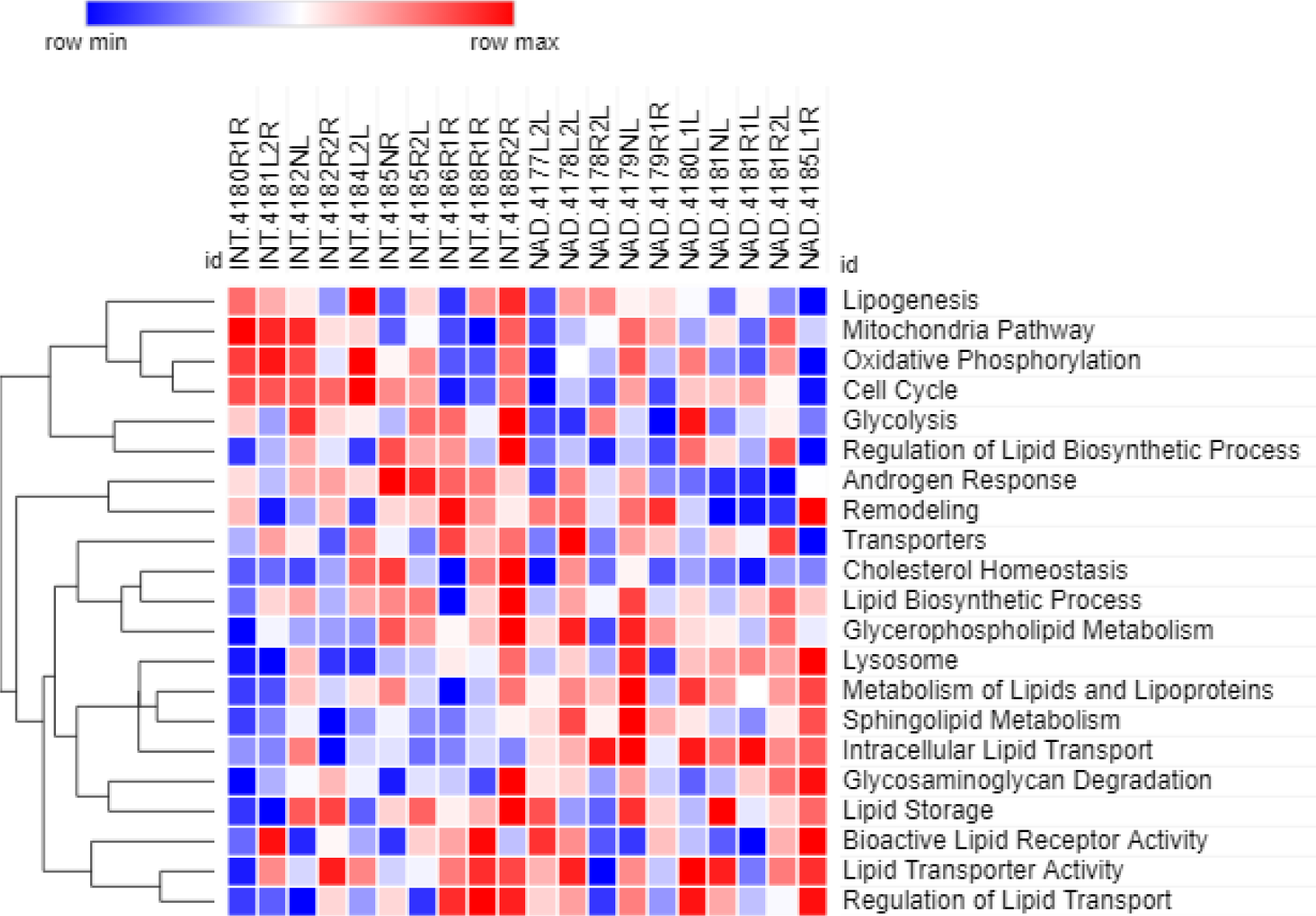
Therapy-induced reprograming of metabolic networks in a longitudinal PCa tumor xenograft model of CRPC progression. (A) Gene set variation analysis (GSVA) of ten tumors at PSA nadir (NAD) compared to ten tumors from sham-castrated mice (INT, intact) was generated based on previously published microarray analysis of LNCaP xenograft tumors resected at different time points after castration [33]. Heatmaps were generated with a hierarchical clustering algorithm using completed linkage and Euclidean distance measures and scaled by z score (red=positive z score, blue=negative z score).

To investigate therapy-induced longitudinal changes to lipid metabolism on the functional and molecular level, the AR-positive, androgen-dependent LNCaP PCa cell line was grown for up to 21 days in the presence of Enzalutamide (Enz) to block AR function. Pairwise sequential comparisons of transcriptome data generated from samples taken prior (day 0) and 7, 14 and 21 days post commencement of Enz treatment showed that LNCaP cells reached transcriptional stasis after 14 days of Enz, i.e., the set of 4524 differentially expressed genes (D0 vs D14) remained unchanged when compared to 21 days Enz (D0 vs D21, Fig. 2A). This suggested that the phenotype induced by Enz was transcriptionally established by day 14 of Enz treatment and persisted for another week. This longitudinal analysis further highlighted that Enz treatment periods beyond prevailing protocols to assess the acute response to ATTs *in vitro* (2-4 days) [16] identified an additional 908 (25%) differentially expressed genes.

**Figure 2:**
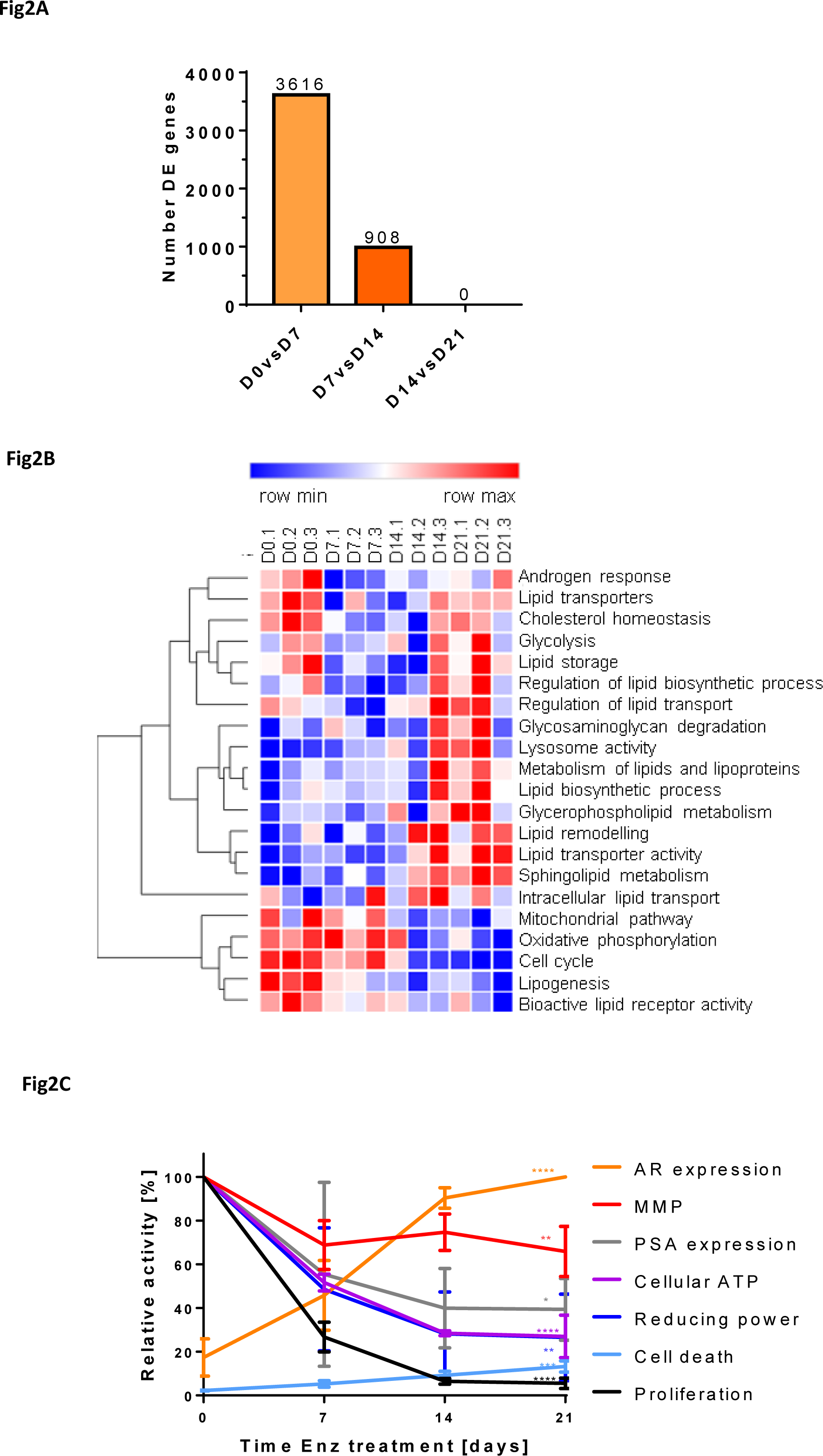
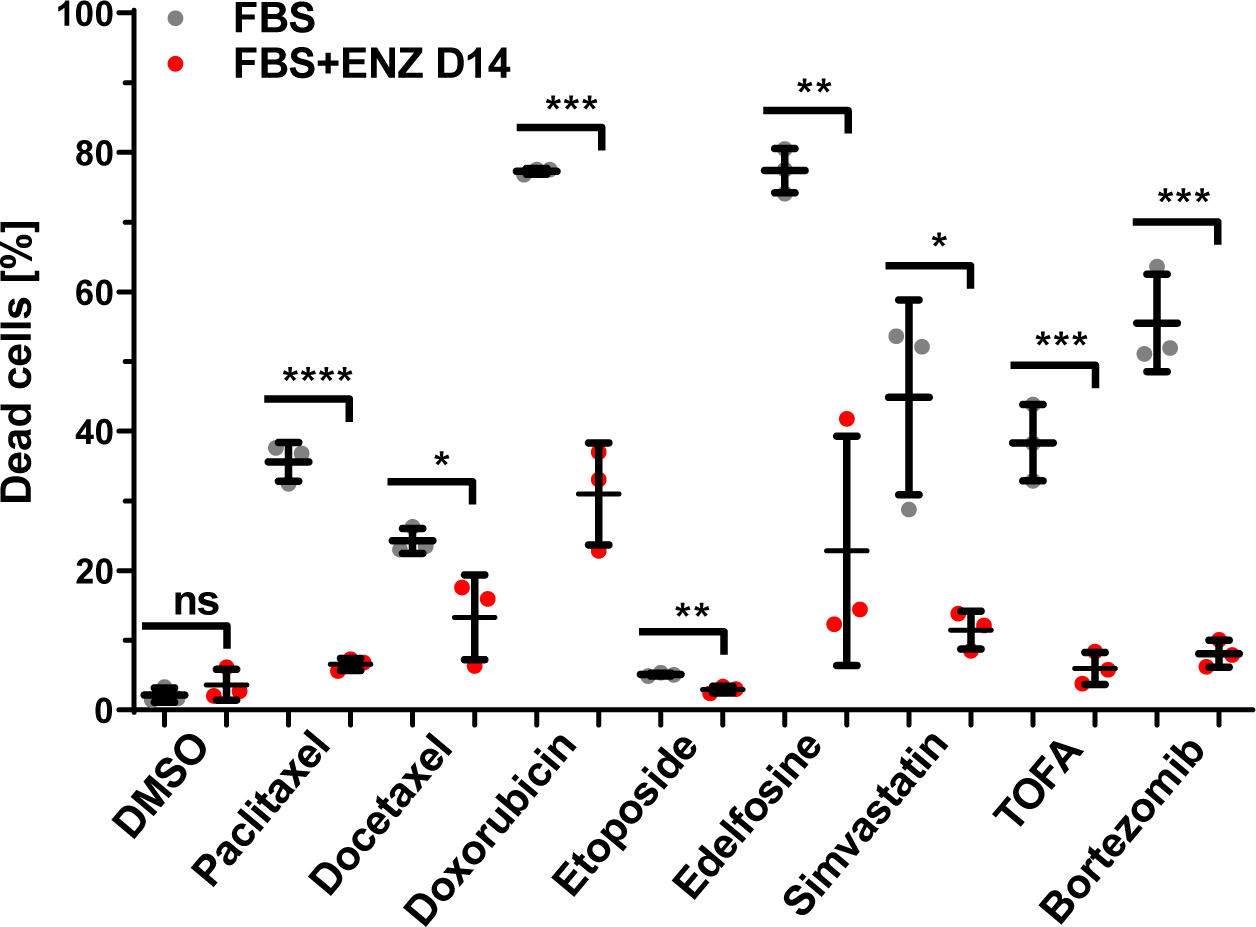
Therapy-induced metabolic reprogramming of LNCaP PCa cells is associated with cellular quiescence and multi drug tolerance. (A) LNCaP cells were treated for 0, 7, 14 and 21 days with AR antagonist Enzalutamide (Enz, 10 µM), and samples were analyzed by microarray analysis. The number of differentially expressed genes between indicated pairwise comparisons are shown (FDR: 1.5fold, p<0.05). (B) Gene set enrichment analysis of the microarray data described in A for the analysis of indicated pathways. (C) LNCaP cells were treated as in A, and samples were analyzed by qRT-PCR (AR and PSA mRNA expression), live-cell confluence imaging (proliferation), CellTiter-Glo assay (cellular ATP levels), PrestoBlue assay (cellular reducing power), quantitative single-cell imaging (qSCI) of MitoTracker Orange CMTMRos (mitochondrial membrane potential, MMP) and dead/live staining (cell death) for functional characterization of Enz-induced metabolic changes (n=3, activities relative to day 0 with the exception of AR levels, where levels were calculated relative to day 21). (D) LNCaP cells were cultured in growth media without (FBS) or with Enz (10 µM, FBS+ENZ D14) for 14 days including the final 24 h in the presence of vehicle control (DMSO) or the indicated compounds (paclitaxel 2.5 µM, docetaxel 2.5 µM, doxorubicin 2.5 µM, etoposide 5 µM, edelfosine 7.5 µM, simvastatin 30 µM, TOFA 60 µM, bortezomib 2.5 µM). The percentage of dead cells was calculated based on qSCI of Hoechst 3342 (total cell count) and propidium iodide (dead cells) staining (n=3 wells/treatment with >4000 cells/well, ns=not significant, *p<0.05 **p<0.01 ***p<0.001 ****p<0.0001, One-way ANOVA followed by Dunnett’s multiple comparisons test compared to FBS, representative result of 2 independent experiments).

GSVA yielded enrichment scores that clustered pathways into three types of adaptive responses to Enz treatment when compared to control (day 0): Pathways that changed to and persisted as either negatively or positively enriched and dynamic pathways that, after temporarily displaying negative enrichment (days 7 and 14), reverted to a positive enrichment seen prior treatment commencement. GSVA revealed negative enrichment for known androgen-activated pathways, e.g. lipogenesis, cell cycle and mitochondrial pathways, including oxidative phosphorylation (Fig. 2B). Pathways with positive enrichment were associated with lysosomal function, lipid metabolism, lipid remodeling and extra- and intracellular lipid transport (Fig. 2B). Dynamic pathways with transient negative enrichment that reverted to their initial positive enrichment seen prior the start of treatment were associated with the androgen response, lipid transporters, cholesterol homeostasis, glycolysis, lipid storage and regulation of lipid biosynthesis and lipid transport (Fig. 2B). GSVA suggested a substantial reconfiguration of the lipid metabolism network in response to Enz over the treatment period of 21 days and were consistent with the *in vivo* results (Fig. 1). Consistent with this, protein mass spectrometry of LNCaP cells treated for 0 and 21 days with Enz and gene ontology analysis revealed enrichment of membrane lipid metabolic processes, glycolipid metabolism and lysosome function (Fig. S1B and Table S2).

Extended cell culture (several months) in androgen-depleted or AR antagonized conditions ultimately yielded androgen-independent or Enz-resistant CRPC cells that reinitiated proliferation [6, 8, 34, 35]. Our comparative signature scoring of transcriptomics data sets of relevant studies performed in cell culture leading to Enz resistance in LNCaP and C4-2B cells [26], in castrated mice progressing to primary and secondary CRPC after serial tumor xenograft transplantation of LAPC9 and LNCaP cells [27] and matching PCa tumors from patients before and after androgen deprivation therapy for 22 weeks [28] by GSVA using lipid metabolism related gene sets showed extensive concordance between cell culture models, pre-clinical tumor xenograft models and clinical samples of ATT resistance (Fig. S1C). This analysis further highlighted the dynamic nature of some therapy-induced metabolic changes, potentially limiting their value as co-treatment options based on the therapeutic window.

The observed time-dependent increase in AR mRNA expression by Enz over 21 days (Fig. 2C, Fig. S2A) was consistent with reported resistance mechanisms to ATT [2, 4]. mRNA levels of *PSA*/*KLK3* remained decreased (Fig. 2C) and a large subset of other AR-activated and suppressed genes (Fig. S2B) did not change, suggesting that AR activity remained suppressed. Consistent with the de-differentiation effect of AR-antagonism, transcriptional levels of several stemness markers were significantly increased (Table S1) [36]. Furthermore, Enz-treated LNCaP cells showed a time-dependent strong reduction in cellular reducing power (−70%), ATP levels (−60%) and mitochondrial membrane potential (−30%), which stabilized after 14 days (Fig. 2C, Fig. S2C). Proliferation almost completely ceased after 14 days of Enz treatment, but cell death was only modestly increased (Fig. 2C, Fig. S2D) which was possibly due to protection via significantly increased autophagy (Fig. S2E). This low cycling phenotype was underpinned by a strong downregulation of genes associated with cell cycle progression (Fig. 2B). Enz-treated LNCaP cells adopted a spindle-like cell morphology marked by a reduced cell size, increased cell perimeter and major axis length (Fig. S2F). Transcriptionally, metabolically and proliferation-wise, LNCaP cells responded to extended Enz treatment by entering into a state of cellular quiescence. Associated with this phenotype was the acquisition of multi-drug tolerance (MDT) to a range of lipogenesis inhibitors and clinically relevant chemotherapeutics for CRPC in LNCaP, C4-2B and DuCaP cells (Fig. 2D and S2G), indicating that this adaptive response to Enz is shared across multiple AR-positive PCa cell lines. Notably, culturing of LNCaP cells in androgen-depleted growth media (CSS) for 14 days showed similar adaptive responses when compared to Enz, e.g., cellular ATP levels and reducing power, proliferation and MDT (Figs. S2C, S2D, and S2G), suggesting that both ATTs cause similar metabolic reprograming that leads to the persister cell state.

### Androgen-targeted therapy increased cellular lipid content

Above results suggested major changes to lipid metabolism networks by Enz. Previous studies, including our own work, have demonstrated that androgens strongly enhance cellular lipid content of PCa cells through enhanced lipogenesis [37, 38] and lipid uptake [16] and that AR antagonists block these lipid supply pathways. To measure Enz-induced changes to the cellular content of neutral lipids, lipid droplets, phospholipids and free cholesterol, we used quantitative single cell imaging (qSCI) of two fluorescent lipophilic dyes, Nile Red and Filipin. Nile Red is able to distinguish between phospholipids and neutral lipids [e.g. triacylglycerols (TAGs) and cholesteryl esters (CEs) stored in lipid droplets] based on their difference in hydrophobicity having separate fluorescence emission maxima [39]. Filipin is a selective fluorescent stain for free, non-esterified cholesterol which is mainly found in the plasma membrane [40]. Contrary to the expected lipid depletion resulting from loss of androgen-enhanced lipid supply, qSCI of Nile Red-stained LNCaP cells demonstrated that Enz significantly increased neutral lipid and phospholipid content with increasing time of treatment (Fig. 3A-C). Morphometric analysis of lipid droplets (LDs) showed that the LD number and average total area of LDs per cell was also significantly increased with treatment time (Fig. 3B left and middle panel). Consistent with this, mRNA expression of the LD surface protein PLIN1 was significantly increased by Enz treatment (Fig. 3B right panel). qSCI of Filipin-stained LNCaP cells showed that Enz induced a significant increase in free cholesterol content (Fig. 3D). Similar observations were made when LNCaP cells were maintained in androgen-depleted media (CSS, data not shown). For molecular analysis of the Enz-induced longitudinal changes to the lipidome, we next ran electrospray ionization mass spectrometry (ESI-MS) of lipidomics of lipid extracts prepared from LNCaP cells treated with vehicle control (FBS) or for 0, 7, 14 and 21 days with Enz. Sparse partial least squares discriminant analysis showed that each time point sample group of the Enz-treatments clustered distinctly from the vehicle treated control group (FBS) and each other (Fig. S3A). LNCaP cells significantly increased cellular levels of sphingomyelin (SM), phosphatidylcholine (PC), phosphatidylethanolamine (PE), phosphatidylserine (PS) and phosphatidylglycerol (PG) over the time course of treatment (Fig. 3E top panel, Fig. S3B-F). While all major membrane lipid classes were increased by Enz, neutral lipids like CEs and TAGs stored within LDs were significantly decreased (Fig. 3E bottom, Figs. S3F and S3G). Of the top 50 deregulated lipid species measured, 86% of these were found to have increased abundance with increasing time of Enz treatment (Fig. S3I). Notably, the majority of these were PC and PE phospholipids that make up ~70% of the lipid content of mammalian cells [41], providing further evidence that Enz increased cellular lipid content. Together, qSCI and lipidomics analyses demonstrated that Enz induced substantial remodeling of all major lipid classes, leading to a net increase in cellular lipid content.

**Figure 3:**
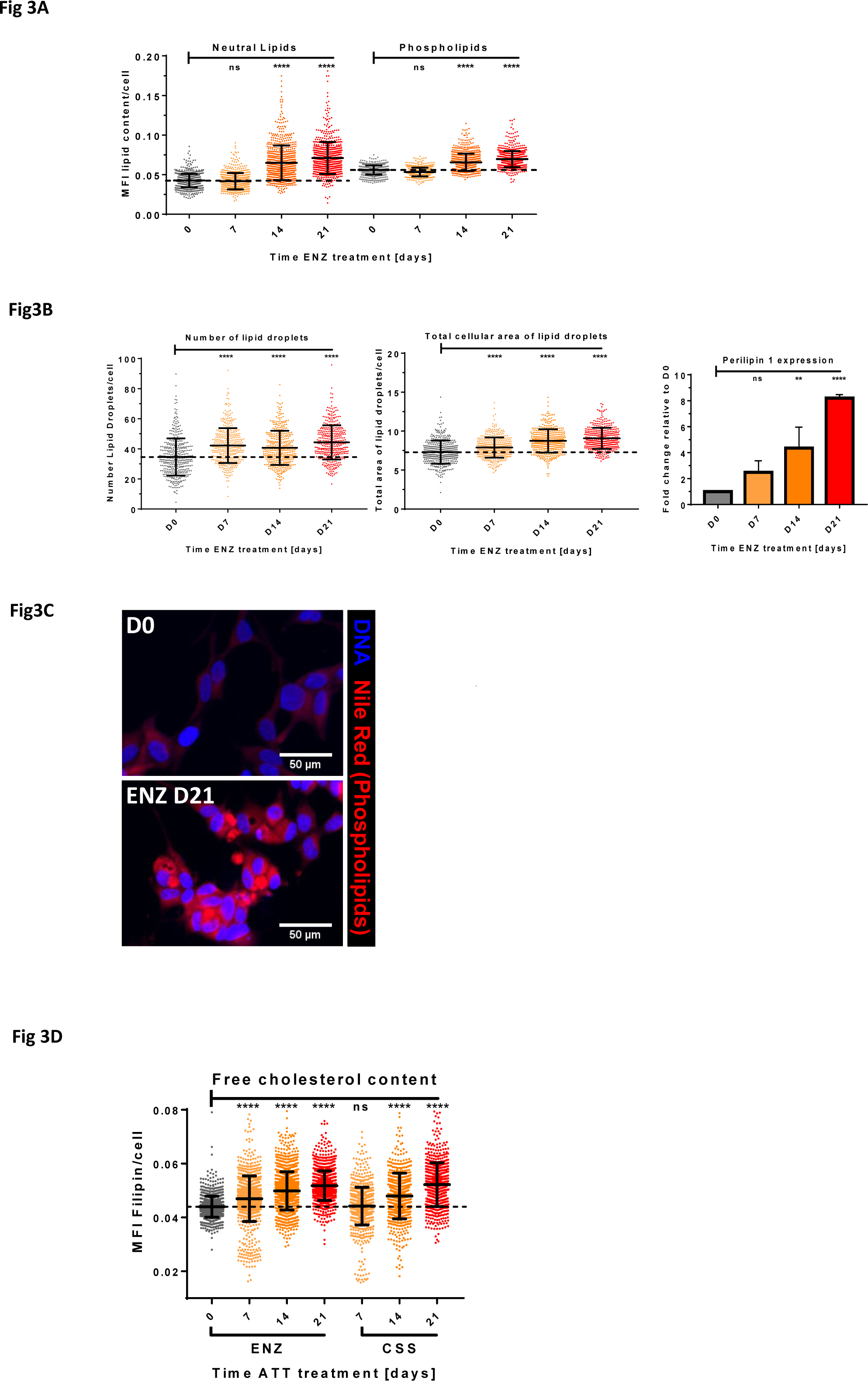
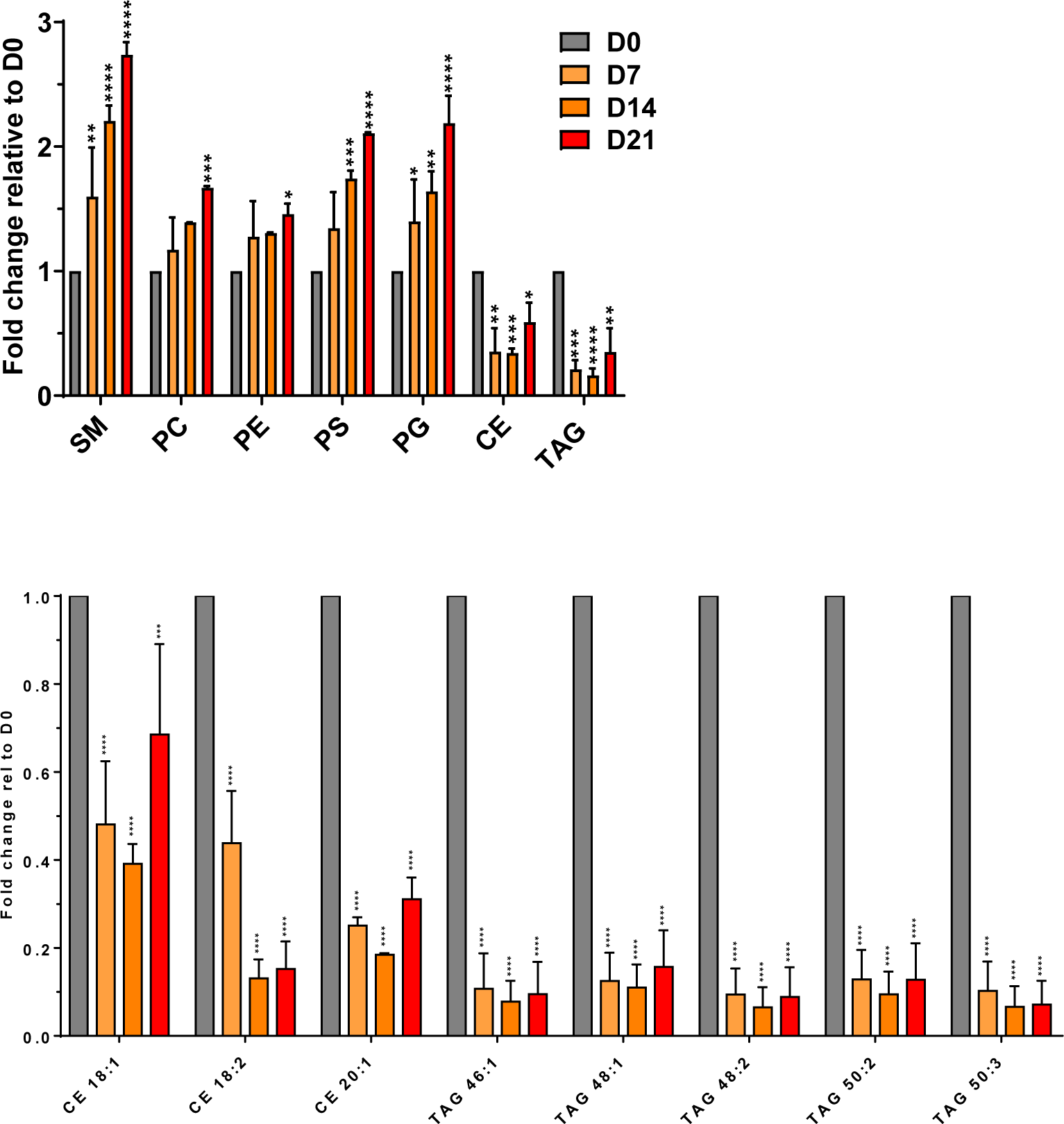
Enzalutamide increased cellular lipid content. LNCaP cells were treated for the indicated times with Enz (10 µM) and analyzed by qSCI based on the mean fluorescence intensity (MFI) of Nile Red to measure (A) cellular neutral lipid and phospholipid content and (B) number of lipid droplets (LDs)/cell and total area of LDs/cell (n>9000 cells/treatment, mean±SD, representative result of 3 independent experiments). Perilipin mRNA expression was measured by qRT-PCR (n=3, mean±SD). (C) Representative images of D0 and D21 showing phospholipid and DNA staining as described in (A). (D) LNCaP cells were treated as in A or grown in androgen-depleted media (CSS, charcoal-stripped serum) for the indicated times, stained with Filipin, and free cholesterol was measured by qSCI (n>9000 cells/treatment, mean±SD, representative result from 2 independent experiments). (E) LC/MS shotgun lipidomics of intact lipid species of LNCaP cells treated as described in A. Total combined sum compositions of indicated lipid classes (top, SM, sphingomyelin; PC, phosphatidylcholine; PE, phosphatidylethanolamine; PS, phosphatidylserine; PG, phosphatidylglycerol; CE, cholesterolester; TAG, triacylglycerol), and individual CE and TAG species (bottom) are reported as fold change relative to D0 (n=2, mean±SD). Statistical analysis: A-D=One-way ANOVA followed by Dunnett’s multiple comparisons test compared to D0; E=Two-way ANOVA followed by Tukey’s multiple comparisons test relative to D0; ns=not significant, *p<0.05 **p<0.01 ***p<0.001 ****p<0.0001

### Enhanced lipid uptake fuels cellular lipid accumulation

As shown above, therapy-induced persister cells displayed multidrug tolerance to lipogenesis inhibitors (Fig. 2D), suggesting that *de novo* synthesis played a limited role in cellular lipid accumulation. Indeed, metabolic tracing of ^13^C-acetate by mass spectrometry revealed that *de novo* cholesterol synthesis decreased by 71% (p>0.0001) after Enz treatment of LNCaP cells for 21 days when compared to control (Fig. 4A). No significant incorporation of ^13^C-glucose, another key carbon source for lipogenesis, into lipids was measured (data not shown), and already low basal glucose uptake was significantly reduced by Enz (Fig. S4A). Consistent with this, transcriptomics and proteomics analysis showed that key enzymes of *de novo* cholesterol and FA synthesis pathways were significantly downregulated by Enz treatment (Fig. 4B, Table S1), suggesting that alternative lipid supply pathways were responsible for therapy-induced lipid accumulation.

**Figure 4:**
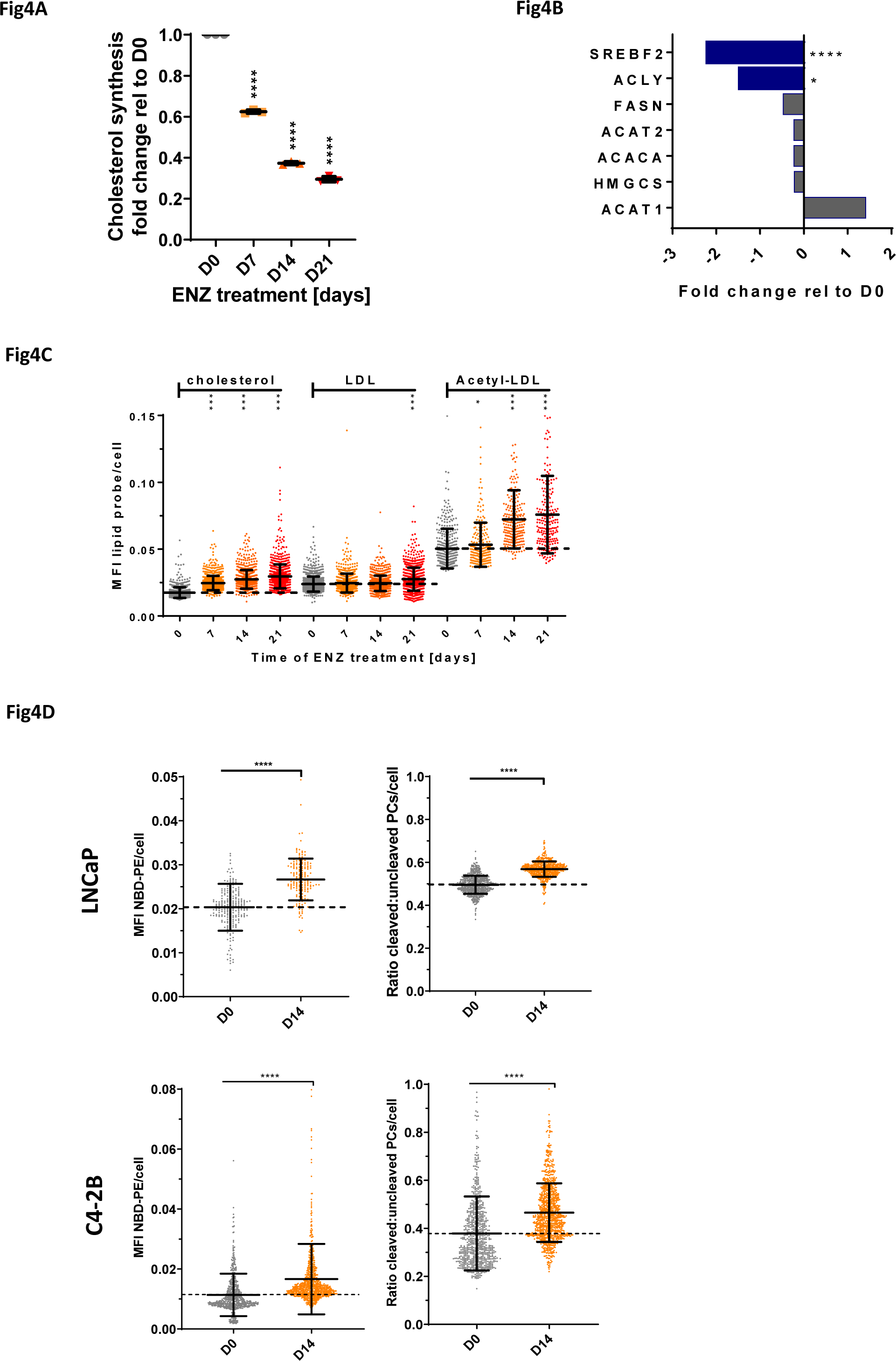
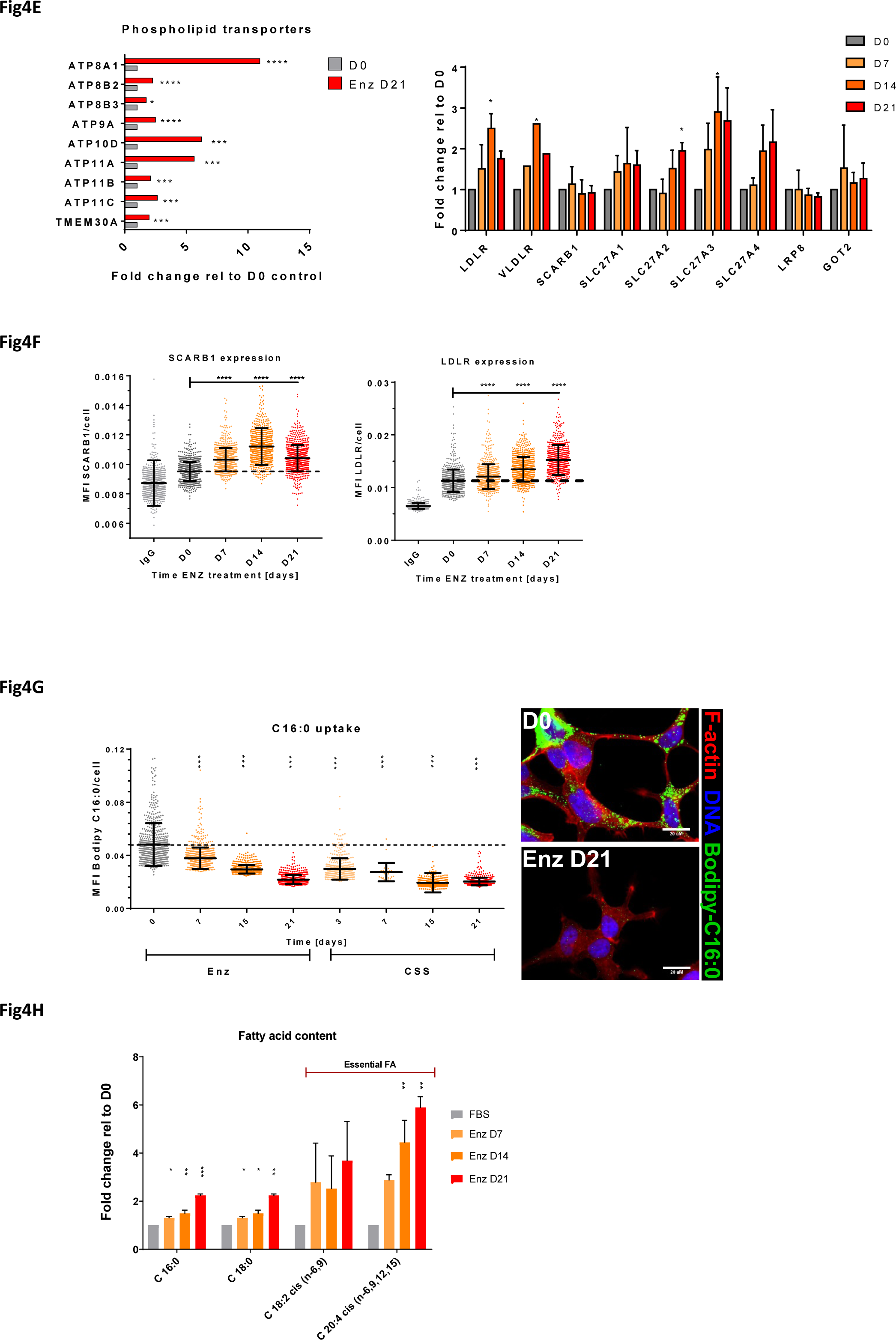
Increased cellular lipid levels are fueled by enhanced lipid uptake. (A) LNCaP cells were treated for the indicated times with Enz (10 µM) including the final 72h in the presence of ^13^C-acetate. Incorporation of ^13^C into cholesterol via lipogenesis was measured by GC/MS (n=3, mean±SD). (B) Protein (blue) and mRNA (gray) expression of indicated enzymes was measured by quantitative proteomics with isobaric mass tag labels and microarray analysis, respectively (n=3, mean±SD). (C) LNCaP cells were treated as described in A, and cellular uptake of indicated fluorescent lipid probes was measured by qSCI based on the mean fluorescence intensity (MFI) of NBD-cholesterol, LDL(DiL), acetylated-LDL(DiL) or (D) NBD-PE (left panel) and PC-A2 (right panel) in LNCaP and C4-2B cells (n>9000 cells, mean±SD, representative of 2 independent experiments). (E) Fold-change mRNA expression changes of indicated lipid transporters relative to D0 of Enz-treatment were calculated from the microarray data described in Figure 2A (left panel) or measured by qRT-PCR (right panel). (F) LNCaP cells were treated for the indicated times with Enz (10 µM), and protein expression of lipid transporters SCARB1 (left) and LDLR (right) was measured using immunofluorescence microscopy (IgG=antibody control, n>9000 cells, mean±SD, representative result from 2 independent experiments). (G) LNCaP cells were treated with Enz (10 µM) or grown in androgen-depleted media (CSS, charcoal-stripped serum) for the indicated times, and FA uptake was measured by qSCI based on the MFI of cellular Bodipy-C16:0 (n>9000 cells, mean±SD, representative result from 3 independent experiments). (H) LNCaP cells were treated for the indicated times with Enz (10 µM), and fatty acid content was analyzed by GC/MS FAME (n=2, mean±SD). Statistical analysis: A-H=One-way ANOVA followed by Dunnett’s multiple comparisons test compared to D0; *p<0.05 **p<0.01 ***p<0.001 ****p<0.0001

To measure cellular lipid uptake, qSCI with various fluorophore labeled lipid probes (NBD-cholesterol, DiI-LDL, DiI-acetylated LDL, Bodipy-C16:0, NBD-phosphatidylethanolamine, Bodipy-phosphatidylcholine) were conducted. Enz treatment significantly increased cellular uptake of free NBD-cholesterol and DiI-complexed low density lipoprotein (LDL) and acetylated LDL in LNCaP cells, as well as NBD-phosphatidylethanolamine and Bodipy-lyso-phosphatidylcholine in LNCaP and C4-2B cells (Fig. 4C-D) in a time-dependent manner. Consistent with increased lipid uptake and cellular content, Enz significantly increased the mRNA levels of several lipid transporters, including phospholipid transporters and co-factors belonging to the family of P4-ATPases/flippases (Fig. 4E, left panel; Table S1) and other cargo-selective lipid transporters, including low-density lipoprotein receptor (LDLR) (Fig. 4E right panel). Protein expression of several of these lipid transporters was upregulated by several orders of magnitude in clinical samples of bone metastatic CRPC when compared to localized, hormone naïve PCa and normal prostate gland [42] (Fig. S4B), suggesting that enhanced lipid uptake might remain activated throughout the development of ATT resistance and CRPC. LDL particles are a bulk supply of apolipoproteins and complex lipids, including CEs, TAGs, and phospholipids to cells via receptor-mediated endocytosis of LDLR and scavenger receptor SCARB1 [43, 44]. Consistent with increased cellular uptake of LDL and acetylated LDL, protein expression of lipid transporters LDLR and SCARB1 was significantly enhanced by Enz. (Fig. 4F). Surprisingly, uptake of saturated free FA (Bodipy-C16:0) was drastically decreased by both androgen-depleted serum (CSS) and Enz treatment over time (Fig. 4G), suggesting that ATT-induced changes to transporter-mediated lipid uptake are specific. Gas chromatography-mass spectrometry (GC/MS) analysis of derivatized FAs released upon saponification of complex lipids revealed that saturated FAs (C16:0 and C18:0) and essential PUFAs linoleic acid (C18:2n-6) and its derivative arachidonic acid (C20:4n-6) were significantly increased by Enz with increasing time of treatment (Figs. 4H and S4C). Together, above results suggested that saturated FA requirements from exogenous sources were preferably met by uptake of complex lipids (e.g., TAGs, CEs, phospholipids) which are also rich sources of PUFAs (Fig. S4D). Despite this observed cargo selectivity, Enz also enhanced cargo non-selective mechanisms involved in lipid scavenging, e.g. macropinocytosis [45] and tunneling nanotubes (TNTs, Figs. S4E and S4F) [46]. Together, these results demonstrated that Enz-treated PCa cells overall enhanced lipid uptake through cargo-selective and non-selective mechanisms.

### Therapy-induced lipid remodeling increases membrane fatty acid desaturation and fluidity

Above results showed that Enz affected all major lipid classes and strongly increased cellular content of phospholipids, suggesting changes to membrane composition and function. Exemplified by lipid species of the phosphatidylethanolamine (PE) phospholipid class, PEs with shorter FA chains (34:2 and 36:2) were significantly decreased, while PEs with longer and more unsaturated FA chains (38:3, 38:4, 38:5 and 38:6) were significantly increased by Enz treatment compared to control (Fig. 5A). Similar trends were seen in SM, PC, PE, PS, and PG lipid classes (Fig. S3B-F), where an enrichment of phospholipids with long-chain PUFAs was observed. Integrated analyses of all dysregulated fatty acid species detected by GC/MS FAME analysis (Fig. S4C) or proteins detected by proteomics (Fig. S1B) with the mRNA expression changes by microarray (Fig. 2) of LNCaP cell treated with vehicle control or Enz for 21 days demonstrated significant enrichment of multiple pathways involved in PUFA metabolism (Fig. S5A), including those of essential PUFAs (e.g. linoleic acid and arachidonic acid).

**Figure 5:**
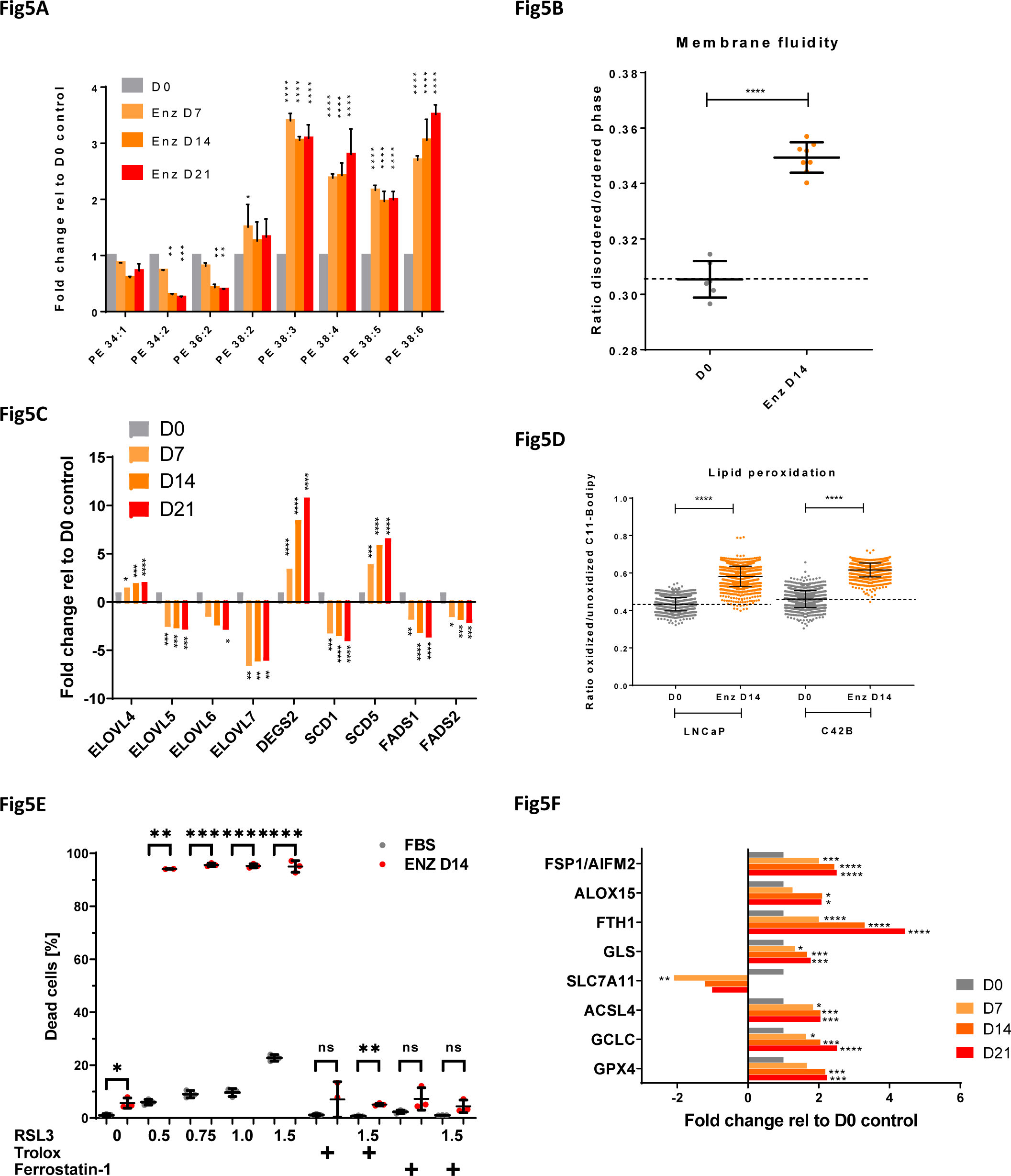
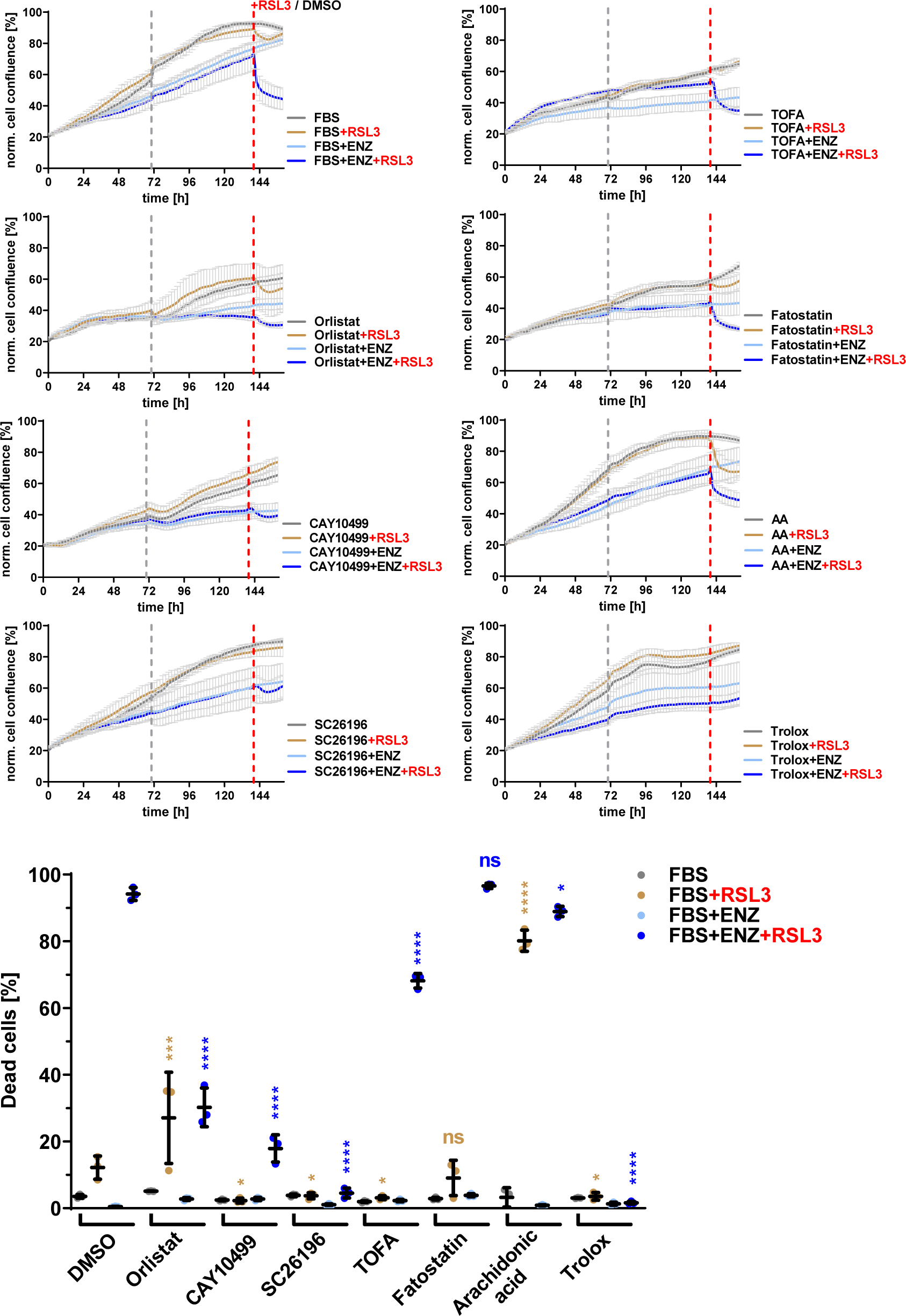
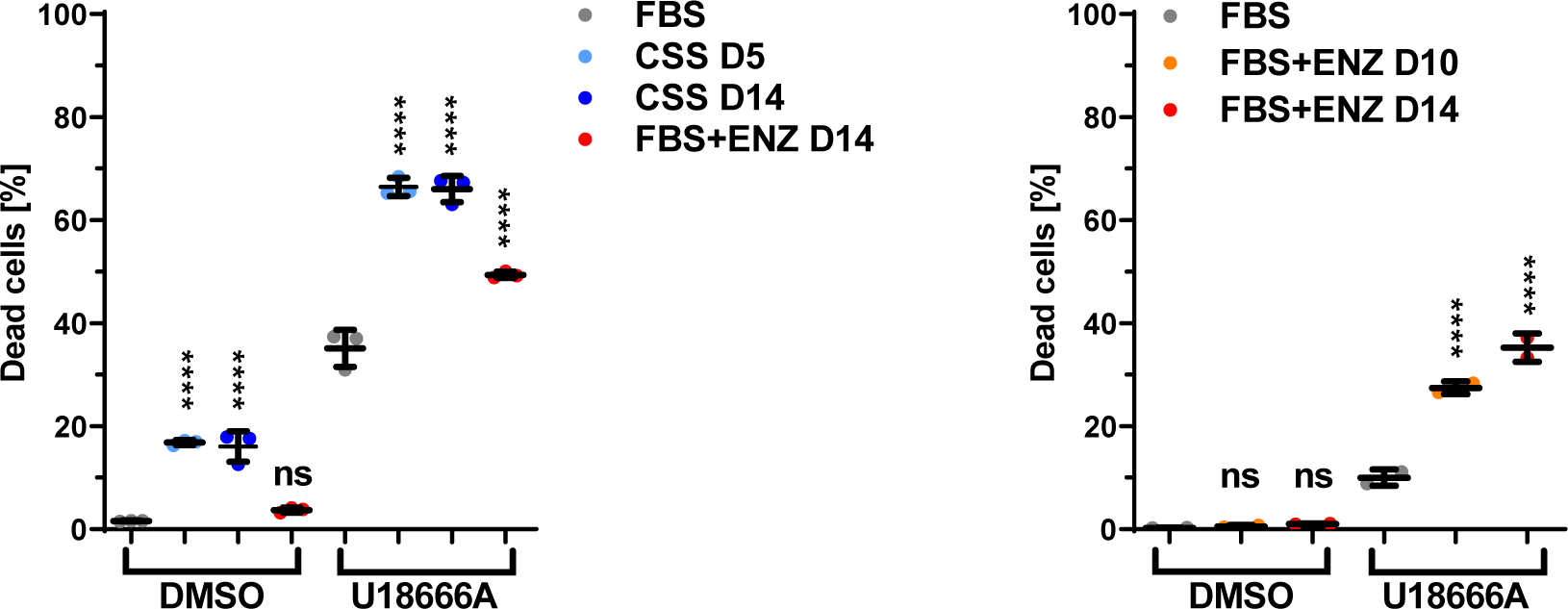
Therapy-induced lipid remodeling leads to increased membrane PUFA levels and lipid peroxidation and underpins GPX4 dependence. (A) LNCaP cells were treated for the indicated times with Enz (10 µM), and lipids were analyzed by LC/MS. Individual sum compositions of phosphatidylethanolamine are shown (n=2, mean±SD). (B) LNCaP cells were treated as in A, and membrane fluidity was measured by qSCI based on the mean fluorescence intensity (MFI) of Di-4-ANEPPDHQ (n=8 wells, ~750 cells/well, mean±SD, representative results from 2 independent experiments). (C) Relative mRNA expression changes of indicated lipid remodeling enzymes (elongases and desaturases) were calculated from the microarray data described in Figure 2A. (D) LNCaP cells were treated for the indicated times with Enz (10 µM), and lipid peroxidation was measured by qSCI based on the mean fluorescence intensity (MFI) of Bodipy-C11 (n>9000 cells, mean±SD, representative results from 2 independent experiments). (E) LNCaP cells were cultured in growth media without (FBS) or with Enz (10 µM, FBS+ENZ D14) for 14 days including the final 24 h in the presence of DMSO (0) or the indicated concentrations (0.5-1.5 µM) of GPX4 inhibitor RSL3. Lipid peroxide scavenger Trolox (100 µM) and ferroptosis inhibitor Ferrostatin-1 (10 µM) were used as controls to demonstrate cell death through ferroptosis. The percentage of dead cells was calculated based on qSCI of Hoechst 3342 (total cell count) and propidium iodide (dead cells) staining (n=3 wells/treatment with >4000 cells/well, representative result of 3 independent experiments). (F) Relative mRNA expression changes of indicated key factors of ferroptosis and glutathione homeostasis were calculated from the microarray data described in Figure 2A. (G) LNCaP cells seeded in a 96 well plate were co-treated with the indicated compounds (orlistat 10 µM, CAY10499 17.5 µM, SC26196 30 µM, TOFA 2.5 µM, fatostatin 5 µM, arachidonic acid (AA) 20 µM, trolox 100 µM) in the absence (FBS) or presence of Enz (10 µM, FBS+ENZ), and cell confluence was measured every 2 hours by live cell imaging with an IncuCyte system (n=3, mean±SD). The gray dotted line indicates the time point of media change. After 7 days of co-treatment, cells were treated with vehicle control (DMSO/FBS) or RSL3 (1.25 µM) to assess GPX4 dependence by monitoring changes to cell confluence for another day. The red dotted line indicates the start of RSL3 treatment. Cell death was then measured by qSCI (bottom graph) as described in (E; n=3 wells/treatment with >4000 cells/well, representative result of 2 independent experiments). (H) LNCaP (left panel) and C4-2B cells (right panel) were cultured in growth media (FBS), androgen-depleted growth media (CSS for 5 and 14 days) or growth media supplemented with Enz (10 µM, FBS+ENZ) for 10 and 14 days including the final 24 h in the presence of vehicle control (DMSO) or U18666A (50 µM). The percentage of dead cells was calculated based on qSCI as described in (E; n=3 wells/treatment of LNCaP cells and n=2 wells/treatment for C4-2B cells with >4000 cells/well, representative of 2 independent experiments). Statistical analysis: A=Two-way ANOVA followed by Tukey’s multiple comparisons test relative to D0; B-G=One-way ANOVA followed by Dunnett’s multiple comparisons test compared to D0/FBS (B-F) or FBS+RSL3 (G, light brown labels) and FBS+ENZ+RSL3 (G, blue labels); ns=not significant, *p<0.05 **p<0.01 ***p<0.001 ****p<0.0001

Consistent with increased PUFA content, Enz-treated LNCaP cells displayed increased membrane fluidity (Fig. 5B). Regarding the mechanisms underlying the increase in desaturation and elongation of PUFAs, we noted a significant reduction of PUFA containing storage lipids (CEs and TAGs; Figs. S3G and S3H). Apart from enzymes that play primarily a role in the elongation of very long FAs (>24 carbon atoms) and the desaturation of sphingolipids and mono-unsaturation of FAs (*ELOVL4* [47], *DEGS2* and *SCD5*), mRNA expression of most FA elongases and desaturases was significantly decreased by Enz (Fig. 5C. Table S1). In support of this, a review of microarray data from our LNCaP tumor xenograft model of CRPC progression [33] revealed that the expression of elongases and desaturases was overall reduced in regressing tumors and tumors at PSA nadir when compared to tumors from sham-castrated mice (Fig. S5B and S5C). Notably, this expression pattern reversed and increased in recurring tumors and CRPC when compared to tumors from sham-castrated mice. Furthermore, FA elongation and *de novo* FA synthesis share the requirement of malonyl-CoA as a C2 carbon donor, which is produced by acetyl-CoA carboxylase (ACACA). As shown in (Fig. 4B), ACACA mRNA expression was reduced, providing further support that Enz-treated cells were increasingly acquiring PUFAs from uptake of serum-derived lipids (Fig. S4D) and by mobilizing PUFAs from lipid storage (e.g. from CEs and TAGs) with ongoing therapy (Figs. S3G and S3H).

### PUFA enrichment of membrane lipids is associated with increased lipid peroxidation and dependency on GPX4 activity

High PUFA content of membrane phospholipids renders cells susceptible to oxidative stress through iron-dependent lipid peroxidation. In contrast, PUFAs stored as CEs and TAGs within lipid droplets are protected [48]. Excess amounts of PEs containing oxidized arachidonic acid (C20:4) or adrenic acid (C22:4) have been shown to trigger ferroptosis, a form of regulated, non-apoptotic cell death [12]. As shown above, Enz increased cellular levels of arachidonic acid (Fig. 4H) and PEs esterified with PUFAs (Fig. 5A). qSCI with the lipid peroxidation sensor C11-Bodipy and ratiometric analysis revealed that treatment of LNCaP and C4-2B cells for 14 days with Enz significantly increased lipid peroxidation (Fig. 5D). Glutathione peroxidase (GPX4) is critical for the reduction of toxic lipid hydroperoxides to their corresponding non-deleterious alcohols at the expense of reduced glutathione and thereby prevents cellular damage and induction of ferroptosis. LNCaP, C4-2B, DuCaP and 22RV1 cells treated with Enz or CSS displayed hypersensitivity to the GPX4 inhibitor RSL3 as indicated by significantly increased cell death (Fig. 5E and S5D). Thus, several AR-positive PCa cell lines demonstrated increased dependency on GPX4 activity in response to ATTs, a characteristic previously shown to be associated with drug-induced persister cells in other types of cancer in response to different therapies [9, 49]. Cytotoxicity of RSL3 was completely suppressed by antioxidant and lipid peroxide scavenger Trolox, a vitamin E derivative, and ferroptosis inhibitor Ferrostatin-1 (Figs. 5E and S5D), confirming ferroptosis as the major cell death mechanism. Enhanced GPX4 dependence was detected as early as 4-8 days after commencement of ATTs (Enz and ADT) in LNCaP and C4-2B cells (Fig. S5E) and was fully reverted when cells were given a therapy holiday for three or more days (data not shown). Furthermore, mRNA levels of several key factors of ferroptosis and redox homeostasis, including the recently identified ferroptosis suppressor protein 1 (FSP1/AIFM2) [10, 11] were significantly altered by Enz in LNCaP cells (Fig. 5F), and gene ontology analysis of protein expression changes (Fig. S1B) showed significant enrichment of pathways related to oxidative stress and selenocysteine metabolism which, an amino acid critical for the antioxidant function of selenoproteins (e.g. glutathione peroxidases, thioredoxin reductases, iodothyronine deiodinases, Fig. S5F).

Next, we tested if above processes associated with lipid remodeling are critical mechanisms that underpin GPX4 dependence of PCa persister cells. Indeed, development of ferroptosis hypersensitivity, as measured by RSL3-induced cell death, was significantly suppressed in LNCaP (Fig. 5G) and C4-2B cells (data not shown) when Enz was combined with non-toxic doses of inhibitors directed against lipase activity (Orlistat and CAY10499) [50] or FA delta-6 desaturation (SC26196) [51]. Co-inhibition of the critical lipogenesis transcription factor SREBF1 with Fatostatin [52] or ACACA (acetyl-CoA to malonyl-CoA conversion) and SCD1 (FA desaturation) with dual inhibitor TOFA [53] showed no or only limited reduction in ferroptosis sensitivity (Fig. 5G). The high protective efficiency of broad spectrum lipase inhibitor CAY10499 and FADS2 desaturase inhibitor SC26196 against RSL3-induced cell death in ENZ-treated LNCaP cells was comparable to that of Trolox (Fig. 5G) or Ferrostatin-1 (data not shown), suggesting that the co-inhibited processes are fundamental for developing Enz-induced GPX4 dependence. Similar results were observed when LNCaP cells were cultured in CSS, though the effect and efficacy of some co-treatments were different when compared to direct inhibition of AR with Enz, including that of fatostatin (Fig. S5G), suggesting that changes to the media composition (e.g. free FA levels, Fig. S4C) by charcoal-stripping might have confounding effects. In support of this, supplementation of growth media with the PUFA arachidonic acid strongly intensified RSL3 sensitivity in parental, non-ATT treated cells (Fig. 5G), whereas reducing serum concentration was protective (data not shown). This suggested a functional link of PUFA uptake mechanisms with ferroptosis sensitivity. Thus, interference with lipid supply and lipid remodeling pathways that were altered by ATTs (Enz and ADT) efficiently suppressed the acquisition of ferroptosis hypersensitivity. This demonstrated that therapy-induced GPX4 dependence can be modulated by rationally informed co-treatments targeting lipid remodeling and by altering PUFA and antioxidant levels of serum.

As shown above, acquisition of the ATT-induced persister phenotype was associated with strongly reduced proliferation, enhanced tolerance to several PCa-relevant chemotherapeutics and ferroptosis hypersensitivity. Ferroptosis induction as a therapeutic strategy shows promise in multiple malignancies [54]. However, dietary PUFA intake and antioxidant supplementation are factors that could alter ferroptosis sensitivity as suggested by our *in vitro* findings (Fig. 5G). Moreover, acquisition of fully developed therapy resistance is associated with reactivation of cell proliferation and enhanced metabolic activity and oxidative stress [6, 8, 34, 35], suggesting dynamic changes to ferroptosis sensitivity and a limited therapeutic window. Hence, we sought to identify inhibitors of metabolic pathways that both proliferating parental cells and quiescent persister cells were critically dependent on, promising a larger therapeutic window and limiting the likely effects of changes in the proliferation status on drug sensitivity. Our transcriptional, translational and functional analyses above indicated that ATTs increased cholesterol uptake through receptor-mediated endocytosis (LDLR and SCARB1) and lysosomal cholesterol processing. Indeed, Enz treatment of LNCaP and C4-2B cells and culture of LNCaP cells in CSS significantly increased toxicity of U18666A (Fig. 5H), an inhibitor of the lysosomal Niemann-Pick C1 protein (NPC1) which together with NPC2 is critical for lysosome to cytoplasm transport of exogenous cholesterol [55]. Notably, NPC1 and NPC2 protein expression were increased in localized PCa tumors and further increased in bone metastatic lesions when compared to normal prostate gland (Fig. S5H). Lysosomal activity has also previously associated with the development of drug resistance in cancer [56]. Thus, targeting the supply of exogenous cholesterol via NPC1 antagonism might present a promising co-treatment strategy to extend the efficacy of ATTs and delay the development of therapy resistance in PCa, a currently unmet clinical need.

## Discussion

Despite initial tumor regression following targeted cancer therapies, development of drug resistance and disease progression ultimately leading to patient death remain major clinical challenges of many pharmacological interventions in various types of cancer, including PCa. Previous studies of resistance to androgen receptor-targeted therapies (ATTs) in PCa analyzed models after extensive treatment periods (months to years) with fully developed resistance as indicated by reactivated cell proliferation [6, 8, 34, 35]. Different to these studies, we hypothesized that delineating early events of ATT-induced metabolic reprogramming might provide novel co-targets that could extend the efficacy of ATTs, thereby delaying therapy failure and disease progression to currently incurable CRPC and Enz-resistant CRPC.

Our longitudinal study of the early adaptive response to androgen deprivation therapy (ADT) and Enz (0-21 days) in PCa cells made several key discoveries related to reprogrammed lipid metabolism (summarized in Fig. 6). Moreover, our work provided several novel potential targets for intervention strategies to combat ATT resistance that need pre-clinical evaluation. Importantly, our study is first in class to demonstrate that therapy-induced lipid remodeling and lipid supply plasticity are critical mechanisms that underpin GPX4 dependence and ferroptosis hypersensitivity in persister cells. Thus, our findings might have wider implications for our understanding of early events of therapy resistance since GPX4 dependence has been reported in several types of cancer in response to different targeted therapies [9, 49].

**Figure 6:**
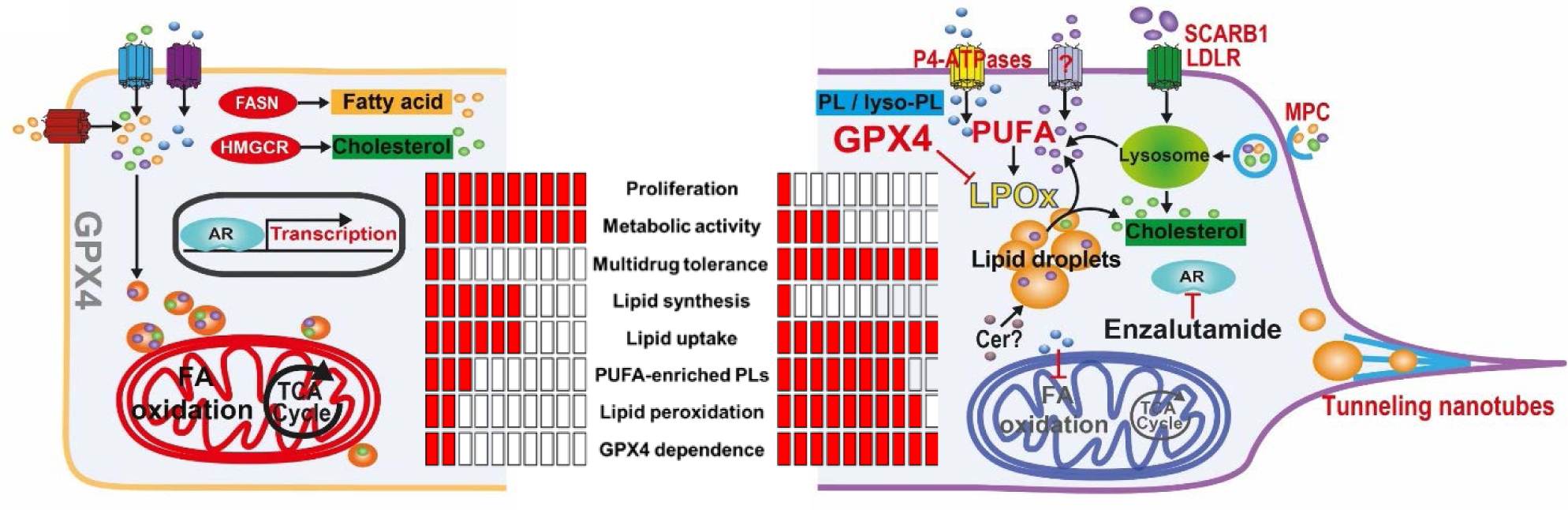
Graphic representation of changes to lipid supply and lipid remodeling that underpin therapy-induced multidrug tolerance and ferroptosis hypersensitivity in PCa cells. In the androgen-stimulated, proliferative state (left), the cellular lipid pool is fueled by both *de novo* lipogenesis (DNL) of cholesterol (green) and FAs (orange) and lipid uptake. Mitochondrial FA oxidation and TCA cycle activity provide energy and precursor for biomass synthesis. In the therapy-induced quiescent state, DNL is reduced, whereas lipid uptake through cargo-selective (lipid transporters) and non-selective transport mechanisms (tunneling nanotubes and macropinocytosis, MPC) is enhanced. Therapy-induced lipid remodeling includes mobilization of storage lipids (CEs and TAGs) through lipases, increased levels of all major phospholipid species and their enrichment with PUFAs (purple), leading to increased membrane fluidity, lipid peroxidation (LPOx) and GPX4 dependence. Increased and decreased activities are highlighted in red and gray, respectively. Cer, ceramides.

Consistent with previous studies that showed that subpopulations of breast and lung cancer cells, called drug-induced persister cells, entered a quiescent state in response to anti-cancer treatments ([9, 49] and reviewed in [57]), our functional studies showed that ATT-induced cellular quiescence was characterized by decreased proliferation, ATP production and mitochondrial activity (Figs. 2C, S2C and S2D). Previous studies have also reported a quiescence state of surviving cells after three weeks of androgen-deprivation [6, 34, 35] and re-initiation of proliferation after ~6 months of treatment [6].

Unexpectedly, in light of cellular quiescence and low bioenergetic activity, Enz-induced remodeling of all major lipid classes (an energy-intensive process) was characterized by substantial accumulation of phospholipids and depletion of typical storage lipids (TAGs and CEs, Figs. 3 and S3) resident in lipid droplets (LDs). Yet, LD number and size (Fig. 3B) were increased. Therapy-induced LD formation in the context of growth inhibition has been previously demonstrated in cancer cells in response to oncogenic signaling inhibition [58], however the lipid composition of LDs was not delineated. It was recently shown that ceramides are converted to acylceramides via DGAT2 and stored in lipid droplets [59]. While DGAT1 and DGAT2 are both involved in TAG synthesis, DGAT2 also drives acylceramide synthesis and subsequent storage in lipid droplets. DGAT2 expression was upregulated in Enz-treated cells while DGAT1 was downregulated (Table S1), suggesting that increased acylceramide synthesis and lipid droplet storage caused the measured changes to lipid droplet morphometry. Ceramide metabolism was also highlighted by our comparative gene signature analysis (Fig. S1C). Given the heterogeneous nature of lipid droplets within cell populations [60] and that their accumulation has recently been described as a mechanism of drug resistance in colorectal cancer [61], renal cell carcinoma [58], and breast cancer cell lines [62], further investigations into lipid droplet biology under therapy stress are warranted.

Aimed to identify the source of increased lipid content, stable isotope carbon tracing revealed little glycolytic contribution to FA and cholesterol synthesis (data not shown) and a 70% reduction of cholesterol synthesis from acetate-derived carbon (Fig. 4A). While lipogenesis can be fueled by other carbon sources (glutamine and FAs), enhanced lipid synthesis from alternative fuels was not supported by the measured bioenergetic, redox and mitochondrial activity statuses (Fig. 2C). Moreover, the transcriptome and proteome changes of Enz-treated LNCaP cells collectively indicated a reduction in oxidative phosphorylation (Fig. 2B) and, more importantly, a decrease of multiple key enzymes of *de novo* FA and cholesterol biosynthesis (Fig. 4B). These results were consistent with reduced toxicity of several lipogenesis inhibitors (Figs. 2D and S2G). Instead, therapy-induced lipid remodeling and increased lipid content was fueled by enhanced lipid uptake via cargo-selective (transporters, Fig. 4C-F) and non-selective uptake mechanisms (macropinocytosis and TNTs, Figs. S4E and S4F). We frequently observed mitochondria and lipid droplets within Enz-induced TNTs (Fig. S4F), suggesting that these conduits of intercellular transport might contribute considerable amounts of lipid biomass. Similarly, macropinocytosis of cell debris including membrane components have been previously shown to significantly contribute to the cellular lipid pool [45]. Notably, the lipid supply switch to uptake excluded uptake of saturated FAs as indicated by reduced uptake of C16:0-Bodipy after extended ATT treatments (Enz and ADT, Fig. 4G), indicating that other uptake mechanisms or transporters were responsible for the increased cellular levels of saturated FAs and PUFAs. Together, these results demonstrate that ATTs induced a switch of lipid supply from a mix of *de novo* synthesis and exogenous sources to predominantly uptake of exogenous lipids via multiple mechanisms. Thus, lipid supply plasticity is a novel metabolic mechanism associated with ATT resistance development in PCa, and enhanced lipid scavenging can serve purposes other than biomass and energy production to support malignant cell proliferation. It is therefore critical to acknowledge that lipid supply plasticity during the development of ATT resistance limits the therapeutic window of individual treatments directed against either uptake or synthesis.

Only recently has attention been given to lipid uptake and to map the lipid transporter landscape in cancer cells [42, 63], including our own work in PCa [16]. The observed Enz-induced increase in protein expression of both LDLR and SCARB1 along with increased transcript levels of several other lipid transporters (Figs. 4E and 4F), supported the functional measurements obtained by qSCI of fluorescent lipid probes. They were concordant with previously reported transcriptional changes observed in our LNCaP tumor xenograft progression model in which recurring and CRPC tumors expressed higher levels of several lipid transporters and contained higher lipid content, including PUFAs, compared to tumors resected before castration [33]. Furthermore, increased expression of several lipid transporters were also observed in bone metastasis of CRPC patients (Fig. S4B, [16]); however, whether this is indirectly caused by the lipid-rich environment present in adult bone tissue (high adiposity) or directly by ATTs remains to be shown.

A major limitation of pharmacological targeting of PCa lipid supply is the scarcity of inhibitors and the incomplete understanding of mass contributions of different lipid supply pathways to the cellular lipid pool, their redundancies and cross-talk and compensatory capacities in response to reduced lipid synthesis, i.e. lipid supply plasticity. Lipogenesis on the other hand has remained a major target in cancer therapy, yielding an arsenal of inhibitors; unfortunately, with limited clinical success. There is increasing acknowledgement that various lipid scavenging pathways, including transporter-mediated uptake, tunneling nanotubes [46], lipid-conjugated albumin uptake [64] and macropinocytosis [18] are critical supply routes of exogenous lipids fueling bioenergetic processes and biomass production that underpin tumor growth and survival. Recent studies of cancer cells expressing the lipogenic phenotype, including PCa, estimated that >70% of the lipid-derived carbon biomass is derived from uptake and only 30% from synthesis based on carbon tracing of serum-derived free FAs [65, 66]. However, these estimates are limited by the fact that the serum lipidome is highly complex and that >95% of serum FAs are acyl conjugates across all lipid classes, including TAGs, CEs and phospholipids. Thus, the carbon contribution from the collective uptake of all lipid classes to lipid biomass in proliferating cancer cells is likely to be even higher than the above estimates. Despite this knowledge gap, we showed that blocking cholesterol uptake via NPC1 inhibition with U18666A (Fig. 5H) is a promising strategy to exploit the increased demand of ATT-treated PCa cells for exogenous cholesterol (Fig. 4C). Cholesterol is a critical storage lipid (as acyl-ester in lipid droplets) and serves as a precursor for the steroidogenic pathway in which testosterone can be endogenously synthesized in PCa tumors [67], and that activation of the steroidogenic pathway is an ATT-induced adaptive response contributing to CRPC progression despite castrate levels of circulating androgens [68–70]. Additionally, non-esterified, membrane cholesterol is shown to affect lipid raft composition which may influence oncogenic signaling [71], further highlighting the critical dependence of PCa cells on cholesterol for growth and survival, particularly under therapy stress. However, targeting cholesterol supply as a therapeutic strategy in PCa solely focused on the inhibition of *de novo* cholesterol synthesis, i.e. inhibition of the early steps of the mevalonate pathway (HGMCR with statins) [72]. Thus, co-targeting of lysosomal processing of exogenous cholesterol with U18666A and synthesis pathways with statins in the context of ATTs may prove a potent strategy that targets aforementioned aspects of cholesterol metabolism as well as sensitizes to ferroptosis through loss of intermediates of the mevalonate synthesis pathway (i.e. isopentenyl pyrophosphate and farnesyl diphosphate) that are critical for GPX4 [11] and FSP1 activities [13, 14].

In addition to Enz-enhanced lipid uptake and content, our work described extensive lipid remodeling that was characterized by increased FA elongation and desaturation (Fig. 5A) of phospholipids. Both processes have previously been associated with PCa incidence and aggressiveness [73, 74]. Phospholipids containing long-chain FAs along with membrane bound cholesterol are critical for membrane stabilization and are involved in lipid raft formation where oncogenic growth signaling occurs [71, 74]. Enz-induced high PUFA levels in phospholipids were associated with increased membrane fluidity (Fig 5B) which is thought to play a role in reduced sensitivity to chemotherapy reagents (i.e. multidrug tolerance, Fig. 2D) [75] and increases permeability and cell mobility and facilitates membrane-centered biological processes (vesicle formation, membrane fusion, cell-cell interaction, modulation of membrane enzymes, receptors, and transporters) [76]. Increased PUFA content of phospholipids does, however, generate a metabolic vulnerability [77]: increased oxidative damage through reactive oxygen species (ROS)-induced lipid peroxidation (Fig. 5D). If not enzymatically repaired by GPX4 at the expense of reduced glutathione, accumulation of lipid hydroperoxides can lead to cell death via ferroptosis. Consistent with this, Enz-treated PCa cells showed reduced reductive power and hypersensitivity to selective GPX4 inhibitor, RSL3 (Fig. 5E). ADT and Enz-induced sensitivity to GPX4 inhibition of PCa cells was reminiscent of lapatinib- and vemurafenib-induced drug tolerance and emergence of “persister cells” in melanoma, breast, lung and ovarian cancer [9] and therapy-resistance-associated high-mesenchymal state cancer cells [78]. It was speculated that GPX4 dependence in these therapy-induced states relied on the FA composition of the lipid bilayers. Our results of therapy-induced lipid remodeling causing PUFA enrichment provides first-in-class evidence to support this hypothesis. More importantly, the diversity of targeted therapies causing this shared GPX4 dependence suggests that lipid remodeling leading to PUFA enrichment of phospholipids might be a common mechanism associated with therapy resistance that could be pursued as therapeutic target in multiple types of cancer. However, the therapeutic window is likely to be small and restricted to the low cycling, quiescence phase [9] because high membrane PUFA levels would not be compatible with enhanced proliferation [48] due to higher metabolic activities that increase ROS levels and oxidative stress via lipid peroxidation. Such stress would be further aggravated by higher demands in reducing power (NADPH) for the recycling of oxidized glutathione and *de novo* lipid synthesis.

Therefore, impeding the acquisition of the therapy-induced persister cell phenotype by co-targeting one or several lipid remodeling processes might be a more promising strategy that extends the efficacy of existing cancer therapies and delays resistance than combatting fully developed and often complex resistance mechanisms. By using enhanced GPX4 dependence (RSL3 sensitivity) as a biomarker of the persister cell phenotype, we demonstrated that co-targeting of several different processes of lipid remodeling protected ATT-treated PCa cells from ferroptosis hypersensitivity (Figs. 5G and S5G). In particular, the broad spectrum lipase inhibitors orlistat and CAY10499 [50] and FADS2 delta-6 desaturase inhibitor SC26196 [51] displayed high protective efficiencies against RSL3-induced cell death in ENZ-treated LNCaP cells when compared to Trolox. This demonstrated that therapy-induced GPX4 dependence can be modulated by rationally informed co-treatments targeting lipid remodeling. We noted that mRNA expression of the majority of FA elongases and desaturases, including SCD1 and FADS2, were reduced after Enz treatment for 7-21 days (Fig. 5C). Yet, inhibitors against these enzymes efficiently suppressed Enz-induced ferroptosis hypersensitivity when co-treated for 7 days (Figs 5G and S5G). Thus, the different treatment periods might explain the divergence between inhibitor activity and target expression, i.e., Enz induces a time-dependent switch from endogenous PUFA synthesis by FA elongation and desaturation to enhanced uptake of exogenous PUFAs as demonstrated for uptake of other lipid species (Figs. 4C and 4D). Notably, FADS2 activity has been recently implicated in cancer plasticity [79], though a functional role of its unsaturated FA products was yet to be identified. Our work indicates a potential functional link between FADS2 activity and therapy-induced GPX4 dependence that warrants further investigations. Whether co-targeting of lipid remodeling in combination with ATTs ultimately delays resistance development is a priority for further studies. Furthermore, the ferroptosis altering effects of PUFA and antioxidant supplementation might have implications for PCa patients undergoing ATT treatment, i.e. changing the duration until therapy failure, that need evaluation.

It is clear from this work that, despite its numerous discoveries, there are many unanswered questions regarding mechanisms of therapy-induced lipid remodeling and how this process can be harnessed to extend efficacies of existing, clinically proven therapies. The most prevalent are confirmation of the PUFA-enriched phospholipid phenotype in other types of cancer displaying therapy-induced GPX4 dependence [9, 78], investigating if dietary aspects like PUFA consumption and antioxidant supplementation affect resistance development and pre-clinical testing of co-therapies against lipid remodeling to combat ATT resistance development and disease progression to currently lethal CRPC.

## Abbreviations

ADT: Androgen deprivation therapy
AR: Androgen Receptor
ATT: Androgen receptor-targeted therapy
CE: Cholesterol ester
Cer: Ceramide
CRPC: Castration-resistant prostate cancer
CSS: Charcoal-stripped serum
DNL: De novo lipogenesis
Enz: Enzalutamide
ESI-MS: Electrospray ionization mass spectrometry
FA: Fatty acid
FDR: False discovery rate
GC/MS: Gas chromatography tandem mass spectrometry
GSVA: Gene set variation analysis
INT: Intact
LC/MS: Liquid chromatography tandem mass spectrometry
LD: Lipid droplet
LDL: Low density lipoprotein
LPC: Lyso-phosphatidylcholine
LPOx: Lipid peroxidation
MDT: Multidrug tolerance
MFI: Mean fluorescence intensity
MPC: Macropinocytosis
NAD: Nadir
PC: Phosphatidylcholine
PCa: Prostate cancer
PE: Phosphatidylethanolamine
PG: Phosphatidylglycerol
PL: Phospholipid
PS: Phosphatidylserine
PUFA: Poly-unsaturated fatty acid
qSCI: quantitative single-cell imaging
ROS: Reactive oxygen species
SM: Sphingomyelin
TAG: Triacylglycerol
TNT: Tunneling nanotube

## Availability of data and materials

Normalized gene expression data of the microarray experiment have been deposited in NCBI’s Gene Expression Omnibus (GEO) and are accessible through GEO Series accession number GSExxx. The proteomics dataset, lipid mass spectrometry data and CellProfiler analysis pipelines are available from the corresponding author on request.

## Acknowledgements

We would like to thank Prof Dr Bart Ghesquière, Mr. Abel Acosta Sanchez and the Metabolomics Expertise Center from the department of Oncology (KU Leuven) and the Center for Cancer Biology (CCB, VIB) for the ^13^C carbon tracing analyses. We thank the Proteomics Core Facility and Translational Cell Imaging Queensland located at the Translational Research Institute for their technical assistance.

## Authors’ contributions

### Conception and design

Martin C Sadowski, Kaylyn D Tousignant

### Development of methodology

Martin C Sadowski, Kaylyn D Tousignant, Anja Rockstroh, Berwyck LJ Poad, Rajesh Gupta, Stephen J Blanksby, Reuben RS Young

### Acquisition of data

Kaylyn D Tousignant, Martin C Sadowski, Ali Talebi, Anja Rockstroh

### Analysis and interpretation of data (i.e. RNAseq analysis, computational analysis)

Kaylyn D Tousignant, Martin C Sadowski, Anja Rockstroh, Atefeh Taherian Fard, Melanie Lehman, Chenwei Wang, Berwyck LJ Poad, Rajesh Gupta, Reuben RS Young, Ali Talebi, Johan V Swinnen, Colleen C Nelson

### Writing, review, and/or revision of the manuscript

Kaylyn D Tousignant, Martin C Sadowski, Melanie Lehman, Stephen J Blanksby, Colleen C Nelson

## Ethics declarations

### Ethics approval and consent to participate

Not applicable.

### Consent for publication

Not applicable

### Supplementary information

Supplementary Info

Supplementary Figures

Supplementary Table S1

Supplementary Table S2

Supplementary Table S3

Supplementary Table S4

Supplementary Table S5

Supplementary Table S6

## References

1. Litwin MS, Tan H: The diagnosis and treatment of prostate cancer: A review. JAMA 2017, 317(24):2532–2542.

2. Heinlein CA, Chang C: Androgen Receptor in Prostate Cancer. Endocrine Reviews 2004, 25(2):276–308.

3. Lonergan PE, Tindall DJ: Androgen receptor signaling in prostate cancer development and progression. Journal of Carcinogenesis 2011, 10:20.

4. Risbridger GP, Davis ID, Birrell SN, Tilley WD: Breast and prostate cancer: more similar than different. Nat Rev Cancer 2010, 10(3):205–212.

5. Dutt SS, Gao AC: Molecular mechanisms of castration-resistant prostate cancer progression. Future oncology (London, England) 2009, 5(9):1403–1413.

6. Lu S, Tsai SY, Tsai M-J: Molecular Mechanisms of Androgen-Independent Growth of Human Prostate Cancer LNCaP-AI Cells1. Endocrinology 1999, 140(11):5054–5059.

7. Höti N, Shah P, Hu Y, Yang S, Zhang H: Proteomics analyses of prostate cancer cells reveal cellular pathways associated with androgen resistance. Proteomics 2017, 17(6):10.1002/pmic.201600228.

8. Kregel S, Chen JL, Tom W, Krishnan V, Kach J, Brechka H, Fessenden TB, Isikbay M, Paner GP, Szmulewitz RZ et al: Acquired resistance to the second-generation androgen receptor antagonist enzalutamide in castration-resistant prostate cancer. Oncotarget 2016, 7(18):26259–26274.

9. Hangauer MJ, Viswanathan VS, Ryan MJ, Bole D, Eaton JK, Matov A, Galeas J, Dhruv HD, Berens ME, Schreiber SL et al: Drug-tolerant persister cancer cells are vulnerable to GPX4 inhibition. Nature 2017, 551:247.

10. Yang WS, SriRamaratnam R, Welsch ME, Shimada K, Skouta R, Viswanathan VS, Cheah JH, Clemons PA, Shamji AF, Clish CB et al: Regulation of ferroptotic cancer cell death by GPX4. Cell 2014, 156(1-2):317–331.

11. Seibt TM, Proneth B, Conrad M: Role of GPX4 in ferroptosis and its pharmacological implication. Free Radical Biology and Medicine 2019, 133:144–152.

12. Kagan VE, Mao G, Qu F, Angeli JPF, Doll S, Croix CS, Dar HH, Liu B, Tyurin VA, Ritov VB et al: Oxidized arachidonic and adrenic PEs navigate cells to ferroptosis. Nat Chem Biol 2017, 13(1):81–90.

13. Bersuker K, Hendricks J, Li Z, Magtanong L, Ford B, Tang PH, Roberts MA, Tong B, Maimone TJ, Zoncu R et al: The CoQ oxidoreductase FSP1 acts parallel to GPX4 to inhibit ferroptosis. Nature 2019.

14. Doll S, Freitas FP, Shah R, Aldrovandi M, da Silva MC, Ingold I, Grocin AG, Xavier da Silva TN, Panzilius E, Scheel CH et al: FSP1 is a glutathione-independent ferroptosis suppressor. Nature 2019.

15. Locke JA, Nelson CC, Adomat HH, Hendy SC, Gleave ME, Guns ES: Steroidogenesis inhibitors alter but do not eliminate androgen synthesis mechanisms during progression to castration-resistance in LNCaP prostate xenografts. J Steroid Biochem Mol Biol 2009, 115(3-5):126–136.

16. Tousignant KD, Rockstroh A, Taherian Fard A, Lehman ML, Wang C, McPherson SJ, Philp LK, Bartonicek N, Dinger ME, Nelson CC et al: Lipid uptake is an androgen-enhanced lipid supply pathway associated with prostate cancer disease progression and bone metastasis. 2019:molcanres.1147.2018.

17. Desir S, Dickson EL, Vogel RI, Thayanithy V, Wong P, Teoh D, Geller MA, Steer CJ, Subramanian S, Lou E: Tunneling nanotube formation is stimulated by hypoxia in ovarian cancer cells. Oncotarget 2016, 7(28):43150–43161.

18. Finicle BT, Jayashankar V, Edinger AL: Nutrient scavenging in cancer. Nature Reviews Cancer 2018, 18(10):619–633.

19. Jaishy B, Abel ED: Lipids, lysosomes, and autophagy. Journal of lipid research 2016, 57(9):1619–1635.

20. Schmittgen TD, Livak KJ: Analyzing real-time PCR data by the comparative C(T) method. Nat Protoc 2008, 3(6):1101–1108.

21. Smyth GK: Limma: linear models for microarray data. 2005:397–420.

22. Hänzelmann S, Castelo R, Guinney J: GSVA: gene set variation analysis for microarray and RNA-Seq data. BMC Bioinformatics 2013, 14(1):7.

23. Subramanian A, Tamayo P, Mootha VK, Mukherjee S, Ebert BL, Gillette MA, Paulovich A, Pomeroy SL, Golub TR, Lander ES et al: Gene set enrichment analysis: a knowledge-based approach for interpreting genome-wide expression profiles. Proc Natl Acad Sci U S A 2005, 102(43):15545–15550.

24. Eden E, Navon R, Steinfeld I, Lipson D, Yakhini Z: GOrilla: a tool for discovery and visualization of enriched GO terms in ranked gene lists. BMC Bioinformatics 2009, 10:48.

25. Kolde R, Vilo J: GOsummaries: an R Package for Visual Functional Annotation of Experimental Data. F1000Res 2015, 4:574.

26. Li S, Fong K-w, Gritsina G, Zhang A, Zhao JC, Kim J, Sharp A, Yuan W, Aversa C, Yang XJ et al: Activation of MAPK Signaling by CXCR7 Leads to Enzalutamide Resistance in Prostate Cancer. 2019, 79(10):2580–2592.

27. Li Q, Deng Q, Chao H-P, Liu X, Lu Y, Lin K, Liu B, Tang GW, Zhang D, Tracz A et al: Linking prostate cancer cell AR heterogeneity to distinct castration and enzalutamide responses. Nature Communications 2018, 9(1):3600.

28. Rajan P, Sudbery IM, Villasevil MEM, Mui E, Fleming J, Davis M, Ahmad I, Edwards J, Sansom OJ, Sims D et al: Next-generation Sequencing of Advanced Prostate Cancer Treated with Androgen-deprivation Therapy. European Urology 2014, 66(1):32–39.

29. Morpheus: https://software.broadinstitute.org/morpheus.

30. Levrier C, Sadowski MC, Nelson CC, Healy PC, Davis RA: Denhaminols A-H, dihydro-beta-agarofurans from the endemic Australian rainforest plant Denhamia celastroides. J Nat Prod 2015, 78(1):111–119.

31. Carpenter AE, Jones TR, Lamprecht MR, Clarke C, Kang IH, Friman O, Guertin DA, Chang JH, Lindquist RA, Moffat J et al: CellProfiler: image analysis software for identifying and quantifying cell phenotypes. Genome Biology 2006, 7(10):R100.

32. Butte W: Rapid method for the determination of fatty acid profiles from fats and oils using trimethylsulphonium hydroxide for transesterification. Journal of Chromatography A 1983, 261:142–145.

33. Locke JA, Guns ES, Lehman ML, Ettinger S, Zoubeidi A, Lubik A, Margiotti K, Fazli L, Adomat H, Wasan KM et al: Arachidonic acid activation of intratumoral steroid synthesis during prostate cancer progression to castration resistance. Prostate 2010, 70(3):239–251.

34. Yu P, Duan X, Cheng Y, Liu C, Chen Y, Liu W, Yin B, Wang X, Tao Z: Androgen-independent LNCaP cells are a subline of LNCaP cells with a more aggressive phenotype and androgen suppresses their growth by inducing cell cycle arrest at the G1 phase. Int J Mol Med 2017, 40(5):1426–1434.

35. Xu G, Wu J, Zhou L, Chen B, Sun Z, Zhao F, Tao Z: Characterization of the Small RNA Transcriptomes of Androgen Dependent and Independent Prostate Cancer Cell Line by Deep Sequencing. PLOS ONE 2010, 5(11):e15519.

36. Pfeiffer MJ, Smit FP, Sedelaar JPM, Schalken JA: Steroidogenic enzymes and stem cell markers are upregulated during androgen deprivation in prostate cancer. Mol Med 2011, 17(7-8):657–664.

37. Swinnen JV, Esquenet M, Goossens K, Heyns W, Verhoeven G: Androgens stimulate fatty acid synthase in the human prostate cancer cell line LNCaP. Cancer Res 1997, 57(6):1086–1090.

38. Swinnen JV, Van Veldhoven PP, Esquenet M, Heyns W, Verhoeven G: Androgens markedly stimulate the accumulation of neutral lipids in the human prostatic adenocarcinoma cell line LNCaP. Endocrinology 1996, 137(10):4468–4474.

39. Greenspan P ME, Fowler S: Nile red: a selective fluorescent stain for intracellular lipid droplets. The Journal of Cell Biology 1985, 100(3):965–973.

40. Maxfield FR, Wüstner D: Analysis of cholesterol trafficking with fluorescent probes. Methods in cell biology 2012, 108:367–393.

41. Christie WW: Rapid separation and quantification of lipid classes by high performance liquid chromatography and mass (light-scattering) detection. J Lipid Res 1985, 26(4):507–512.

42. Iglesias-Gato D, Thysell E, Tyanova S, Crnalic S, Santos A, Lima TS, Geiger T, Cox J, Widmark A, Bergh A et al: The proteome of prostate cancer bone metastasis reveals heterogeneity with prognostic implications. Clinical Cancer Research 2018.

43. Ikonen E: Cellular cholesterol trafficking and compartmentalization. Nat Rev Mol Cell Biol 2008, 9(2):125–138.

44. Goldstein JL, Anderson RG, Brown MS: Receptor-mediated endocytosis and the cellular uptake of low density lipoprotein. Ciba Found Symp 1982(92):77–95.

45. Kim SM, Nguyen TT, Ravi A, Kubiniok P, Finicle BT, Jayashankar V, Malacrida L, Hou J, Robertson J, Gao D et al: PTEN Deficiency and AMPK Activation Promote Nutrient Scavenging and Anabolism in Prostate Cancer Cells. 2018, 8(7):866–883.

46. Astanina K, Koch M, Jüngst C, Zumbusch A, Kiemer AK: Lipid droplets as a novel cargo of tunnelling nanotubes in endothelial cells. Scientific reports 2015, 5:11453–11453.

47. Agbaga M-P, Brush RS, Mandal MNA, Henry K, Elliott MH, Anderson RE: Role of Stargardt-3 macular dystrophy protein (ELOVL4) in the biosynthesis of very long chain fatty acids. 2008, 105(35):12843–12848.

48. Bailey Andrew P, Koster G, Guillermier C, Hirst Elizabeth MA, MacRae James I, Lechene Claude P, Postle Anthony D, Gould Alex P: Antioxidant Role for Lipid Droplets in a Stem Cell Niche of Drosophila. Cell 2015, 163(2):340–353.

49. Ramirez M, Rajaram S, Steininger RJ, Osipchuk D, Roth MA, Morinishi LS, Evans L, Ji W, Hsu C-H, Thurley K et al: Diverse drug-resistance mechanisms can emerge from drug-tolerant cancer persister cells. Nature communications 2016, 7:10690–10690.

50. Iglesias J, Lamontagne J, Erb H, Gezzar S, Zhao S, Joly E, Truong VL, Skorey K, Crane S, Madiraju SR et al: Simplified assays of lipolysis enzymes for drug discovery and specificity assessment of known inhibitors. J Lipid Res 2016, 57(1):131–141.

51. Obukowicz MG, Welsch DJ, Salsgiver WJ, Martin-Berger CL, Chinn KS, Duffin KL, Raz A, Needleman P: Novel, selective delta6 or delta5 fatty acid desaturase inhibitors as antiinflammatory agents in mice. J Pharmacol Exp Ther 1998, 287(1):157–166.

52. Li X, Chen YT, Hu P, Huang WC: Fatostatin displays high antitumor activity in prostate cancer by blocking SREBP-regulated metabolic pathways and androgen receptor signaling. Mol Cancer Ther 2014, 13(4):855–866.

53. Mason P, Liang B, Li L, Fremgen T, Murphy E, Quinn A, Madden SL, Biemann HP, Wang B, Cohen A et al: SCD1 inhibition causes cancer cell death by depleting mono-unsaturated fatty acids. PLoS One 2012, 7(3):e33823.

54. Feng H, Stockwell BR: Unsolved mysteries: How does lipid peroxidation cause ferroptosis? PLoS Biol 2018, 16(5):e2006203.

55. Pfeffer SR: NPC intracellular cholesterol transporter 1 (NPC1)-mediated cholesterol export from lysosomes. J Biol Chem 2019, 294(5):1706–1709.

56. Zhitomirsky B, Assaraf YG: Lysosomes as mediators of drug resistance in cancer. Drug Resistance Updates 2016, 24:23–33.

57. Vallette FM, Olivier C, Lézot F, Oliver L, Cochonneau D, Lalier L, Cartron P-F, Heymann D: Dormant, quiescent, tolerant and persister cells: four synonyms for the same target in cancer. Biochemical Pharmacology 2018.

58. Lue HW, Podolak J, Kolahi K, Cheng L, Rao S, Garg D, Xue CH, Rantala JK, Tyner JW, Thornburg KL et al: Metabolic reprogramming ensures cancer cell survival despite oncogenic signaling blockade. Genes Dev 2017, 31(20):2067–2084.

59. Senkal CE, Salama MF, Snider AJ, Allopenna JJ, Rana NA, Koller A, Hannun YA, Obeid LM: Ceramide Is Metabolized to Acylceramide and Stored in Lipid Droplets. Cell metabolism 2017, 25(3):686–697.

60. Petan T, Jarc E, Jusović M: Lipid Droplets in Cancer: Guardians of Fat in a Stressful World. Molecules (Basel, Switzerland) 2018, 23(8):1941.

61. Cotte AK, Aires V, Fredon M, Limagne E, Derangère V, Thibaudin M, Humblin E, Scagliarini A, de Barros J-PP, Hillon P et al: Lysophosphatidylcholine acyltransferase 2-mediated lipid droplet production supports colorectal cancer chemoresistance. Nature communications 2018, 9(1):322–322.

62. Hultsch S, Kankainen M, Paavolainen L, Kovanen R-M, Ikonen E, Kangaspeska S, Pietiäinen V, Kallioniemi O: Association of tamoxifen resistance and lipid reprogramming in breast cancer. BMC cancer 2018, 18(1):850–850.

63. Iglesias-Gato D, Wikström P, Tyanova S, Lavallee C, Thysell E, Carlsson J, Hägglöf C, Cox J, Andrén O, Stattin P et al: The Proteome of Primary Prostate Cancer. European Urology 2016, 69(5):942–952.

64. Bern M, Sand KMK, Nilsen J, Sandlie I, Andersen JT: The role of albumin receptors in regulation of albumin homeostasis: Implications for drug delivery. Journal of Controlled Release 2015, 211:144–162.

65. Hosios Aaron M, Hecht Vivian C, Danai Laura V, Johnson Marc O, Rathmell Jeffrey C, Steinhauser Matthew L, Manalis Scott R, Vander Heiden Matthew G: Amino Acids Rather than Glucose Account for the Majority of Cell Mass in Proliferating Mammalian Cells. Developmental Cell 2016, 36(5):540–549.

66. Balaban S, Nassar ZD, Zhang AY, Hosseini-Beheshti E, Centenera MM, Schreuder M, Lin HM, Aishah A, Varney B, Liu-Fu F et al: Extracellular Fatty Acids Are the Major Contributor to Lipid Synthesis in Prostate Cancer. Mol Cancer Res 2019.

67. Locke JA, Guns ES, Lubik AA, Adomat HH, Hendy SC, Wood CA, Ettinger SL, Gleave ME, Nelson CC: Androgen levels increase by intratumoral de novo steroidogenesis during progression of castration-resistant prostate cancer. Cancer Res 2008, 68(15):6407–6415.

68. Dillard PR, Lin M-F, Khan SA: Androgen-independent prostate cancer cells acquire the complete steroidogenic potential of synthesizing testosterone from cholesterol. Molecular and Cellular Endocrinology 2008, 295(1-2):115–120.

69. Leon CG, Locke JA, Adomat HH, Etinger SL, Twiddy AL, Neumann RD, Nelson CC, Guns ES, Wasan KM: Alterations in cholesterol regulation contribute to the production of intratumoral androgens during progression to castration-resistant prostate cancer in a mouse xenograft model. Prostate 2010, 70(4):390–400.

70. Ghayee HK, Auchus RJ: Basic concepts and recent developments in human steroid hormone biosynthesis. Reviews in Endocrine and Metabolic Disorders 2007, 8(4):289–300.

71. Zhuang L, Kim J, Adam RM, Solomon KR, Freeman MR: Cholesterol targeting alters lipid raft composition and cell survival in prostate cancer cells and xenografts. Journal of Clinical Investigation 2005, 115(4):959–968.

72. Gordon JA, Midha A, Szeitz A, Ghaffari M, Adomat HH, Guo Y, Klassen TL, Guns ES, Wasan KM, Cox ME: Oral simvastatin administration delays castration-resistant progression and reduces intratumoral steroidogenesis of LNCaP prostate cancer xenografts. Prostate Cancer Prostatic Dis 2016, 19(1):21–27.

73. Peck B, Schug ZT, Zhang Q, Dankworth B, Jones DT, Smethurst E, Patel R, Mason S, Jiang M, Saunders R et al: Inhibition of fatty acid desaturation is detrimental to cancer cell survival in metabolically compromised environments. Cancer & Metabolism 2016, 4:6.

74. Tamura K, Makino A, Hullin-Matsuda F, Kobayashi T, Furihata M, Chung S, Ashida S, Miki T, Fujioka T, Shuin T et al: Novel Lipogenic Enzyme ELOVL7 Is Involved in Prostate Cancer Growth through Saturated Long-Chain Fatty Acid Metabolism. Cancer Research 2009, 69(20):8133–8140.

75. Corsetto P, Colombo I, Kopecka J, Rizzo A, Riganti C: ω-3 Long Chain Polyunsaturated Fatty Acids as Sensitizing Agents and Multidrug Resistance Revertants in Cancer Therapy. International Journal of Molecular Sciences 2017, 18(12):2770.

76. van Meer G, Voelker DR, Feigenson GW: Membrane lipids: where they are and how they behave. Nature reviews Molecular cell biology 2008, 9(2):112–124.

77. Bailey AP, Koster G, Guillermier C, Hirst EM, MacRae JI, Lechene CP, Postle AD, Gould AP: Antioxidant Role for Lipid Droplets in a Stem Cell Niche of Drosophila. Cell 2015, 163(2):340–353.

78. Viswanathan VS, Ryan MJ, Dhruv HD, Gill S, Eichhoff OM, Seashore-Ludlow B, Kaffenberger SD, Eaton JK, Shimada K, Aguirre AJ et al: Dependency of a therapy-resistant state of cancer cells on a lipid peroxidase pathway. Nature 2017, 547:453.

79. Vriens K, Christen S, Parik S, Broekaert D, Yoshinaga K, Talebi A, Dehairs J, Escalona-Noguero C, Schmieder R, Cornfield T et al: Evidence for an alternative fatty acid desaturation pathway increasing cancer plasticity. Nature 2019, 566(7744):403–406.

