## Supplementary Info and Figures for "Therapy-induced lipid uptake and remodeling underpin ferroptosis hypersensitivity in prostate cancer"

### Supplementary Methods

#### Cell culture

22RV1 (ATCC, CRL-2505) and DuCaP cells (ATCC, CVCL_2025) were cultured in RPMI medium (Thermo Fisher) supplemented with 5% and 10% FBS, respectively. Medium was changed every 3 days and cells were incubated at 37°C in 5% CO2. For treatments representing androgen-deprivation therapy, cells were cultured in RPMI supplemented with 5% charcoal stripped-serum (CSS). For AR targeted therapies, cells were cultured in 5% or 10% FBS with AR-antagonist Enzalutamide (Enz, 10 µM). Cells were passaged at approximately 80% confluency by trypsinization. Cell lines were genotyped in March 2018 by Genomics Research Centre (Brisbane) and routinely tested for mycoplasma infection.

#### Measurements of cellular ATP levels and reductive power

Cellular ATP levels and reducing power were measured by CellTiter-Glo luminescence and PrestoBlue fluorescence assays, respectively. Briefly, LNCaP cells cultured for the indicted times in either 5% FBS+Enzalutamide (10 µM) or 5% CSS were harvested by trypsinization and seeded into 96-well tissue culture plates (n=3 biological replicates with 4 wells/sample, Corning, Corning, NY, USA) at a density of 6,000 cells/well in their corresponding types of media (RPMI medium supplemented with either 5% FBS, 5%FBS+Enzalutamide (10 µM) or 5% CSS). After 48 hours, CellTiter-Glo and Presto Blue assays were performed according to the manufacturer’s instructions. Luminescence and fluorescence were in a FLUOstar Omega microplate reader (Bio Tek). Measurements were normalized based on total cell count by DNA staining with Hoechst 33342.

#### Quantitative single cell imaging of metabolic parameters by fluorescent microscopy

Prior to seeding, imaging plates were coated with 150 µl Poly-l-ornithine (PLO, Sigma) and washed with PBS to increase cell attachment. PCa cell lines pre-treated with either 5% FBS+Enzalutamide (10 µM) or 5% CSS were harvested three days prior completion of the indicated treatment times by trypsinization and seeded into PLO-coated 96-well Ibidi optical plates (n=3 biological replicates with 4 wells/sample) at a density of 6,000 cells/well in their corresponding types of media (RPMI medium supplemented with either 5% FBS (D0), 5%FBS+Enzalutamide (10 µM) or 5% CSS). To measure glucose uptake, media was removed after three days (completion of full treatment period), and cells were incubated at 37°C for one hour in 65 µL/well solution of glucose-free RPMI (Thermo Fisher) supplemented with 2-NBDG (2-(N-(7_Nitrobenz-2-oxa-1,3-diazol-4-yl)Amino)-2-Deoxyglucose, 50 µM, Thermo Fisher). After incubation, cells were washed and fixed with 4% PFA. Cellular DNA and F-actin was counterstained with DAPI and Alexa Fluor 647 Phalloidin (Thermo Fisher), respectively. Approximately 750 cells/well (3-8 wells/sample) were imaged. For measuring macropinocytosis, growth media was exchanged with 80 µL/well of serum-free RPMI media (0.2% lipid-free BSA) supplemented with 70 kDa dextran-TR (1 mg/mL, Thermo Fisher), and cells were incubated at 37°C for one hour. DNA and lysosomes were counterstained with Hoechst 33342 and LysoTracker Red DND-99 (5 µM, Thermo Fisher). After washing with serum-free media, cells were imaged live. Autophagy was measured by live imaging of ATT-treated cells stained with Cyto-ID reagent (2 µL/mL, Enzo Life Sciences) for 60 min at 37°C. Total mitochondrial mass per cell (independent of membrane potential) was measured by live imaging of MitoTracker Green FM-stained cells (200 nM, Thermo Fisher). All images were acquired with the InCell 2200 automated fluorescence microscope system (GE Healthcare Life Sciences) at 10x and 40x magnifications. Quantitative image analysis of mean fluorescence intensity (MFI) and morphometric data was performed with Cell Profiler Software (Broad Institute).

**GC-MS FAME analysis of serum**

Extractions of lipids from FBS and CSS (Sigma Aldrich) used for cell culture experiments in these studies were performed at a 9:1 MTBE:serum ratio with added internal standard (SPLASH Lipid-o-mix deuterated internal standard obtained from Avanti Polar Lipids, Alabaster, AL) and n-Nonadecanoic acid (C19:0) and briefly vortexed. 770 µL MTBE was added and mixture was incubated for 1 h at room temperature in a shaker. Phase separation was induced by adding 200 µL $NH_4CH_3CO_2$ (150 mM). After vortexing for 20 s, the sample was centrifuged at 2,000 g for 5 min. The upper (organic) phase was collected and stored at -80°C, then diluted into 2:1 MeOH:$CHCl_3$ with 7.5 mM $NH_4CH_3CO_2$ for MS analysis. Hydrolysis and derivatization of lipids extracts to fatty acid methyl esters (FAME) was performed on-line using trimethylsulfonium hydroxide [1]. FAMEs were analyzed with a gas chromatograph coupled to a mass spectrometer (GCMS – TQ8040; Shimadzu, Kyoto, Japan). The separation was carried out on a RTX-2330 capillary column (cyanopropyl stationary phase, 60 m x 0.25 mm, film thickness 0.20 µM; Restek, Bellefonte, PA, USA) and the electron ionization energy was set at 70 eV. Conditions for the analysis of FAMEs were as follows: carrier gas, He: column flow at 1 mL/min; 22:1 split ratio, injection volume 1 µL; injector temperature 240 °C; interface and ion source temperature 260 °C. GC oven temperature was maintained at 100 °C for 1 min, thermal gradient 100 °C to 140 °C at 10 °C / min, 140 °C to 175 °C at 6 °C / min, 175 °C to 200 °C at 10 °C / min and hold for 1 min, followed by 200 °C to 250 °C at 5 °C / min and hold for 4 mins. The data were acquired with Q3 scan mode from *m/z* 50 – 650. For data collection the MS spectra were recorded from 4.6 min to 28.33 mins. The data was processed in GCMSsolution software (Shimadzy, Kyoto, Japan). Fatty acids were identified based on retention time alignment with reference FAMEs from a mixture of standards (Food Industry FAME mix (37 components), Restek, Bellefonte,

PA USA). All samples were normalised to internal standard C19:0 and peak areas of relevant FAMEs was subtracted from negative control samples.

**Quantitative mass spectrometry of isobaric mass tag-labeled peptides**

LNCaP cells were treated with DMSO or Enzalutamide (10 µM) for 21 days as described above. Cell lysates from 3 independent experiments were prepared (6 samples) and peptides were labelled according to manufacturer's protocol (ThermoFisher, cat. 90064). Briefly, proteins were precipitated overnight using 6 volumes of acetone. Dried pellet was resuspended in 50 mM TEAB, and samples were digested in trypsin overnight. Each sample was labeled with a different TMT label reagent (isobaric mass tag) for 1 hour, after which 5% hydroxylamine was added to quench the reaction and equal amounts of each sample were combined. Following C18 clean-up, 2 ug of total protein was injected for analysis on ESI QExactive HCD mass spectrometer. Data were analyzed using Proteome Discoverer$^{TM}$ software (Thermo Fisher), and data integration of proteomics and transcriptomics data was carried out with MetaboAnalyst software (Mc Gill University).

**Statistical Analysis**

Statistical analysis were performed with Graphpad Prism 8.3 (Graphpad Software, San Diego, CA) and R Studio (RStudio, Boston, MA). Data reported and statistical tests are described in the figure legends.

**Supplementary Figure Legends**

**Figure S1**: **Longitudinal analysis of therapy-induced reprograming of metabolic networks in PCa xenograft model of CRPC progression and PCa patients pre- and post ADT**

(A) Gene set variation analysis (GSVA) of the transcriptome data obtained by microarray analysis of LNCaP tumor xenografts resected at the indicated therapy phases to monitor CRPC progression [2]: prior castration (intact, INT, n=10), regressing PSA post castration (REG, n=6), PSA nadir (NAD, n=10), rising PSA (recurring, REC, n=6), PSA exceeding intact (castrate-resistant, CR, n=6) [2]. The heatmap was generated with a hierarchical clustering algorithm using completed linkage and Euclidean distance measures and scaled by row z score (red=positive z score, blue=negative z score). The time points of tumor resection are indicated in the schematic. (B) Protein lysates of LNCaP cells treated for 0 and 21 days with Enz (10 µM) were trypsin digested, and peptides labelled with isobaric mass tag labels for quantitative proteomics analysis (n=3). Partial least squares (PLS) discriminant analysis shows the separate clustering of both treatments (top left panel). Volcano plot of differentially regulated proteins relative to day 0 (D0; FDR: fold change ≥2.0, p≤0.05). The 25 most significantly differentially expressed proteins are named (red label).

(H)

(C) Heatmap showing fold-changes of the top 50 most significantly dysregulated proteins relative to D0 was generated with a hierarchical clustering algorithm using completed linkage and Euclidean distance measures and scaled by row z score (red=positive z score, blue=negative z score).

(D) Integrated analysis of proteomics and transcriptomics data measured by microarray (Fig. 2) to identify significantly enriched metabolic pathways of the Kyoto Encyclopedia of Genes and Genomes (KEGG) was conducted with MetaboAnalyst 4.0 software [5].

(C) Indicated transcriptomics data sets from this study and study-relevant Gene Expression Omnibus data [3, 4] were analyzed by comparative signature scoring of lipid metabolism related gene sets using GSVA. Non-scaled bubble plots were created with the Morpheus webtool, with color indicating the direction of change in the GSVA score (red=increased scores/gene sets increase in overall expression, blue=decreased scores/gene sets decrease in overall expression; sample sources: gray=PCa patient, pink=LAPC9, yellow=LNCaP, orange=C4-2B; $ENZ^R$=Enz resistant cell lines, CRPC=castrate resistant tumor xenograft after castration, $1^{st}$ and $2^{nd}$=series of transplantation, Post=patient samples 22 weeks after androgen deprivation therapy (ADT), Pre=matching patient samples prior ADT). Genes of the Li et al 2018 signatures [4] are listed in Table S4.

**Figure S2: Functional and morphological characterization of therapy-induced changes**
LNCaP cells were treated for the indicated times with Enz (10 µM), and (A) AR mRNA expression was analyzed by qRT-PCR (left panel, n=3). (B) Maintained suppression of AR signaling is exemplified by the heatmap of microarray data showing classical androgen regulated genes that were differentially expressed at indicated time points of Enz treatment. For hierarchical clustering, an algorithm of completed linkage and Euclidean distance measures was used, and heatmap was scaled by row z score (red=positive z score, blue=negative z score). (C) Metabolic activity of LNCaP cells treated for the indicated times with Enz (10 µM) or androgen-depleted media (CSS) was analyzed as a measure of cellular ATP levels by CellTiter-Glo assay (top left panel, n=3, mean±SD), reductive power by PrestoBlue (top right panel, n=3, mean±SD) and mitochondrial membrane potential (MMP) by qSCI of MitoTracker Orange CMTMRos (bottom left panel, n>9000 cells, mean±SD, results are a representative of three independent experiments). Representative images are shown (bottom right panel, 40x magnification). (D) Proliferation of cells treated as in B was assessed based on cell confluence measured by phase contrast imaging every 2 hours for 96 hours (IncuCyte, n=3, mean±SD). Representative images (10x magnification) taken at 96 h of IncuCyte experiment demonstrate morphological difference between control (D0) and 21 days of treatment with Enzalutamide (Enz D21). (E) The percentage of cell death (left panel) was calculated based on the ratio of dead cells (propidium iodide-positive) relative to total cell count (Hoechst 33342-positive) by qSCI (n>9000 cells, mean±SD, results are a representative of three independent experiments). Autophagy (right panel) was assessed in LNCaP cells treated for 14 days with Enz (10 µM) by qSCI of CytoID-stained cells (right, n>9000 cells, mean±SD, results are a representative of three independent experiments). (F) Morphometric analysis based on qSCI of F-actin staining (cell mask) to measure cell area (left), cell perimeter (middle) and major axis length through cell body (right) (n>9000 cells, results are a representative of three independent experiments). (G) C4-2B (top panel) and DuCaP cells (middle panel) were cultured in growth media without (FBS) or with Enz (10 µM, FBS+ENZ D14) for 14 days including the final 24 h in the presence of vehicle control (DMSO) or the indicated compounds (paclitaxel 2.5 µM, docetaxel 2.5 µM, doxorubicin 2.5 µM, etoposide 5 µM, edelfosine 7.5 µM, TOFA 60 µM, RSL3 1.5 µM). LNCaP cells (bottom panel) were cultured in androgen-depleted growth media (CSS)for 14 days and co-treated as above. The percentage of dead cells was calculated based on qSCI of Hoechst 3342 (total cell count) and propidium iodide (dead cells) staining (n=3 wells/treatment with >4000 cells/well, representative result of 2 independent experiments). Statistical analysis for A-G: mean±SD, *p<0.05, **p<0.01, ***p<0.001, ****p<0.0001, One-way ANOVA followed by Dunnett's multiple comparisons test compared to D0/FBS).

**Figure S3**: **Lipidomics demonstrate therapy-induced lipid remodeling of all major lipid species**
LNCaP cells were treated for the indicated times with Enz (10 µM), and lipids were extracted and analyzed by LC/MS shotgun lipidomics (n=2 with 3 technical replicates each=6 data points). (A) Partial least squares discriminant analysis with MetaboAnalyst software [5] showed the separate clustering of the four time points of Enz treatment (FBS=D0 of Enz treatment, Enz D7, Enz D14 and Enz D21). LC/MS analysis of fold-changes of (B) sphingomyelines (SM), (C) phosphatidylcholines (PC), (D) phosphatidylethanolamine (PE), (E) phosphatidylserines (PS), (F) phosphatidylglycerols (PG), (G) cholesterol-esters (CE) and (H) triacylglycerols (TAGs) in indicated samples relative to FBS/D0 (n=2, mean±SD, Two-way ANOVA followed by Tukey's multiple comparisons test). (I) Heatmap of the top 50 dysregulated lipid species relative to D0 (identified by One-way ANOVA and Fisher's least significant difference method) was generated with a hierarchical clustering algorithm using completed linkage and Euclidean distance measures and scaled by row z score (red=positive z score, blue=negative z score; LPC=lyso-phosphatidylcholine).

**Figure S4**: **ATTs reduce glucose uptake but increase lipid transporter expression and cargo-nonselective mechanisms of nutrient uptake**
(A) Following 14 days of Enz treatment (10 µM), glucose uptake of LNCaP and C42B cells was measured by qSCI of NBD-2DG (n>9000 cells, mean±SD, results are a representative of 3 independent experiments). (B) Protein expression analysis of indicated lipid transporters and two lipogenesis enzymes (FASN and HMGCR) in PCa patient samples of normal gland (gray), localized primary tumor and (blue) and bone metastasis (red) as reported in the Iglesias-Gato proteome data set [6]. (C) LNCaP cells were treated for the indicated times with Enz (10 µM), and fatty acid content was analyzed by GC/MS FAME (n=2, mean±SD). (D) Fatty acid composition of fetal bovine serum (FBS) and charcoal-stripped FBS (CSS) used in these studies was analyzed by GC/MS FAME. Cholesterol and TAG concentrations are according to the supplier's certified analysis. (E) LNCaP cells were treated as described in (A), and macropinocytosis was measured by qSCI of 70 kDa dextran-TR (n>9000 cells, results are a representative of 2 independent experiments). (F) LNCaP cells were treated for the indicated times with Enz (10 µM), and the presence of tunneling nanotubes (TNTs) connecting cells was measured by counting 100 F-actin stained cells (top left panel). Representative images of Enz D21 stained with MitoTracker Orange CMTMRos (yellow) and Bodipy (green) are shown (40x magnification). Arrows highlight TNTs containing active mitochondria and lipid droplets. Fold-change of mRNA expression of genes regulating TNT maturation and maintenance based on microarray analysis are shown (bottom panel, n=3). Statistical analysis for A, D and E: mean±SD, *p<0.05, **p<0.01, ***p<0.001, ****p<0.0001, One-way ANOVA followed by Dunnett's multiple comparisons test compared to D0).

**Figure S5: Integrated fatty acid lipidomics and transcriptomics analysis and longitudinal analysis of LNCaP tumor xenograft model of CRPC progression highlight ATT-induced reprograming of PUFA metabolism**
(A) Integrated analyses (top panel) of dysregulated fatty acid species detected by GC/MS FAME (Fig. S4C) or (bottom panel) proteins detected by protein mass spectrometry (Fig. S1B) with transcriptomics data (Fig. 2) of LNCaP cells treated with vehicle control (FBS) or Enz (10 µM) for 21 days by MetaboAnalyst software 4.0 [5] (Mc Gill University) identified significantly enriched metabolic pathways of the Kyoto Encyclopedia of Genes and Genomes (KEGG) associated with PUFA metabolism. mRNA expression of indicated (B) fatty acid elongases and (C) fatty acid desaturases was measured by microarray in LNCaP tumor xenografts resected at the indicated therapy phases to monitor CRPC progression after

castration [2] (see Figure S1 for more details) and were calculated relative to expression in tumors from sham-castrated mice (intact, n=10): regressing PSA post castration (REG, n=6), PSA nadir (NAD, n=10), rising PSA (recurring, REC, n=6), PSA exceeding intact (castrate-resistant, CRPC, n=6) . The heatmaps were generated with a hierarchical clustering algorithm using completed linkage and Euclidean distance measures and scaled by row z score (red=positive z score, blue=negative z score). (D) C4-2B (top left panel), DuCaP (top right panel), 22Rv1 (middle panel) were cultured in growth media without (FBS) or with Enz (10 µM, FBS+ENZ D14) for 14 days including the final 24 h in the presence of DMSO (0) or the indicated concentrations (0.5-1.5 µM) of GPX4 inhibitor RSL3. Lipid peroxide scavenger Trolox (100 µM) and ferroptosis inhibitor Ferrostatin-1 (10 µM) were used as controls to demonstrate cell death through ferroptosis. LNCaP cells (bottom panel) were cultured in androgen-depleted growth media (CSS) for 14 days and co-treated as above. The percentage of dead cells was calculated based on qSCI of Hoechst 3342 (total cell count) and propidium iodide (dead cells) staining (n=3 wells/treatment with >4000 cells/well, representative result of 2 independent experiments). (E) LNCaP (left panel) were cultured in FBS or androgen-depleted growth media (CSS) and C4-2B cells (right panel) were cultured or in FBS or FBS+ENZ (10 µM) for the indicated times (D=days) including the final 24 h in the presence of DMSO (0) or RSL3 (1.25 µM). The percentage of dead cells was calculated as described in D (n=3 wells/treatment with >4000 cells/well, representative result of 2 independent experiments). (F) Gene ontology analysis (Panther, reactome version 65) of 334 significantly changed proteins of the proteomics data set (Fig. S1B) show significant enrichment of pathways related to oxidative stress and selenocysteine metabolism. The complete full list is shown in Table S3. (G) LNCaP cells seeded in a 96 well plate were co-treated with the indicated compounds (orlistat 10 µM, CAY10499 17.5 µM, SC26196 30 µM, TOFA 2.5 µM, fatostatin 5 µM, arachidonic acid (AA) 20 µM, trolox 100 µM) and cultured in FBS or androgen-depleted growth media (CSS). After 7 days of co-treatment, cells were treated with vehicle control (DMSO/FBS) or RSL3 (1.25 µM) to assess GPX4 dependence by measuring cell death by qSCI as described in D (n=3 wells/treatment with >4000 cells/well, representative result of 2 independent experiments). (H) Protein expression analysis of NPC1 and NPC2 in PCa patient samples of normal gland (gray), localized primary tumor and (blue) and bone metastasis (red) as reported in the Iglesias-Gato proteome data set [6].

### References


1. Butte, W., *Rapid method for the determination of fatty acid profiles from fats and oils using trimethylsulphonium hydroxide for transesterification.* Journal of Chromatography A, 1983. **261**: p. 142-145.
2. Locke, J.A., et al., *Arachidonic acid activation of intratumoral steroid synthesis during prostate cancer progression to castration resistance.* Prostate, 2010. **70**(3): p. 239-51.
3. Rajan, P., et al., *Next-generation Sequencing of Advanced Prostate Cancer Treated with Androgen-deprivation Therapy.* European Urology, 2014. **66**(1): p. 32-39.
4. Li, Q., et al., *Linking prostate cancer cell AR heterogeneity to distinct castration and enzalutamide responses.* Nature Communications, 2018. **9**(1): p. 3600.
5. Chong, J., et al., *MetaboAnalyst 4.0: towards more transparent and integrative metabolomics analysis.* Nucleic Acids Research, 2018. **46**(W1): p. W486-W494.
6. Iglesias-Gato, D., et al., *The proteome of prostate cancer bone metastasis reveals heterogeneity with prognostic implications.* Clinical Cancer Research, 2018.

**Fig S1A**

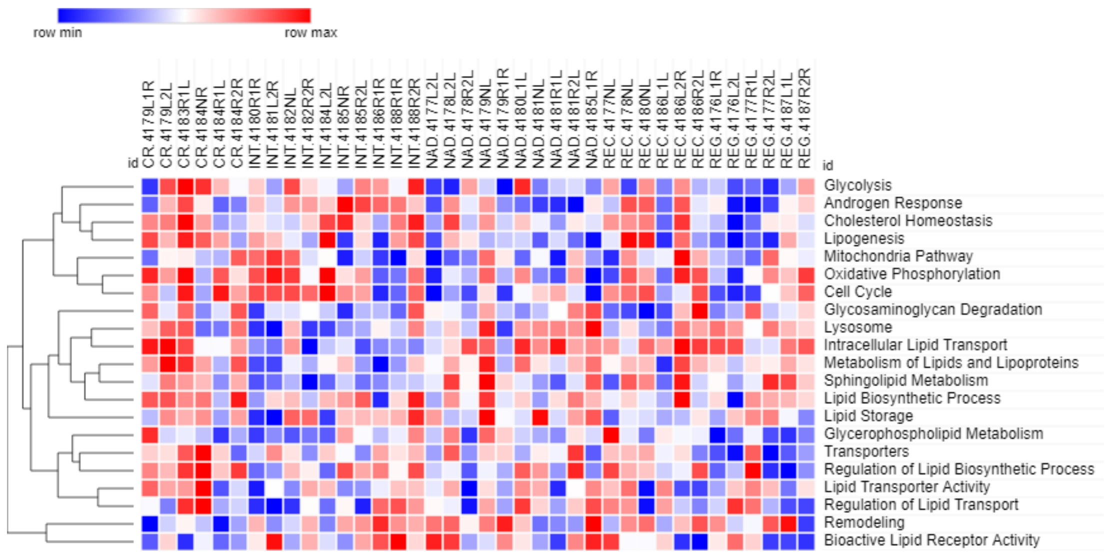

row min
row max
id
CR.4179L1R
CR.4179L2L
CR.4183R1L
CR.4184NR
CR.4184R1L
CR.4184R2R
INT.4180R1R
INT.4181L2R
INT.4182NL
INT.4182R2R
INT.4184L2L
INT.4185NR
INT.4185R2L
INT.4186R1R
INT.4188R1R
INT.4188R2R
NAD.4177L2L
NAD.4178L2L
NAD.4178R2L
NAD.4179NL
NAD.4179R1R
NAD.4180L1L
NAD.4181NL
NAD.4181R1L
NAD.4181R2L
NAD.4185L1R
REC.4177NL
REC.4178NL
REC.4180NL
REC.4186L1L
REC.4186L2R
REC.4186R2L
REG.4176L1R
REG.4176L2L
REG.4177R1L
REG.4177R2L
REG.4187L1L
REG.4187R2R
id
Glycolysis
Androgen Response
Cholesterol Homeostasis
Lipogenesis
Mitochondria Pathway
Oxidative Phosphorylation
Cell Cycle
Glycosaminoglycan Degradation
Lysosome
Intracellular Lipid Transport
Metabolism of Lipids and Lipoproteins
Sphingolipid Metabolism
Lipid Biosynthetic Process
Lipid Storage
Glycerophospholipid Metabolism
Transporters
Regulation of Lipid Biosynthetic Process
Lipid Transporter Activity
Regulation of Lipid Transport
Remodeling
Bioactive Lipid Receptor Activity


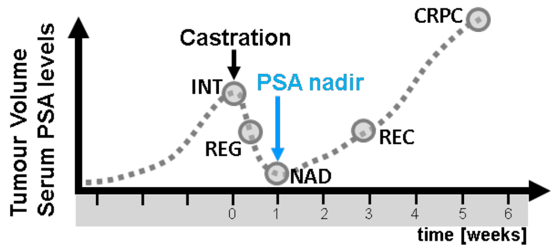

Tumour Volume
Serum PSA levels
Castration
CRPC
INT
PSA nadir
REG
REC
NAD
0
1
2
3
4
5
6
time [weeks]

**Fig S1B**

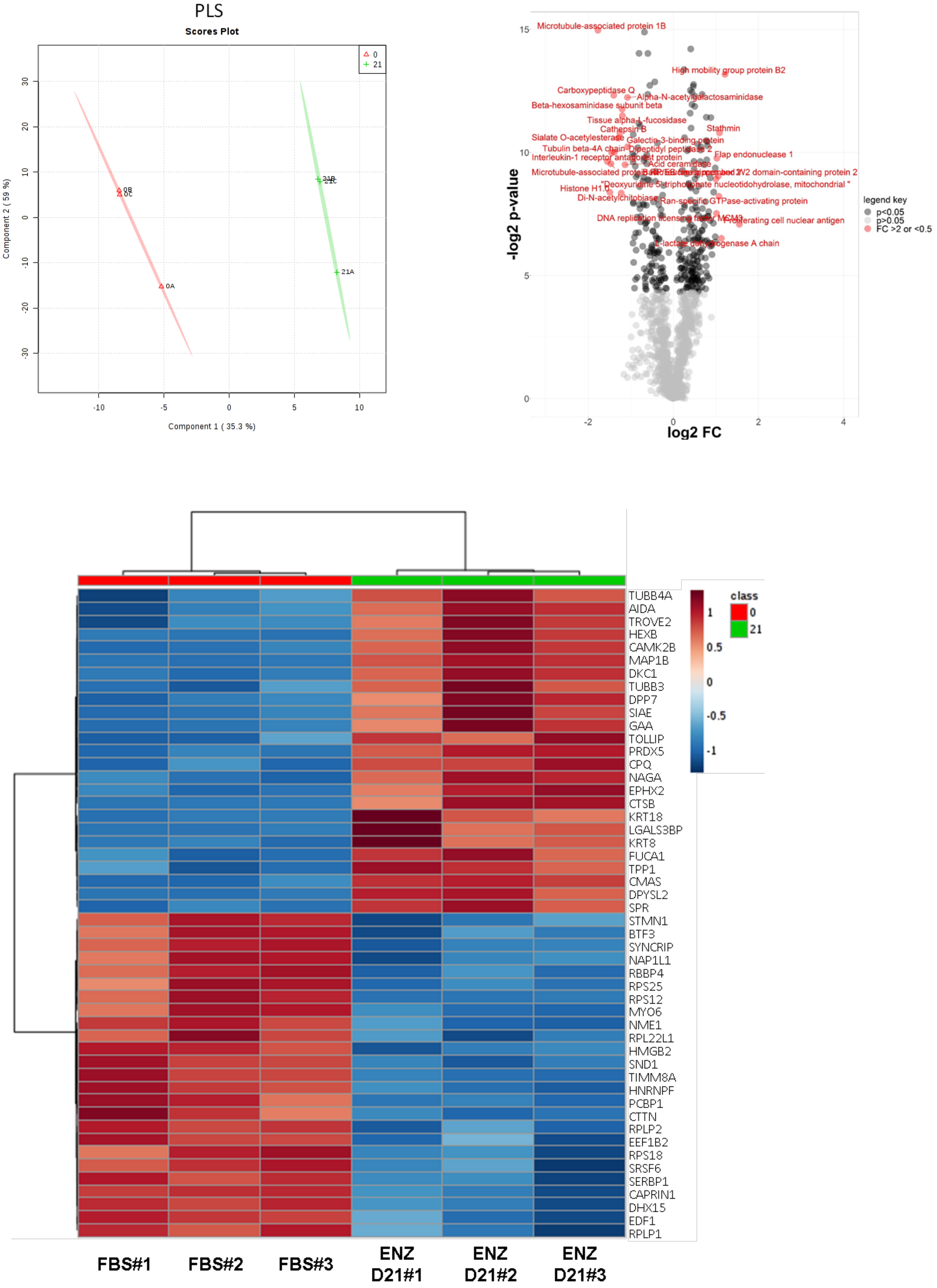

PLS
Scores Plot
0
21
0B
0C
0A
21B
21C
21A
Component 1 ( 35.3 %)
Component 2 ( 59 %)
-10
-5
0
5
10
30
20
10
0
-10
-20
-30
Microtubule-associated protein 1B
High mobility group protein B2
Carboxypeptidase Q
Alpha-N-acetylgalactosaminidase
Beta-hexosaminidase subunit beta
Tissue alpha-L-fucosidase
Cathepsin B
Stathmin
Sialate O-acetylesterase
Galectin-3-binding protein
Tubulin beta-4A chain
Dipeptidyl peptidase 2
Interleukin-1 receptor antagonist protein
Flap endonuclease 1
Acid ceramidase
Histone H1.0
Di-N-acetylchitobiase
Ran-specific GTPase-activating protein
Proliferating cell nuclear antigen
L-lactate dehydrogenase A chain
-log2 p-value
log2 FC
15
10
5
0
-2
0
2
4
legend key
p<0.05
p>0.05
FC >2 or <0.5
TUBB4A
AIDA
TROVE2
HEXB
CAMK2B
MAP1B
DKC1
TUBB3
DPP7
SIAE
GAA
TOLLIP
PRDX5
CPQ
NAGA
EPHX2
CTSB
KRT18
LGALS3BP
KRT8
FUCA1
TPP1
CMAS
DPYSL2
SPR
STMN1
BTF3
SYNCRIP
NAP1L1
RBBP4
RPS25
RPS12
MYO6
NME1
RPL22L1
HMGB2
SND1
TIMM8A
HNRNPF
PCBP1
CTTN
RPLP2
EEF1B2
RPS18
SRSF6
SERBP1
CAPRIN1
DHX15
EDF1
RPLP1
class
0
21
1
0.5
0
-0.5
-1
FBS#1
FBS#2
FBS#3
ENZ D21#1
ENZ D21#2
ENZ D21#3

Fig S1C

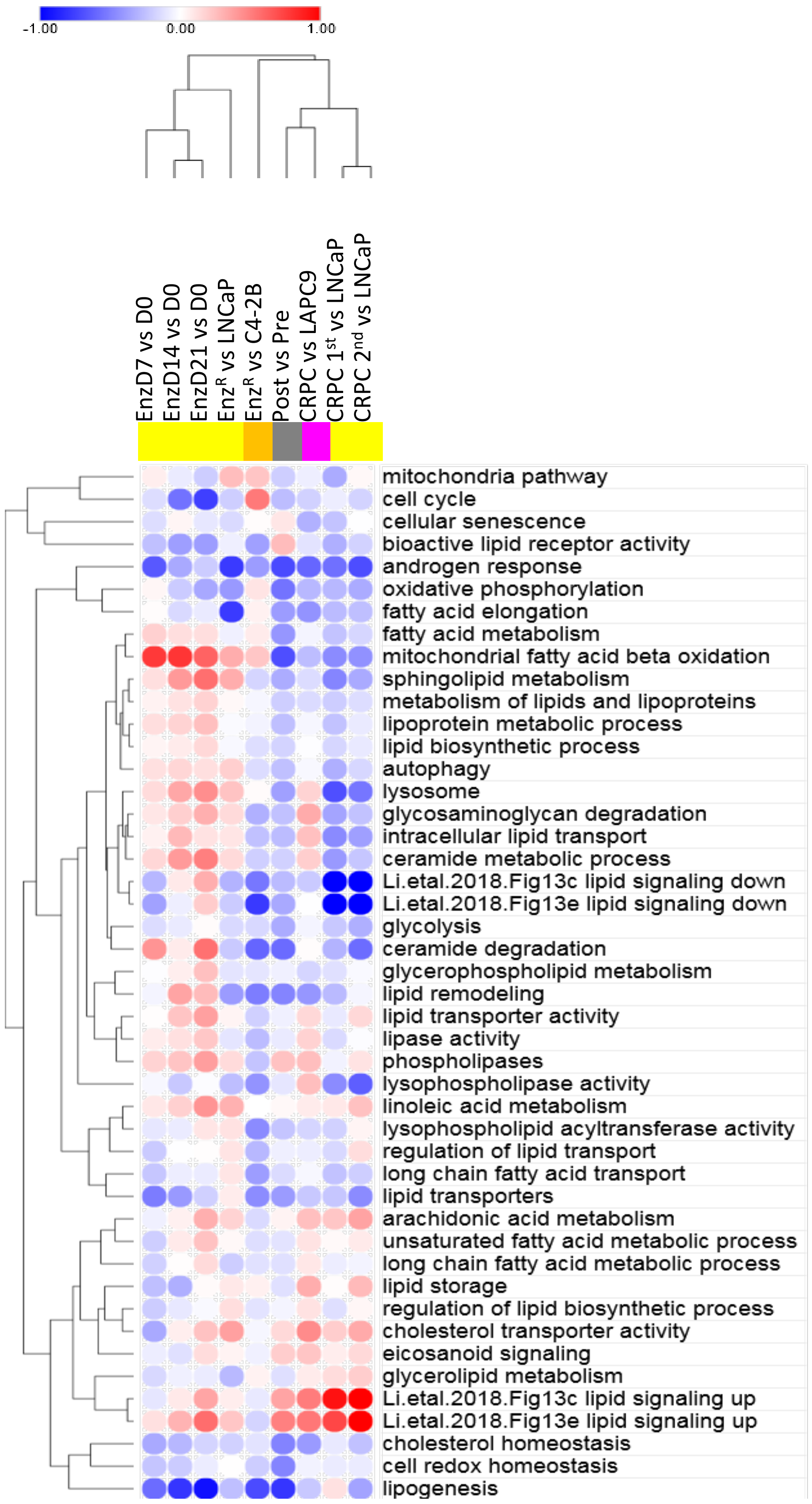
-1.00
0.00
1.00
EnzD7 vs D0
EnzD14 vs D0
EnzD21 vs D0
EnzR vs LNCaP
EnzR vs C4-2B
Post vs Pre
CRPC vs LAPC9
CRPC 1st vs LNCaP
CRPC 2nd vs LNCaP
mitochondria pathway
cell cycle
cellular senescence
bioactive lipid receptor activity
androgen response
oxidative phosphorylation
fatty acid elongation
fatty acid metabolism
mitochondrial fatty acid beta oxidation
sphingolipid metabolism
metabolism of lipids and lipoproteins
lipoprotein metabolic process
lipid biosynthetic process
autophagy
lysosome
glycosaminoglycan degradation
intracellular lipid transport
ceramide metabolic process
Li.etal.2018.Fig13c lipid signaling down
Li.etal.2018.Fig13e lipid signaling down
glycolysis
ceramide degradation
glycerophospholipid metabolism
lipid remodeling
lipid transporter activity
lipase activity
phospholipases
lysophospholipase activity
linoleic acid metabolism
lysophospholipid acyltransferase activity
regulation of lipid transport
long chain fatty acid transport
lipid transporters
arachidonic acid metabolism
unsaturated fatty acid metabolic process
long chain fatty acid metabolic process
lipid storage
regulation of lipid biosynthetic process
cholesterol transporter activity
eicosanoid signaling
glycerolipid metabolism
Li.etal.2018.Fig13c lipid signaling up
Li.etal.2018.Fig13e lipid signaling up
cholesterol homeostasis
cell redox homeostasis
lipogenesis

FigS 2A

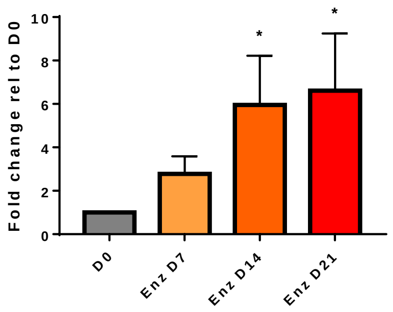


FigS 2B

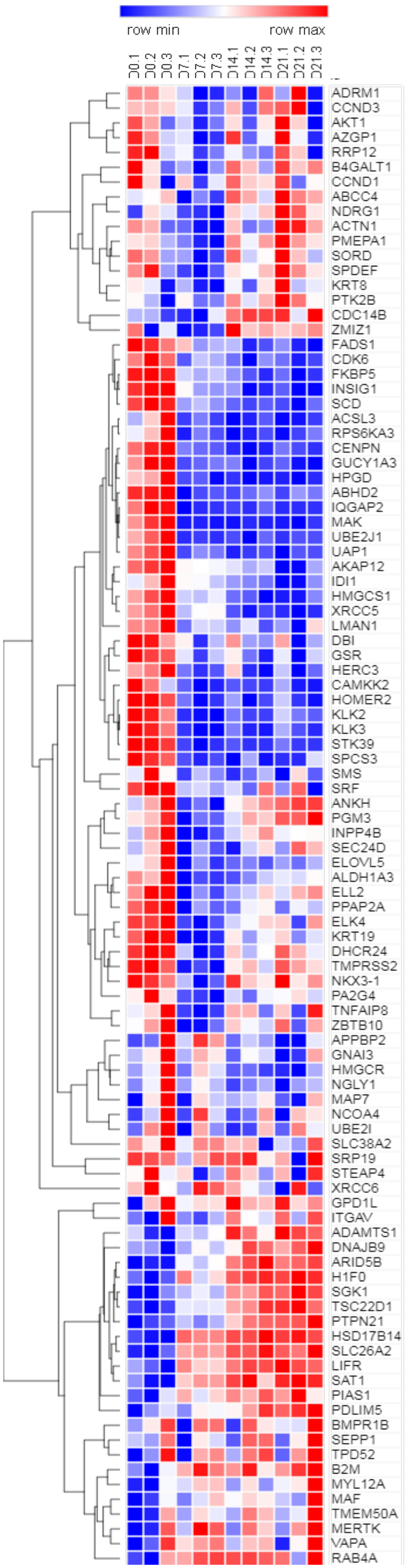

FigS 2C

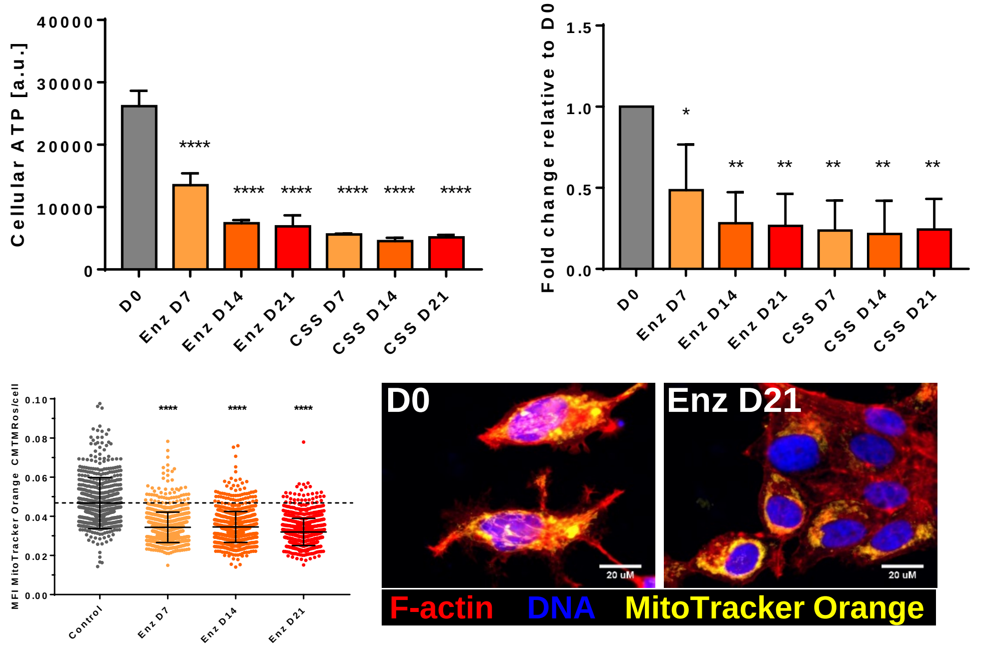


FigS 2D

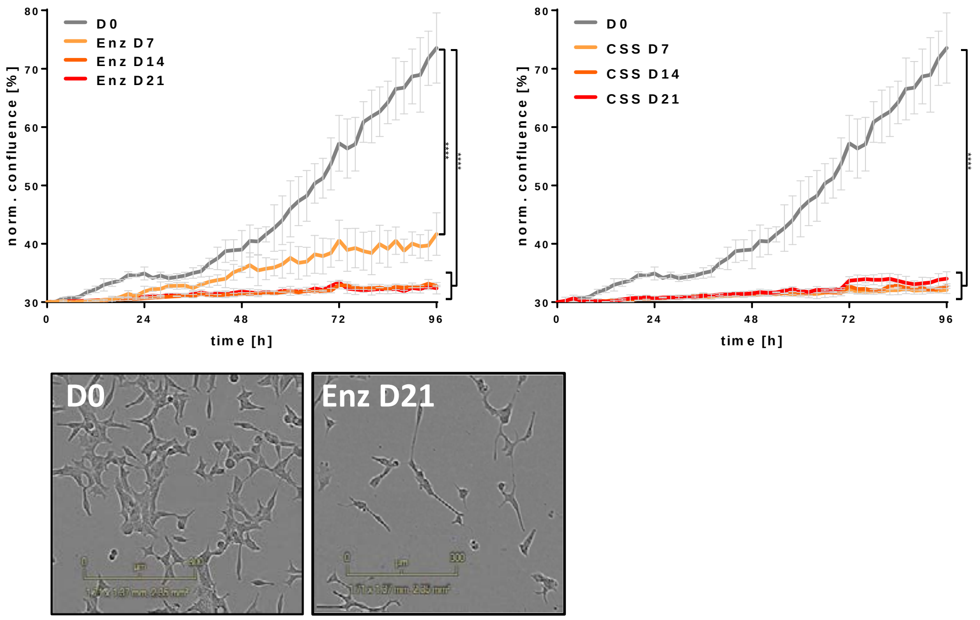


FigS 2E

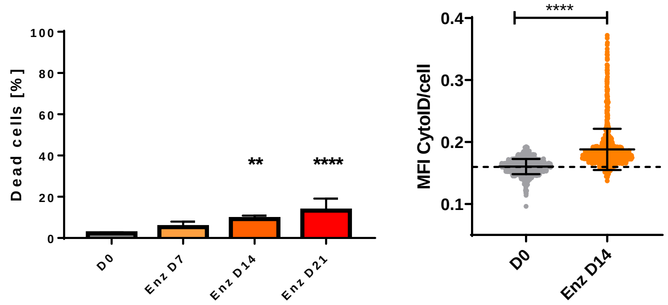


FigS 2F

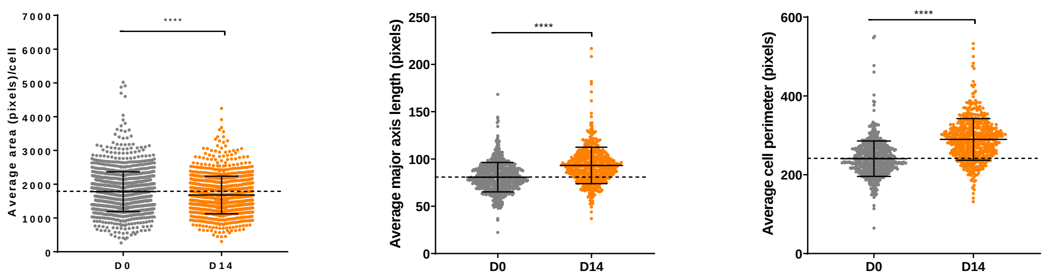

FigS 2G

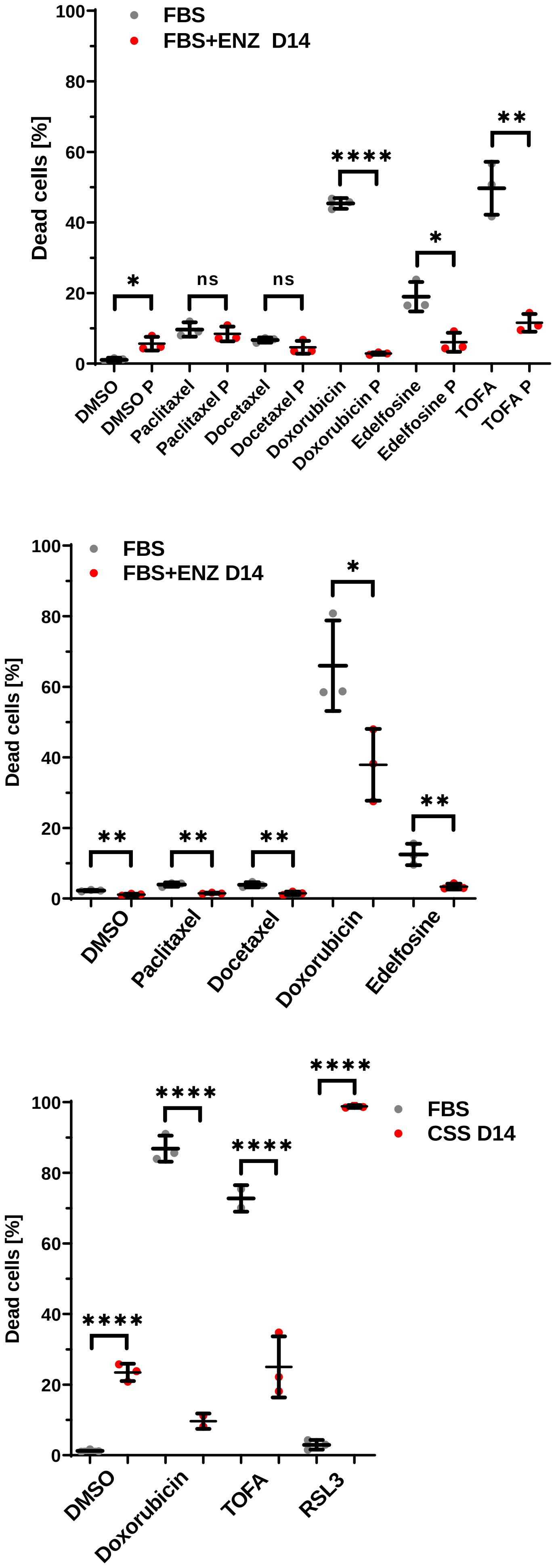

FBS
FBS+ENZ D14
Dead cells [%]
100
80
60
40
20
0
*
ns
ns
****
*
**
DMSO
DMSO P
Paclitaxel
Paclitaxel P
Docetaxel
Docetaxel P
Doxorubicin
Doxorubicin P
Edelfosine
Edelfosine P
TOFA
TOFA P
FBS
FBS+ENZ D14
Dead cells [%]
100
80
60
40
20
0
**
**
**
*
**
DMSO
Paclitaxel
Docetaxel
Doxorubicin
Edelfosine
FBS
CSS D14
Dead cells [%]
100
80
60
40
20
0
****
****
****
****
DMSO
Doxorubicin
TOFA
RSL3

Fig S3A

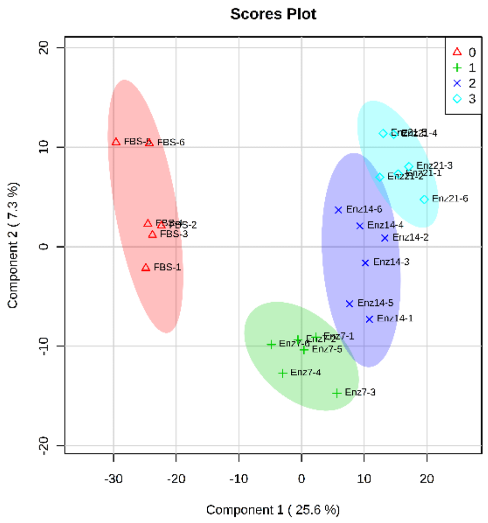


Fig S3B

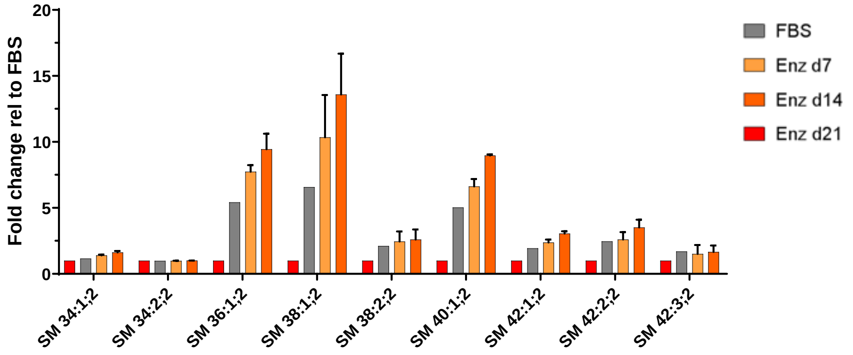


Fig S3C

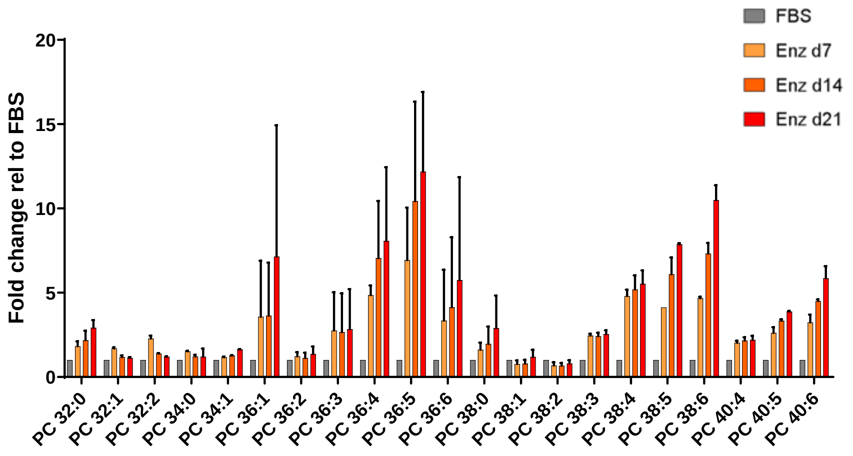


Fig S4D

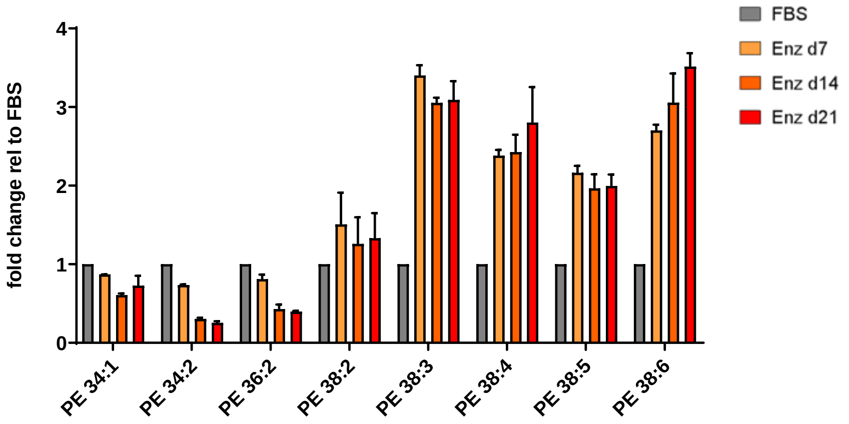

**Fig S3E**

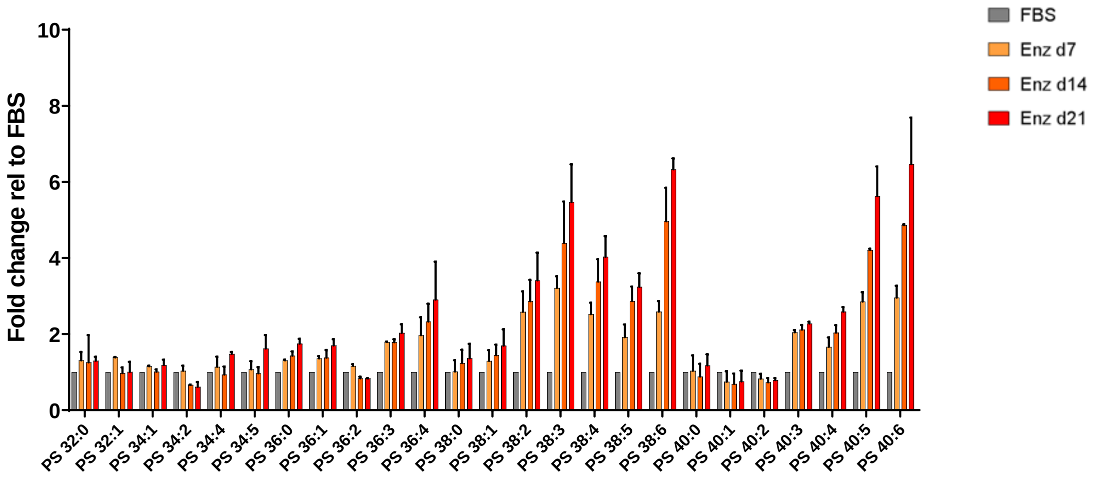


**Fig S3F**

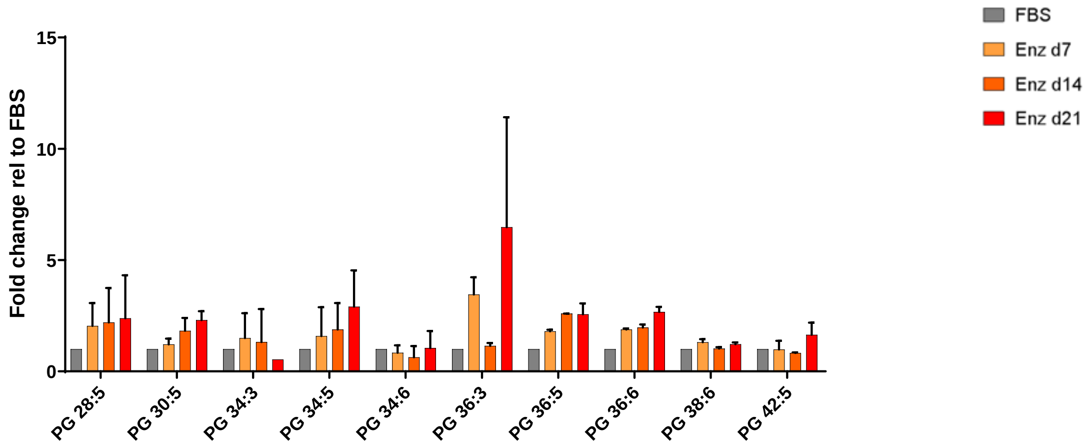


**Fig S3G**

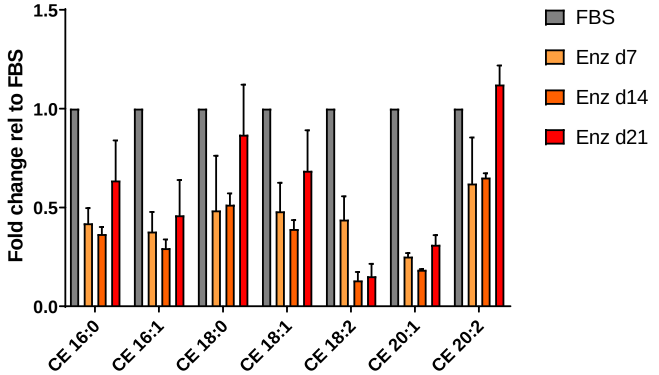


**Fig S3H**

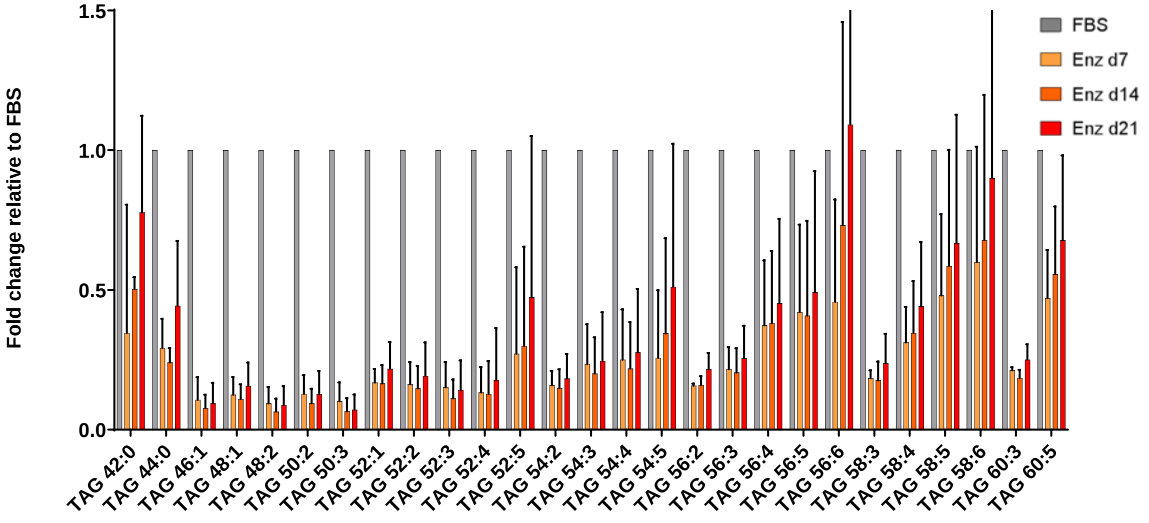

**Fig S3I**

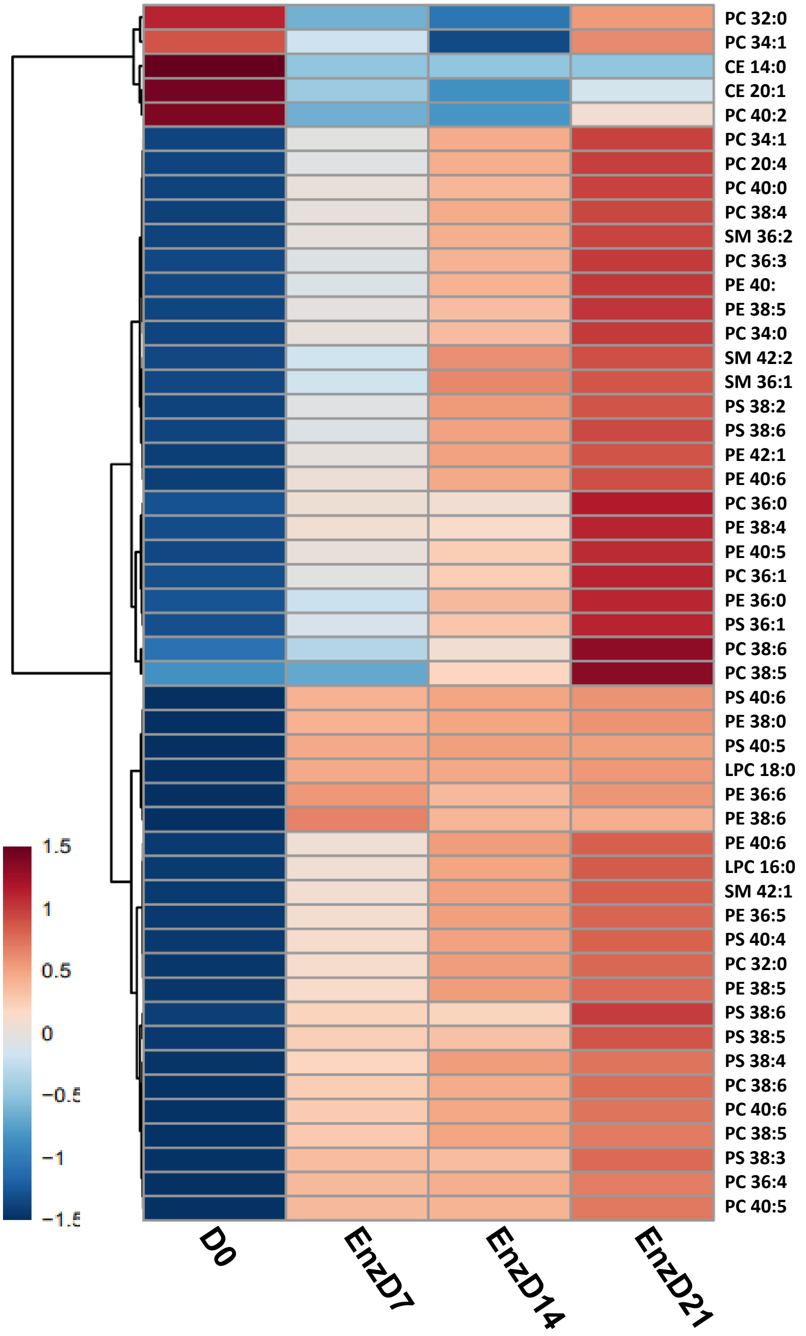
PC 32:0
PC 34:1
CE 14:0
CE 20:1
PC 40:2
PC 34:1
PC 20:4
PC 40:0
PC 38:4
SM 36:2
PC 36:3
PE 40:
PE 38:5
PC 34:0
SM 42:2
SM 36:1
PS 38:2
PS 38:6
PE 42:1
PE 40:6
PC 36:0
PE 38:4
PE 40:5
PC 36:1
PE 36:0
PS 36:1
PC 38:6
PC 38:5
PS 40:6
PE 38:0
PS 40:5
LPC 18:0
PE 36:6
PE 38:6
PE 40:6
LPC 16:0
SM 42:1
PE 36:5
PS 40:4
PC 32:0
PE 38:5
PS 38:6
PS 38:5
PS 38:4
PC 38:6
PC 40:6
PC 38:5
PS 38:3
PC 36:4
PC 40:5
1.5
1
0.5
0
−0.5
−1
−1.5
D0
EnzD7
EnzD14
EnzD21

FigS 4A

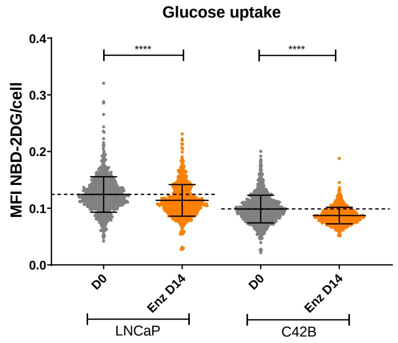


FigS 4B

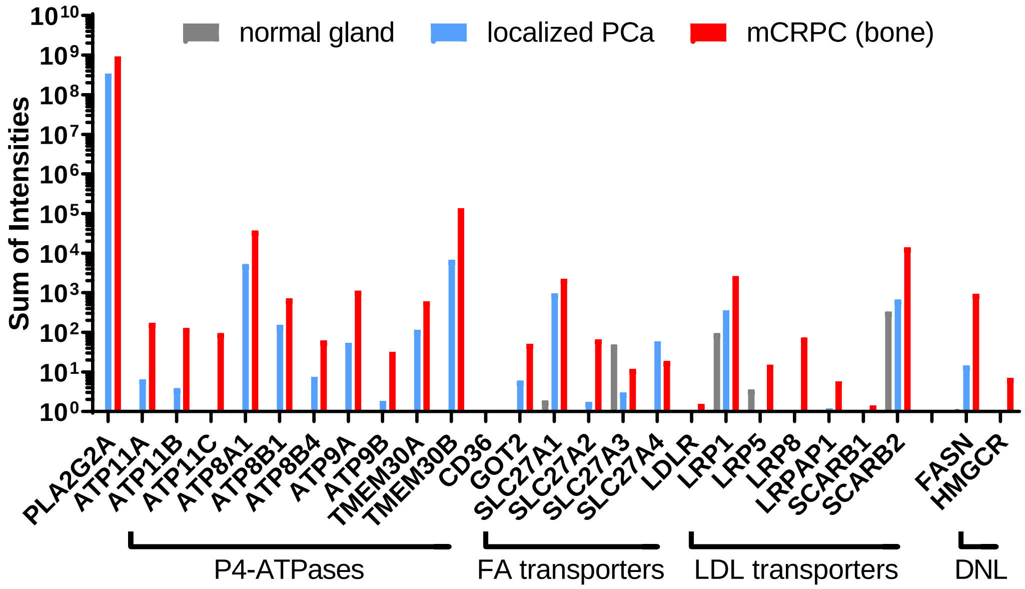


FigS 4C

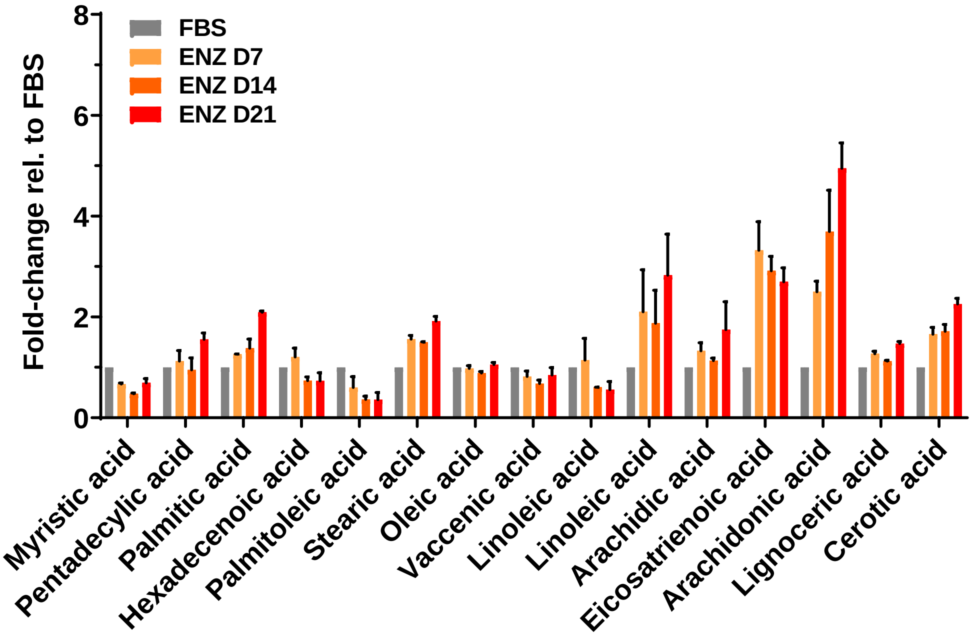

FigS 4D

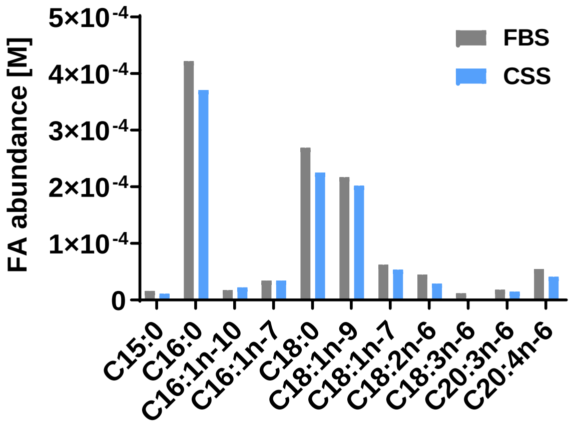


| | FBS | CSS |
|---|---|---|
| cholesterol | 34 mg/dL | 36 mg/dL |
| TAGs | 60 mg/dL | 66 mg/dL |

FigS 4E

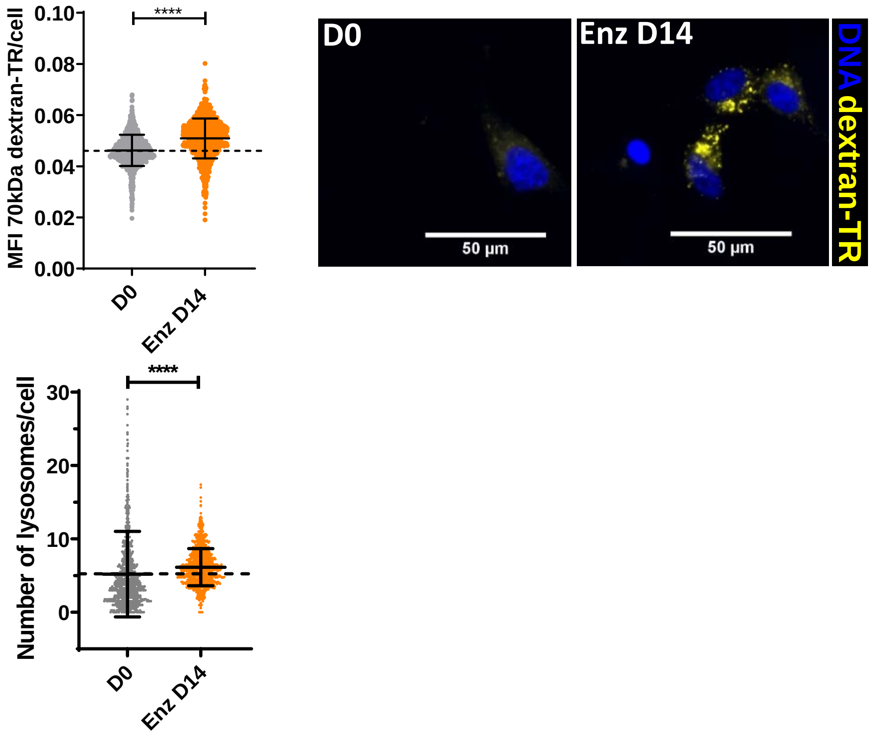


Fig S4F

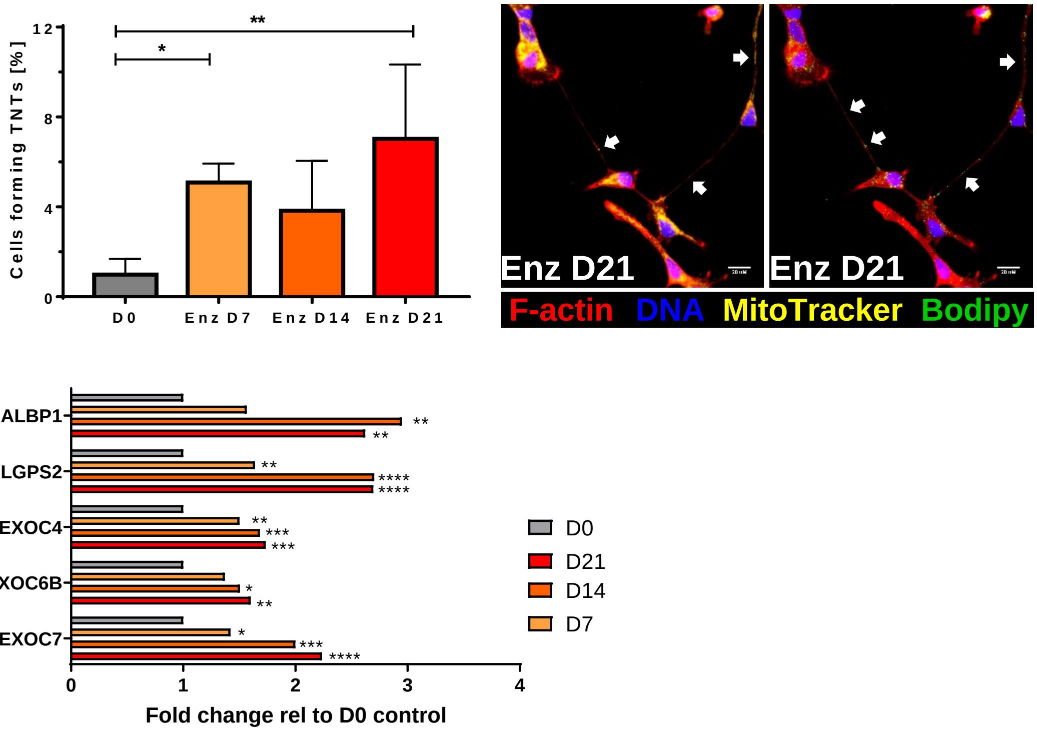

### Fig S5A

| KEGG pathway name | | Number of genes | P-value |
|---|---|---|---|
| Arachidonic acid metabolism | | 13 | 2.34e-10 |
| Linoleic acid metabolism | | 9 | 2.86e-10 |
| Retinol metabolism | | 8 | 0.000011 |
| Biosynthesis of unsaturated fatty acids | | 5 | 0.000025 |
| Drug Metabolism | | 7 | 0.0015 |
| Metabolism of xenobiotics | | 7 | 0.00253 |
| Fatty acid metabolism | | 4 | 0.0267 |
| Fatty acid elongation | | 3 | 0.0435 |

| KEGG pathway name | Number of proteins | P-value |
|---|---|---|
| **Biosynthesis of unsaturated fatty acids** | **5** | **3.99e-5** |
| **Glycosphingolipid biosynthesis** | **4** | **8.95e-4** |
| **Fatty acid metabolism** | **4** | **0.0062** |
| Fatty acid elongation | 1 | 0.0605 |

### Fig S5B

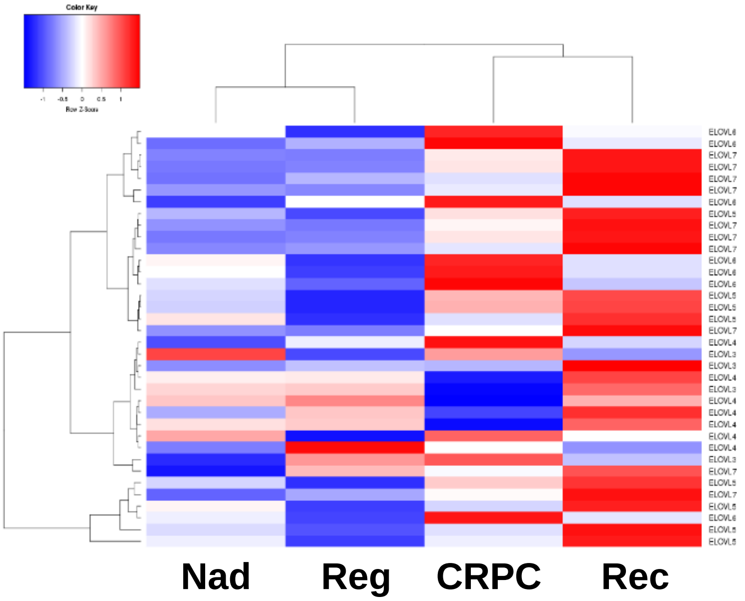


### Fig S5C

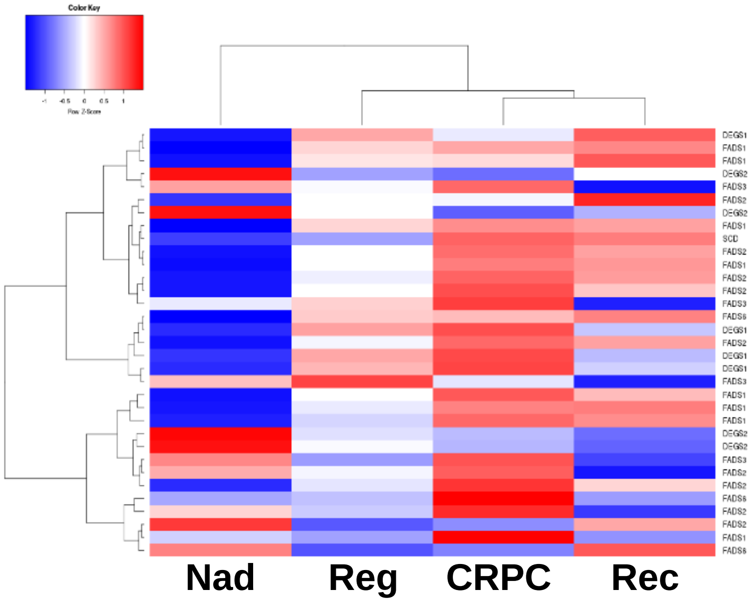

**Fig S5D**

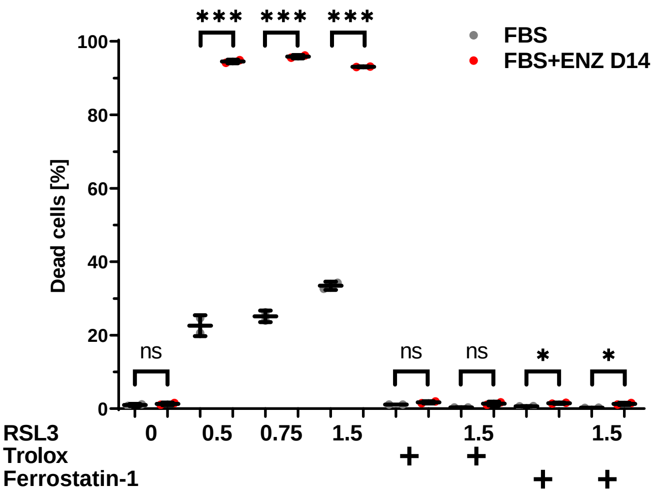
FBS
FBS+ENZ D14
Dead cells [%]
100
80
60
40
20
0
*** *** ***
ns ns ns * *
RSL3 0 0.5 0.75 1.5 1.5 1.5
Trolox + +
Ferrostatin-1 + +

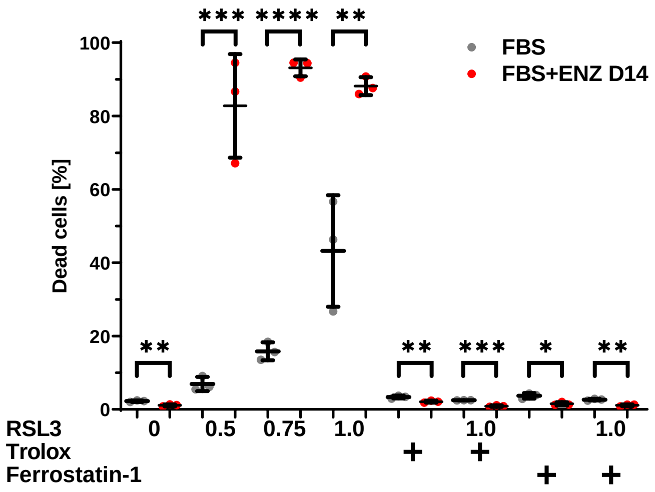
FBS
FBS+ENZ D14
Dead cells [%]
100
80
60
40
20
0
*** **** **
** ** *** * **
RSL3 0 0.5 0.75 1.0 1.0 1.0
Trolox + +
Ferrostatin-1 + +

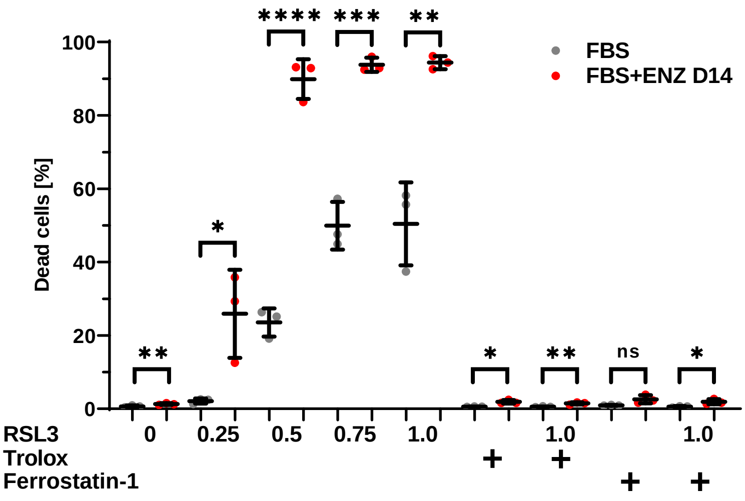
FBS
FBS+ENZ D14
Dead cells [%]
100
80
60
40
20
0
**** *** **
*
** * ** ns *
RSL3 0 0.25 0.5 0.75 1.0 1.0 1.0
Trolox + +
Ferrostatin-1 + +

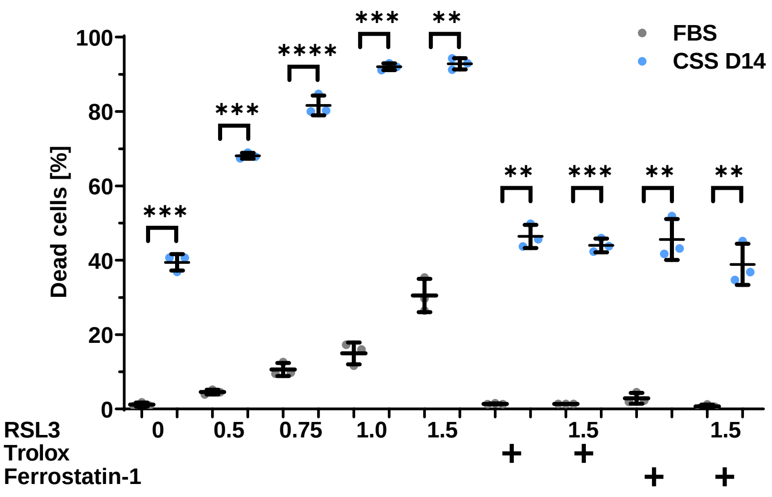
FBS
CSS D14
Dead cells [%]
100
80
60
40
20
0
*** **
****
***
***
** *** ** **
RSL3 0 0.5 0.75 1.0 1.5 1.5 1.5
Trolox + +
Ferrostatin-1 + +

**Fig S5E**

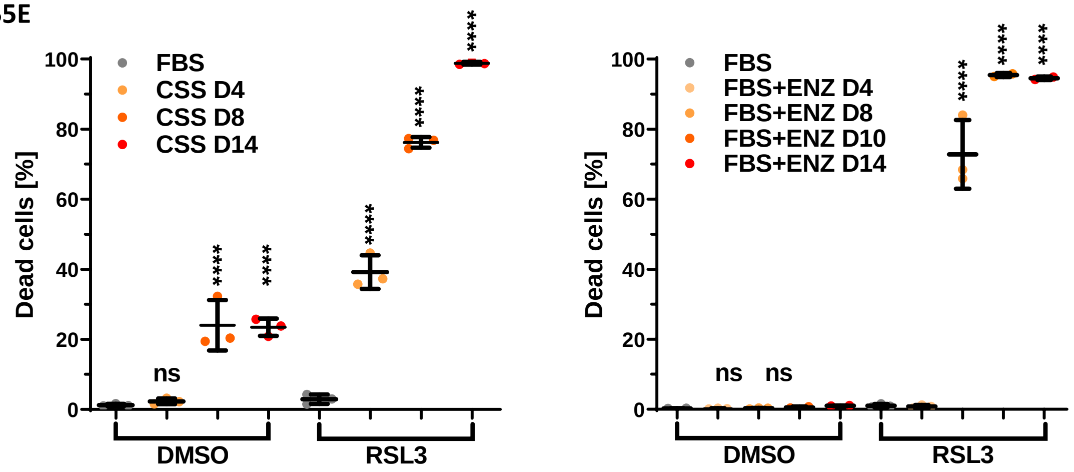


**Fig S5F**

| Reactome pathways | # reference list | # detected | # expected | +/- | Fold Enrichment | raw P value | FDR |
|---|---|---|---|---|---|---|---|
| Selenocysteine synthesis (R-HSA-2408557) | 93 | 51 | 1.48 | + | 34.47 | 1.48E-55 | 2.70E-53 |
| Selenoamino acid metabolism (R-HSA-2408522 | 116 | 51 | 1.85 | + | 27.64 | 1.01E-51 | 1.30E-49 |
| eNOS activation (R-HSA-203615) | 11 | 3 | 0.17 | + | 17.14 | 1.22E-03 | 2.47E-02 |
| Metabolism of nitric oxide (R-HSA-202131) | 15 | 3 | 0.24 | + | 12.57 | 2.61E-03 | 4.23E-02 |
| eNOS activation and regulation (R-HSA-20376 | 15 | 3 | 0.24 | + | 12.57 | 2.61E-03 | 4.17E-02 |
| Cellular responses to stress (R-HSA-2262752) | 393 | 19 | 6.25 | + | 3.04 | 3.01E-05 | 9.55E-04 |

**Fig S5G**

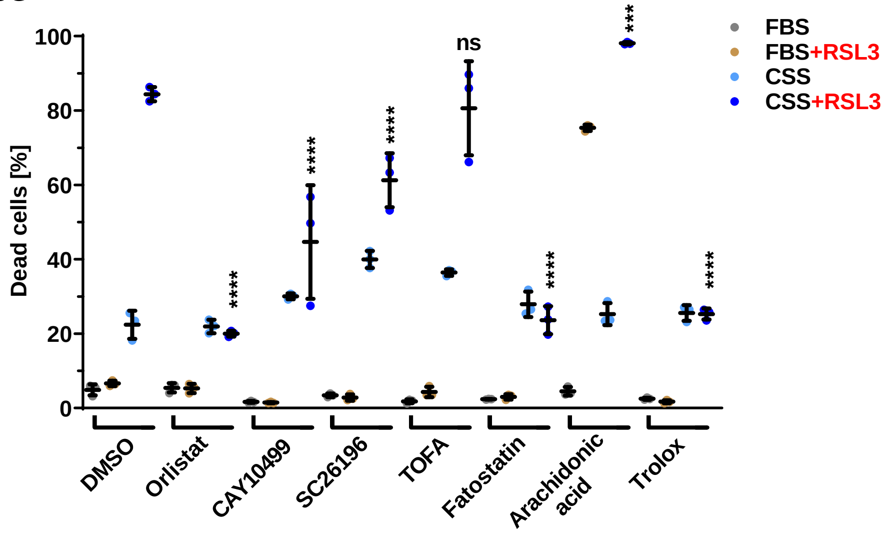


**Fig S5H**

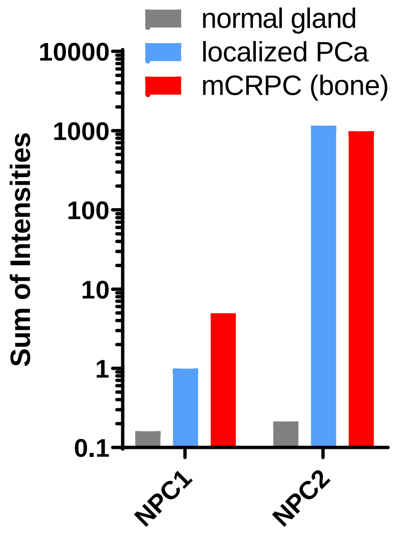
