## Supplementary material for "Therapy-induced lipid uptake and remodeling underpin ferroptosis hypersensitivity in prostate cancer": Table S1

|  | Fold change EnzD21 relative to D0 | p-val |
| --- | --- | --- |
| *Fatty acid elongation and desaturation*  ELOVL4  ELOVL5  ELOVL6  ELOVL7  DEGS2  SCD5  SCD1  FADS1  FADS2 | 2.07  -1.8829  -2.8738  -6.0917  10.8279  6.6161  -4.0693  -3.678  -1.5625 | 9.46E-05  0.000137  0.000226  1.15E-06  2.47E-07  4.92E-06  3.47E-05  5.58E-06  4.00E-05 |
| *Phospholipase activity*  **PLA2G2A**  PLD6  PLA2G16  PLA2G7  PLA2G12A  PLA2G4C  PLA2G5  PLA2G3 | 25.43  5.3686  3.4807  1.9485  1.9798  1.9337  -2.2416  -1.751 | 0.0122  2.82E-05  2.61E-05  0.000172  1.68E-05  0.01521  0.00026  5.73E-05 |
| *Phospholipid metabolism*  LPCAT4  LPCAT3  DGAT2  DGAT1  AGPAT2  AGPAT1  DAGLB  CHKA  PTDSS1 | 3.6328  2.4476  2.5756  -1.6133  2.2111  1.5432  1.5645  -2.2442  -1.4588 | 5.64E-06  6.57E-06  0.000155  0.001085  0.00063  0.006982  0.019658  1.93E-05  0.010547 |
| Lipogenesis  **FASN**  **ACACA**  ACLY  **ACAT1**  **ACAT2**  HMGCR  **HMGCS**  SREBF2 | -0.47289  -0.22538  -1.4748  1.411  -0.2259  2.026  -0.22  -2.2133 | 0.130905  0.195048  0.034312  0.140489  0.131584  0.044  0.448231  0.00022 |
| *Lipid transport*  ATP8A1  ATP8B2  ATP8B3  ATP9A  ATP10D  ATP11A  ATP11B  ATP11C  TMEM30A  LDLR  **SCARB1**  SLC27A1  SLC27A2  SLC27A3  SLC27A4  SLC27A5  FFAR2  SCARF1  FABP3  ABCG1 | 10.9597  2.2638  1.7422  2.4958  6.2332  5.64  2.0971  2.6338  1.979  2.5  1.1  5.518  2.637  -1.9027  2.686  -1.7862  -1.5286  1.5288  1.924  5.106 | 1.62E-07  8.01E-05  1.7422  3.41E-05  0.000172  0.003497  0.00026  0.002365  0.000284  0.0120  0.0001  0.0351  0.030  0.029178  0.064  6.07E-06  0.020756  0.024508  0.000273  0.000316 |
| *Lipid Storage*  PLIN  PPPDC1A  DGAT2  DGAT1 | 8.203  3.6787  2.5756  -1.6133 | 0.0001  2.81E-05  0.000155  0.001085 |
| *Stemness markers*  CD177  CD47  SALL4  CD200  CXCR4  CD99L2  ITGB1  BMI1 | 4.8309  3.1845  3.1315  3.0843  2.7323  2.4307  1.6362  1.608 | 6.77E-07  0.000131  7.64E-05  6.12E-07  0.010872  5.41E-06  0.005135  0.003883 |
| *Androgen regulation*  KLK3  TMPRSS2  FKBP5  PGC | -45.9853  -1.5006  -3.5932  -7.9768 | 3.23E-05  0.016943  0.000113  2.01E-6 |
