## Supplementary material for "Therapy-induced lipid uptake and remodeling underpin ferroptosis hypersensitivity in prostate cancer": Table S5

| Forward 5’ 3’ | Reverse 5’ 3’ |
| --- | --- |
| AR CTGGACACGACAACAACCAG  PSA AGTGCGAGAAGCATTCCCAAC  RPL32 GCACCAGTCAGACCGATATG  SLC27A1 AGGTGGTTCAGTACATCGGG  SLC27A2 GCACATTGCTGATTACCTACC  SLC27A3 TTTCTTCAGGAGGTGAACG  SLC27A4 GTCTTGGAGAAGGAACTGC  SLC27A5 CCCATTTCATCCGCATCC  LDLR GTTGACTCCAAACTTCACTCC  VLDLR TCAGTGTATCCCAGTGTCC  GOT2 GATCCGTCCCATGTATTCC  SCARB1 CCTTGTTTCTCTCCCATCC  PLIN ACCCCCCTGAAAAGATTGCTT  LRP8 GCAAATGAAGACAGTAAGATGG  SREBF2 TCCGCCTGTTCCGATGTAC  FASN CGCTCGGCATGGCTATCT  ACAT1 AATGAACAGAGGATCAACACC  ACAT2 GCCTTCCATTATGGGAATAGG  HMGCS TTCACCATGCCTGGATCACTT  HMGCR GGATGACTCGTGGCCCAGT  ACLY AAACTTGGTCTCGTTGGG | AR CAGATCAGGGGCGAAGTAGA  PSA CCAGCAAGATCACGCTTTTGTT  RPL32 ACTGGGCAGCATGTGCTTTG  SLC27A1 AGAACTCCCCGATTTGGC  SLC27A2 GATGACAGCAGGGTTAAAGC  SLC27A3 GGTGTAGAGCTGCATAAGG  SLC27A4 CAATAGCCGGGTCAAAGC  SLC27A5 GTTGTCCAGTACAAACAGAGG  LDLR GCTTCGTTGATGATATCTGTCC  VLDLR ATACAAAGTTCCTGGAGATGC  GOT2 CCATGACTTTCACTTCTTGC  SCARB1 TTCACAGAGCAGTTCATGG  PLIN GATGGGAACGCTGATGCTGTT  LRP8 GTTTCTCCAGATCAGGTATCC  SREBF2 TGCACATTCAGCCAGGTTCA  FASN CTCGTTGAAGAACGCATCCA  ACAT1 GTGCAATATTCAGCTTCTTTGC  ACAT2 CTATTGCAGCAGAGACAGC  HMGCS ATCTCAAGGGCAACAATTCCC  HMGCR TCGAGCCAGGCTTTCACTTC  ACLY TCGATCAGAAAGTTCTTGAGG |
